# Population genomic structure in Goodman’s mouse lemur reveals long-standing separation of Madagascar’s Central Highlands and eastern rainforests

**DOI:** 10.1101/2020.01.30.923300

**Authors:** George P. Tiley, Marina B. Blanco, José M. Ralison, Rodin M. Rasoloarison, Amanda R. Stahlke, Paul A. Hohenlohe, Anne D. Yoder

## Abstract

The Central Highland Plateau of Madagascar is largely composed of grassland savanna, interspersed with patches of closed-canopy forest. Conventional wisdom has it that these grasslands are anthropogenic in nature, having been created very recently via human agricultural practices. Yet, the ancient origins of the endemic grasses suggest that the extensive savannas are natural biomes, similar to others found around the globe. We use a phylogeographic approach to compare these two competing scenarios. By sampling multiple populations of Goodman’s mouse lemur (*Microcebus lehilahytsara*), a small-bodied nocturnal primate, we reconstruct the phylogeographic and demographic history of these “environmental metronomes” to estimate the time at which their populations diverged, and thus proximally, when their habitats would have become fragmented. We applied coalescent methods to RADseq data to infer phylogenetic relationships, population structure, and migration corridors among sampling sites. These analyses indicate that forest fragmentation occurred rapidly during a period of decreased precipitation near the last glacial maximum and would have affected both the Central Highlands and eastern forests. Though there is clear genomic structure separating the populations of the Central Highland from those of the eastern rainforests, there is also evidence of historical migration between them. Findings support the hypothesis that the Central Highland savanna predates human arrival, indicating that it is a natural landscape that has long impacted the population dynamics of Goodman’s mouse lemur, and by extension, other forest-dwelling organisms in Madagascar.

## Introduction

Madagascar is an iconic biodiversity hotspot with high levels of species diversity and endemism (Myers et al., 2000) with the majority of its biodiversity at extreme risk of extinction due to deforestation and forest fragmentation (Greene & Sussman, 1990; Harper, Steininger, Tucker, Juhn, & Hawkins, 2007; Vieilledent et al., 2018). It is widely accepted that the rapid rate of biodiversity loss is due largely to human activity and agricultural practices (Farris et al., 2017; Styger, Rakotondramasy, Pfeffer, Fernandes, & Bates, 2007). Even so, the timing of and degree to which humans have transformed and fragmented Madagascar’s closed-canopy forests has been a point of contention. Much of central Madagascar is now covered with savanna; traditionally thought to be strictly the product of fires from slash-and-burn agriculture and cattle farming (Perrier de la Bâthie, 1921; Humbert, 1927; Gade, 1996). The best available archeological evidence places humans in Madagascar between two thousand years ago (KYA) and ten KYA (Dewar & Wright, 1993; Dewar et al., 2013; Hansford et al., 2018; Mitchell, 2019). This timeline coincides with evidence of increased frequency of fire based on isotope records from stalagmites (Railsback et al., 2020) and charcoal records from sediment cores covering the late to mid Holocene (Burney, 1987a; Gasse & Van Campo, 1998; Virah-Sawmy, Willis, & Gillson, 2010). Pollen abundances from these same sediment cores also imply that the Central Highland vegetation transitioned from woody taxa to grasses during this timeframe, thus supporting the hypothesis that the Central Highland Savanna grasslands are the consequence of human activities. Confusingly, however, it appears that the transition to savanna likely began following Pleistocene glaciations (Burney, 1987b; Gass & Van Campo, 1998), long before the arrival of humans, and that similar shifts in plant composition were not limited to the Central Highlands (Burney, 1993; Virah-Sawmy, Willis, & Gillson 2009). This therefore implies that the Central Highland Savanna (CHS) is not strictly the product of human agricultural practices, and instead, may be the consequence of global climate fluctuations, perhaps facilitated by the natural processes of fire ecology (Bowman & Franklin, 2005).

In support of this second scenario, ecological and phylogenetic studies of Madagascar’s grasses suggest the that the CHS originated well before the Holocene, with levels of endemism comparable to the putatively natural grasslands of Africa (Bond, Silander Jr, Ranaivonasy, & Ratirarson, 2008; Vorontsova et al., 2016). And though few C_4_ grass lineages such as *Aristida* and *Loudetia* dominate the CHS (Koechlin, 1993; Kull, 2003; Solofondranohatra et al., 2018), diversity among endemic C_4_ grass clades would be more consistent with the global diversification of C_4_ grasses during the Miocene (e.g. Edwards & Smith, 2010) rather than as a consequence of a large number of independent migration events (Hackel et al., 2018). We are therefore left with a complex model of Pleistocene climate change, perhaps driving the origins of the CHS as a savanna, but overlaid with deforestation and fragmentation, likely associated with early human activity. Consequently, we hypothesize that present-day distributions of forest-dwelling species, as well as the genetic diversity and connectivity among their populations, should reflect *both* ancient climactic changes as well as recent human-meditated processes. It is our goal to examine phylogenomic patterns among populations of Goodman’s mouse lemur (*Microcebus lehilahytsara*) to disentangle the impacts of these two environmental scenarios.

The geographic distribution of *M. lehilahytsara* is uniquely suited for this purpose. Though it is found in eastern montane forests, it has largely been considered to be a highland specialist (Radespiel et al., 2012; Blanco et al., 2017). Indeed, it is the only mouse lemur species known to inhabit the isolated forest fragments in the CHS (Blanco et al., in review). The extended range of *M. lehilahytsara* may be due to its ability to enter a state of prolonged torpor (Blanco et al., 2017) thus reducing metabolic demands during the period of extended low temperatures in the CHS during the winter season. While prolonged torpor or hibernation is obligatory in *Cheirogaleus*, another member of the family Cheirogaleidae (e.g. Dausmann, Glos, Ganzhorn, & Heldmaier, 2005; Blanco et al., 2018), mouse lemurs vary in their expression of daily torpor or hibernation, with some species reported to display daily torpor (less than 24 hours) exclusively and unable to deposit fat stores to sustain prolonged torpor (e.g. Schmid, 2001; Blanco et al., 2018). Although losses in genetic diversity of endangered species putatively due to human action are observable (Olivieri, Sousa, Chikhi, & Radespiel, 2008; Andriaholinirina et al., 2014a; Andriaholinirina et al., 2014b; Baden et al., 2014), a better reconstruction of pre-human conditions can reveal if some populations were already vulnerable, especially given that some populations have been robust to recent habitat loss and fragmentation (Sgarlata et al., 2018).

A recent phylogeographic study of mouse lemurs that included a very limited representation of *M. lehilahytsara* from the CHS suggested that it diverged from its sister species *M. mittermeieri* in the eastern rainforests near 50 KYA (Yoder et al., 2016). Subsequently, the species status of *M. mittermeieri* has been called into question with compelling genetic evidence indicating that it should be synonymized with *M. lehilahytsara* (Poelstra et al., 2020). This suggests the intriguing possibility that rather than be a highland specialist, as was originally hypothesized, *M. lehilahytsara* may instead be a widely distributed generalist, capable of existing in a broad range of habitat types occurring at varying elevations. Thus, demographic studies of these mouse lemurs may illuminate pre-human conditions in Madagascar given that they can arguably be described as “environmental metronomes,” expanding and contracting their populations to match climatically driven pulses in forest distributions.

Here, we sampled populations of putative *M. lehilahytsara* from multiple sites in the CHS as well as from eastern forests. Using phylogeographic approaches and coalescent methods to analyze a large RADseq dataset, we explored patterns of genetic connectivity among populations. We also estimated divergence times and effective population size (*N*_*e*_) among and between them. Findings show that population structure formed well-before any human influence, with isolation of populations in the CHS and eastern forests occurring before the LGM. However, post-divergence gene flow among the populations suggests that migration was possible between the CHS and eastern forests, posing no more of a geographic barrier than major rivers. Moreover, population sizes were largely robust to past climate change, and have actually increased through the Holocene, even within a 1 Km^2^ forest patch.

## Methods

### Study sites and sampling

*Microcebus lehilahytsara* individuals were captured at five field sites. Three of the locations, from north to south, Riamalandy (S16° 17’ 06.0” E48° 48’ 54.0” 850 m), Ambatovy (S18° 49’ 30.15” E48° 19’ 06.48” 1100 m) and Tsinjoarivo (S19° 40’ 48.5” E47° 46’ 14.5” 1609 m) are mid-elevation rainforests, situated along the eastern escarpment. Forests are fragmented along the corridor; although, the distance between patches has been exacerbated by deforestation since the 1950s (Vieilledent et al. 2018). Annual rainfall is 1400 mm and daily temperatures vary between 14 and 23 °C at Riamalandy (Goodman, Raherilalao, & Wohlauser, 2018), while Tsinjoarivo and Ambatovy, receive 1700-2000 mm of rainfall with temperatures between 15 and 23 °C (Goodman et al. 2018). At their lowest, temperature can approach freezing conditions at Tsinjoarivo (Dausmann & Blanco, 2016). The other two locations, Ambohitantely and Ankafobe (S18° 06’ 23.1” E47° 11’ 13.2” 1472 m) are located within the Central Highlands. Their vegetational components are characteristic of moist evergreen forests, although more open and drier (Goodman et al., 2018). Annual rainfall is around 1460 mm, and daily temperature between 12.5 and 23 °C, though in winter temperatures can drop substantially, below 6 °C (Goodman et al., 2018). In addition to generally drier conditions at Ambohitantely and Ankafobe compared to Ambatovy and Tsinjoarivo, these sites experience greater seasonality of precipitation too (Supplementary Figure S1). Ambohitantely is a 5600 ha protected area 130 Km northwest of Antananarivo that incorporates natural forest, savanna, and exotic tree plantations, but is subject to annual fires and threated by forest loss. Samples were collected in 2004 (Riamalandy), 2006 (Ambohitantely), 2007 (Tsinjoarivo), between 2009 and 2015 (Ambatovy), and between 2015 and 2016 (Ankafobe). Mouse lemurs were captured using live Sherman traps and released back to the forest later the same day. For description of capture methods see Blanco et al. (2017).

### RADseq library preparation and sequencing

*Microcebus lehilahytsara* samples were gathered from existing collections and a combination of genomic DNA from available tissue and previous whole-genome amplifications were used in order to maximize sampling of these rare and vulnerable species. Library preparations for RAD sequencing followed Ali et al. (2016), using a single-digest protocol with the restriction enzyme SbfI. A median insert size of 380 bp was selected for paired-end sequencing at the Duke Sequencing Core on an Illumina Hi-Seq 4000 with an anticipated 15x coverage per individual.

### Assembly and orthology evaluation

Data were assembled using STACKS v2.0b (Catchen, Amores, Hohenlohe, Cresko, & Postlethwait, 2011; Catchen, Hohenlohe, Bassham, Amores, & Cresko, 2013) using *de novo* methods (Supplementary Methods). Final phased loci (we use locus or loci to refer to a contiguous RAD fragment while sites refer to individual base pairs throughout) output by stacks were filtered such that at least three of the five sample geographic locations were represented with a maximum of 50% missing data (Supplementary Methods). These filtering options were applied in order to maintain computational feasibility of downstream model-based hypothesis testing procedures in addition to not biasing our analyses due to uneven sampling from localities. Additionally, most RADseq-based studies rely solely on heuristic measures such as numbers of individuals and some level of heterozygosity for identifying orthologous clusters; we use a different approach for evaluating orthology that we argue is less sensitive to any clustering parameters. We used syntenic evidence from the *Microcebus murinus* reference genome, with BLASTN (Altschul et al.,1990; Camacho et al., 2009) search of assembled RAD loci, to identify clusters susceptible to paralogy as well as merge orthologous clusters that were over-split by STACKS (Supplementary Methods).

An important note regarding our sequence data and library preparation method is that we expect even depth of coverage for the forward read with the restriction enzyme cut site, but lowered coverage for the reverse read with the staggered cut. Therefore, while the longer contigs assembled with the paired-end data are useful for circumscribing orthologous loci, we are skeptical of our ability to consistently call heterozygous loci with high confidence beyond the first 151 bp of each locus; many of our downstream hypothesis tests assume we can accurately call these heterozygous sites. Therefore, for each sequence for each orthologous cluster, we extracted 145 bp, the first 151 bp minus the 6 bp sequenced overhang from the SbfI restriction enzyme, for all future analyses. These 145 bp fragments with heterozygous base calls underwent a final alignment with MUSCLE (Edgar, 2004) v3.8.1551 using default parameters.

### Phylogenetic analyses

We estimated a phylogenies of all *M. lehilahytsara* individuals or of sampling sites using four approaches: 1) concatenated ML analysis with RAxML (Stamatakis, 2014) v8.2.11, 2) topological invariants of quartets approximated with SVDquartets (Chifman & Kubatko, 2014) implemented through PAUP* (Swofford, 2003) v4.0a build 165, 3) a Bayesian implementation of the multispecies coalescent (MSC) model for SNP data (Bryant et al., 2012) through BEAST2 (Bouckaert et al., 2014) v2.4.8, 4) full-likelihood implementation of the MSC with joint gene tree estimation using BPP v4.0 (Flouri, Jiao, Rannala, & Yang, 2018). Applying these four different methods allows us to explore uncertainty in phylogeographic history due to model assumptions and the underlying data used for each method.

RAxML analyses were unpartitioned and used the GTR+Γ model of molecular evolution (Tavaré, 1986; Yang, 1994). We conducted 20 unconstrained ML searches and performed 100 bootstrap replicates. We also used the ML framework to perform topological hypothesis testing. Considering that samples from Ambohitantely and Ankafobe are well-supported clade across all analyses, our phylogenetic question reduces to a single quartet. We then tested three competing hypotheses regarding the evolutionary history of Riamalandy, Ambatovy, and Tsinjoarivo. Constrained optimizations of topologies and site likelihoods were performed with RAxML and we evaluated support among our three competing topologies with an approximately unbiased (AU; Shimodaira, 2004) test implemented in CONSEL (Shimodaira & Hasegawa, 2001) v0.20.

RADseq data were evaluated using multiple approaches for topological estimation from invariants. First, we used full-length RAD loci with SVDquartets, which is robust to violations of independence of loci (Chifman & Kubatko, 2014). Second, we analyzed only variable sites that represented at least four individuals. To further examine sensitivity of topology estimation with SVDquartets to our underlying data, we analyzed 10 datasets that only used biallelic SNPS, randomly sampling one SNP per locus. Analyses sampled all quartets and performed 100 bootstrap replicates. Distributions of bootstrap trees from SVDqaurtets and ML analyses were compared using unweighted Robinson-Foulds (RF) distances (Robinson & Foulds, 1981) computed with RAxML.

For SNAPP, we analyzed a single dataset using one biallelic SNP per RAD locus. To reduce the amount of computing time needed for each analysis, we selected only two individuals per sampling location. Two replicates were run to evaluate convergence. Mixing and effective sample sizes were analyzed with TRACER v1.7 (Rambaut et al., 2018), while potential scale reduction factors were calculated with the R package CODA (Plummer, Best, Cowles, & Vines, 2006). Bayes factor analyses were used to explore competing topological hypotheses, complementary to our AU tests in the ML framework. We used stepping-stone integration (Xie et al., 2011) to obtain marginal likelihoods for 20, 40, and 60 steps. It is difficult to predict how many steps are necessary for marginal likelihood estimation, so we used a range to observe little change in differences of marginal likelihoods with increasing steps. Because differences in marginal likelihoods were large, we presented natural log Bayes factors (Kass & Raferty, 1995) as opposed to computing model probabilities. Prior settings and details regarding MCMC options are in the Supplementary Methods (Supplementary Material).

Species tree analyses with BPP used fixed species delimitations. We used full-length RAD loci, considering at least two individuals for sampling location. Eight independent chains collected 2500 posterior samples, which were combined for a posterior of 20000 samples total. Robustness of species tree estimation to prior choice on common ancestor ages and population sizes was explored with intentionally inappropriate priors (Supplementary Material).

### Delimiting population structure and testing migration hypotheses

To elucidate migration patterns between *M. lehilahytsara* populations in the CHS and eastern forests, we developed competing migration hypotheses and tested models with Bayes factors. First, we analyzed biallelic SNPs from RAD loci with STRUCTURE (Pritchard, Stephens, & Donnelly, 2000) v. 2.3.4, using the admixture model and default priors while treating each site unlinked and uncorrelated. We created 20 datasets by sampling 10000 unlinked SNPS per replicate. For each replicate, we ran 20 independent MCMC chains for one to eight clusters assumed *a priori* (*k*) in order to calculate Δ*k* for estimating the most likely number of Hardy-Weinberg groups (Evanno, Regnaut, & Goudet, 2005). Each chain ran for one million generations after a 100000 generation burn-in.

Population structure was also evaluated with fixation index (*F*_*ST*_) statistics. Pairwise *F*_*ST*_ was calculated between populations as described by Weir & Cockerham (1984) for SNPS with less than 15% missing data in ARLEQUIN (Excoffier & Lischer, 2010) v. 3.5.2.2. Pairwise *F*_*ST*_ were used to perform an AMOVA analysis (Excoffier, Smouse, & Quattro, 1992) for Central Highlands (Ambohitantely and Ankafobe) versus eastern (Riamalandy, Ambatovy, and Tsinjoarivo) *M. lehilahytsara*. A Mantel test was used to test the presence of isolation-by-distance using 9999 permutations. Physical distances between populations were calculated with the Geographic Distance Matrix calculator v 1.2.3 (http://biodiversityinformatics.amnh.org/open_source/gdmg; Last Accessed 2 January 2020).

While STRUCTURE analyses can estimate the posterior probabilities of admixture among Hardy-Weinberg groups, this is not a hypothesis test. As an independent evaluation of gene flow between *M. lehilahytsara* populations, we computed Bayes factors for isolation-with-migration models using MIGRATE v4.2.14 (Beerli & Palczewski, 2010). Notably, we did not use the default uniform prior for nucleotide diversity (θ) estimates. Instead, we used independent effective population size (*N*_*e*_) estimates from three mouse lemur species prior to forest fragmentation (Olivieri et al., 2008) to develop a prior distribution for θ (Supplementary Material). Two sets of migration hypothesis testing were conducted on full-length RAD loci. First, we used all individuals binned into either a CHS or eastern population, despite recognizable population substructure. We were limited in the number of loci computationally feasible for obtaining marginal likelihood estimates, and thus analyzed 100 data sets of 100 random RAD loci; missing data was allowed as long as at least two individuals per sampling site were present. The second analysis attempts to differentiate between east-to-west migration and west-to-east migration among Ambatovy, Ambohitantely, and Ankafobe, and does not allow for missing data. The three-population analyses used four diploid individuals per population and compared isolation-with-migration models to those that also included divergence. Posterior estimates of θ was rescaled *N*_*e*_ to using a mutation rate of 1.64 × 10^−8^, which is based on a direct germline estimate for mouse lemurs (Campbell et al., 2020).

### Estimating divergence time of eastern and Central Highland mouse lemurs

We used the multispecies coalescent (MSC) model for divergence time estimation among *M. lehilahytsara* populations with BPP (Rannala & Yang, 2003; Flouri et al., 2018) v4.0. However, estimating divergence times and population sizes efficiently requires a fixed and rooted topology. We used the topology recovered in the majority of analyses here, and used several lines of evidence to support rooting with Riamalandy (Supplementary Material), including a phylogeny of *M. lehilahytsara* individuals and outgroups using previously sequenced mitochondrial data (Table S2). A partitioned ML analysis and bootstrapping was conducted with PAUP* v4.0a build 163 (Swofford, 2003). Phased haploid sequences were used for BPP and at least two individuals per population were required for each locus. We ran eight chains of 2500 samples and evaluated convergence. Final MSC parameter estimates were based on combined chains. Coalescent branch lengths were rescaled to absolute time using a direct germline mutation rate estimate in mouse lemurs (1.64 × 10^−8^; Campbell et al., 2020) and generation time estimate (3.75 years/generation) based on wild (Radespiel et al., 2019) and captive (Zohdy et al., 2014) populations. Robustness of divergence times to prior choice was also evaluated (Supplementary Material).

### Characterizing population size changes during Pleistocene glaciations

We estimated change in population size over time with extended Bayesian skyline plots (Heled & Drummond 2008) with BEAST2 (Bouckaert et al., 2014). Similar analyses have also been successful in discovering king penguin population increases that coincide with Holocene warming (Trucchi et al., 2014). For all sampling locations with four or more individuals, we sampled 100 random RAD loci eight times, with no missing data. Sequences were treated as phased and haploid. Each sampled dataset was run twice to check convergence. Large posteriors were required for agreement on the number of change points and population size estimates between replicates; each chain ran for 500 million generations after a 100 million generation burnin while sampling every 10000 generations. Operators for increasing the sampling of change points followed recommendations by Trucchi et al. (2014). We used linear piecewise reconstructions of population size changes.

## Results

### Circumscribing orthologous RAD loci

All individuals included in our analyses had an average depth of coverage greater than 10x per locus assembled with STACKS. In total, 92,889 loci were recovered across all individuals with an average of 49,201 loci per individual (Supplementary Table S1). On average, 31% of loci were shared between pairs of individual (Figure 1). The number of loci shared between individuals are likely not due to phylogenetic signal, as there was only a slight negative correlation between evolutionary distance and the proportion of shared loci (Pearson’s *r* = −0.19; *p* = 0.001). Rather, this pattern was driven, at least in part, by the number of reads, given a strong positive correlation between the number of loci and the number of reads per individual (Pearson’s *r* = 0.86; *p* = 3.54e-08). Although 92,889 RAD loci were recovered, only 54,018 loci contained at least one biallelic SNP; other loci were completely invariable or consisted of only singletons.

**Figure 1.**
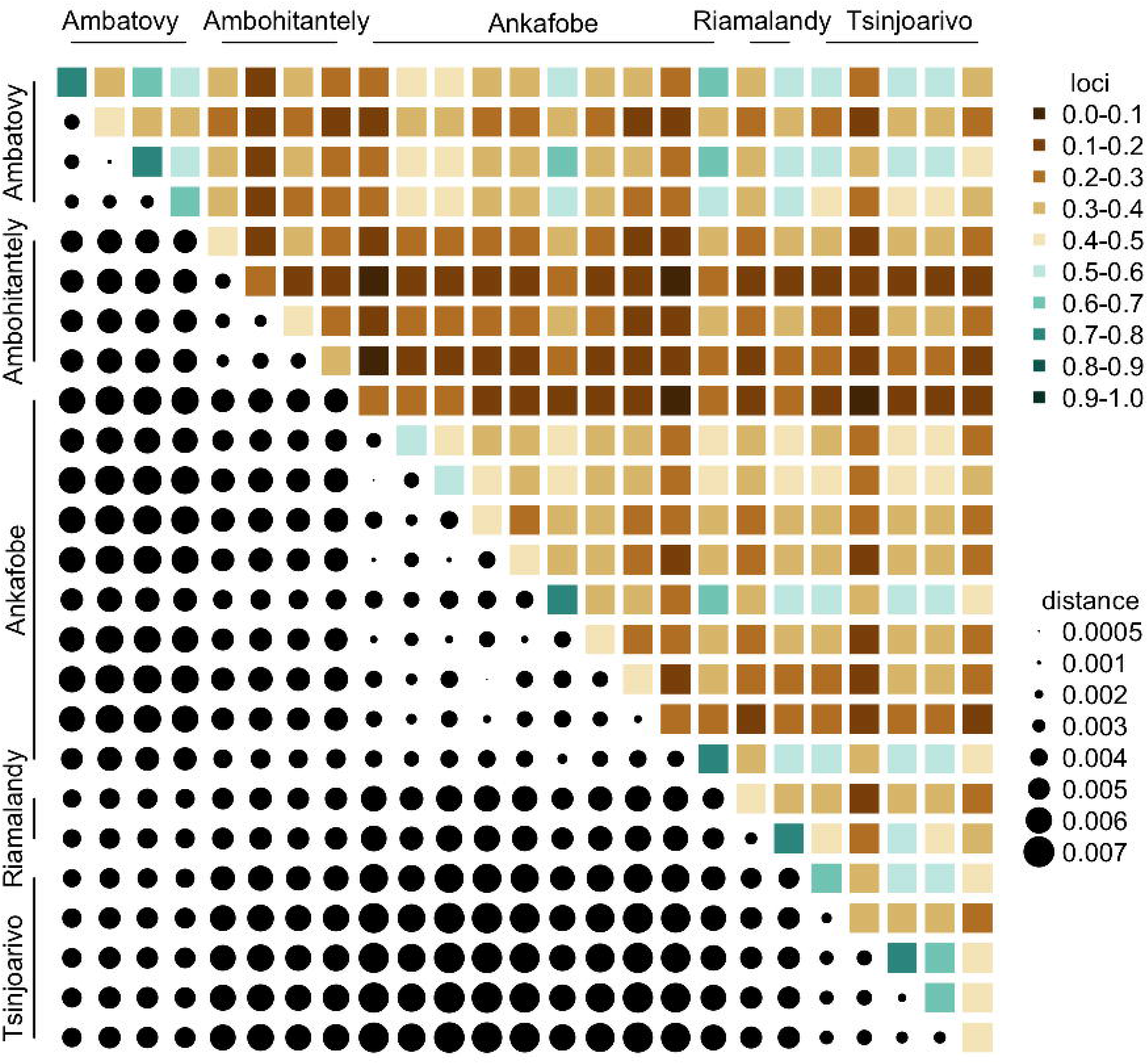
Recovery of RAD loci among *Microcebus lehilahytsara*. The diagonal represents the proportion of total loci recovered for each sample while the upper triangular represents the proportion of total loci shared between two individuals. The lower triangular represents the evolutionary distance between two individuals, which is the patristic distance from the concatenated ML analysis.

### Support for Central Highland and eastern Microcebus lehilahytsara

All phylogenetic analyses recovered a well-supported split between *M. lehilahytsara* from Central Highland (Ambohitantely and Ankafobe) and eastern forest (Riamalandy, Ambatovy, and Tsinjoarivo) localities (Figure 2; Supplementary Figures S2-S8). But the relationships among eastern sampling locations was contentious across analyses. The ML, SNAPP, and BPP analyses supported Ambatovy as sister to Tsinjoarivo (H_TA_; Supplementary Figures S2-S5), although posterior probabilities from SNAPP revealed uncertainty in the bipartition in question. SVDquartets strongly conflicted the predominant H_TA_ topology, favoring Riamalandy sister to Tsinjoarivo (H_RT_) in all bootstrap replicates when analyzing all variable sites (Supplementary Figure S6) or complete RAD loci (Supplementary Figure S7). However, the H_RT_ topology, even with perfect bootstrap support, is sensitive to the underlying data. Analyses of biallelic SNPs alone with SVDquartets recovered a third topology (H_RA_); also with high bootstrap support at bipartitions of sampling locations (Supplementary Figure S8). In one out of ten resampled datasets, when treating all individuals as independent lineages, Ambatovy was not recovered as a monophyletic clade. Topological hypothesis testing differentiated among the three recovered phylogenetic hypotheses: H_TA_, H_RT_, and H_RA_.

**Figure 2.**
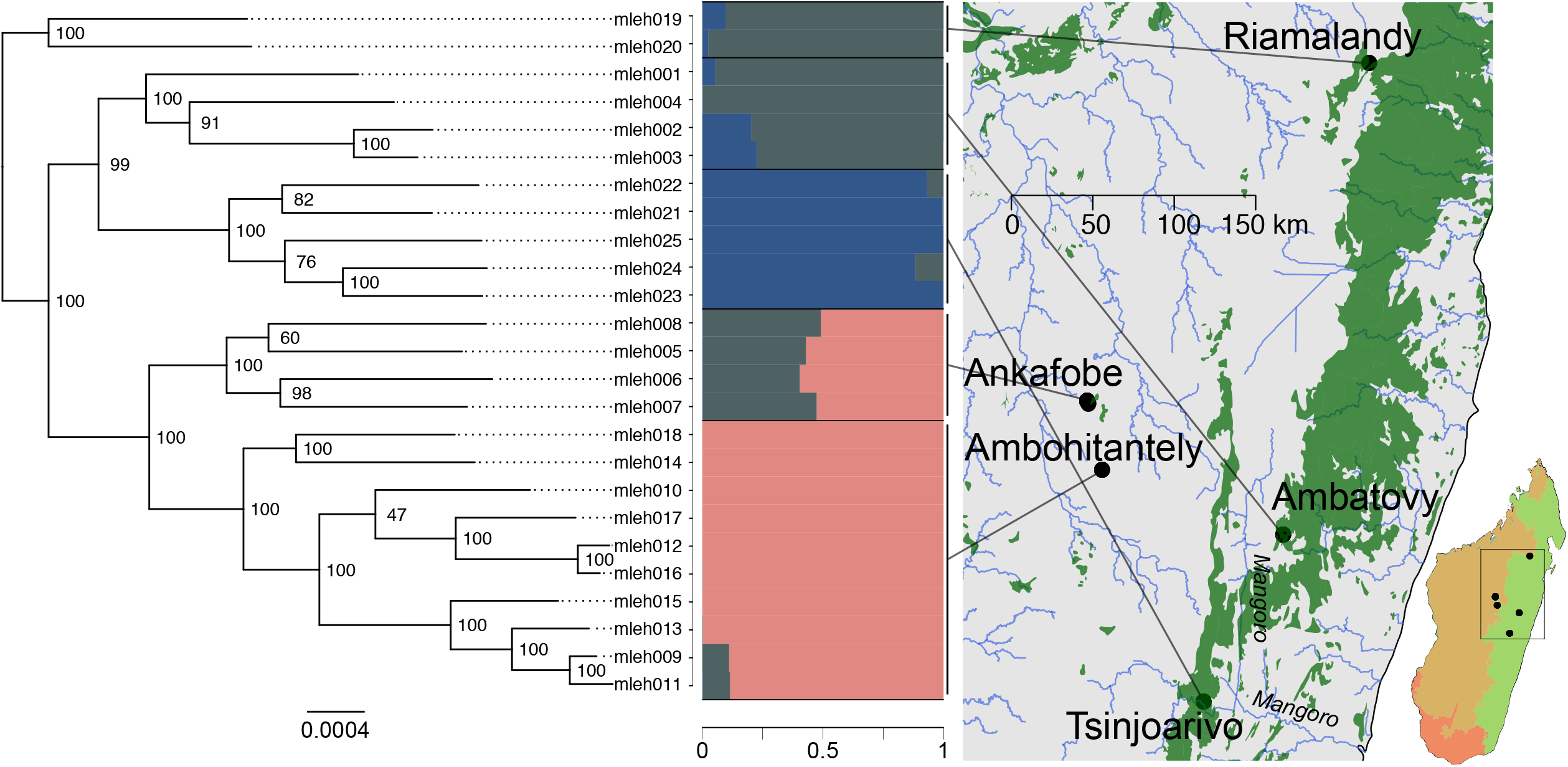
Well-supported phylogenetic lineages of *Microcebus lehilahytsara* reflect genetic structure. The phylogeny is the maximum likelihood tree with bootstrap support at nodes. Branch lengths are expected substitutions per site. Metadata for individual identifiers is available in Table S1. STRUCTURE results for the one SNP per locus dataset is shown with an optimal *k* = 3. The GPS coordinates of each sampling site are shown on the map, with Riamalandy, Ambatovy, and Tsinjoarivo found in the eastern rainforest while Ambohitantely and Ankafobe are fragmented forest patches in the Central Highland Savanna. Rivers are shown in blue while forest cover is shown in green. The specific region of Madagascar studied here shown in the boxed region on the country map. Major ecoregions of Madagascar are shown on the country map. Green shows the moist forest while brown is dry forest and red is spiny forest. Shape files of Madagascar’s ecoregions were taken from Vieilledent et al. (2016). All *M. lehilahytsara* sampling locations are considered wet forest.

AU tests rejected H_RT_, which came from SVDquartets with all variant sites or full loci. But the AU test could not reject H_RA_, despite H_TA_ being present in the majority of ML bootstrap trees and post burn-in trees (Figure 3). Bayes factors from SNAPP provided insights into the prevalence of both hypotheses across analyses. SNAPP supported lumping Riamalandy and Ambatovy together even with increasing step sizes and arguably more precise marginal likelihood estimates as opposed to lumping Ambatovy and Tsinjoarivo (Table 1). This result is surprising, considering H_TA_ was the maximum clade credibility tree from the posterior sample in the unconstrained analyses (Supplementary Figures S3 and S4). All phylogenetic evidence suggests a shared common ancestor for Ambatovy and Tsinjoarivo, but Tsinjoarivo may be more isolated due to barriers to gene flow and therefore more evolutionarily distant than expected. Phylogenetic analyses showed strong support for both CHS and eastern clades, and at least no very recent gene flow in the last few generations as evident by monophyly of our five sampling locations. However, such analyses do not rule out gene flow historical gene flow between the CHS and eastern forests.

**Figure 3.**
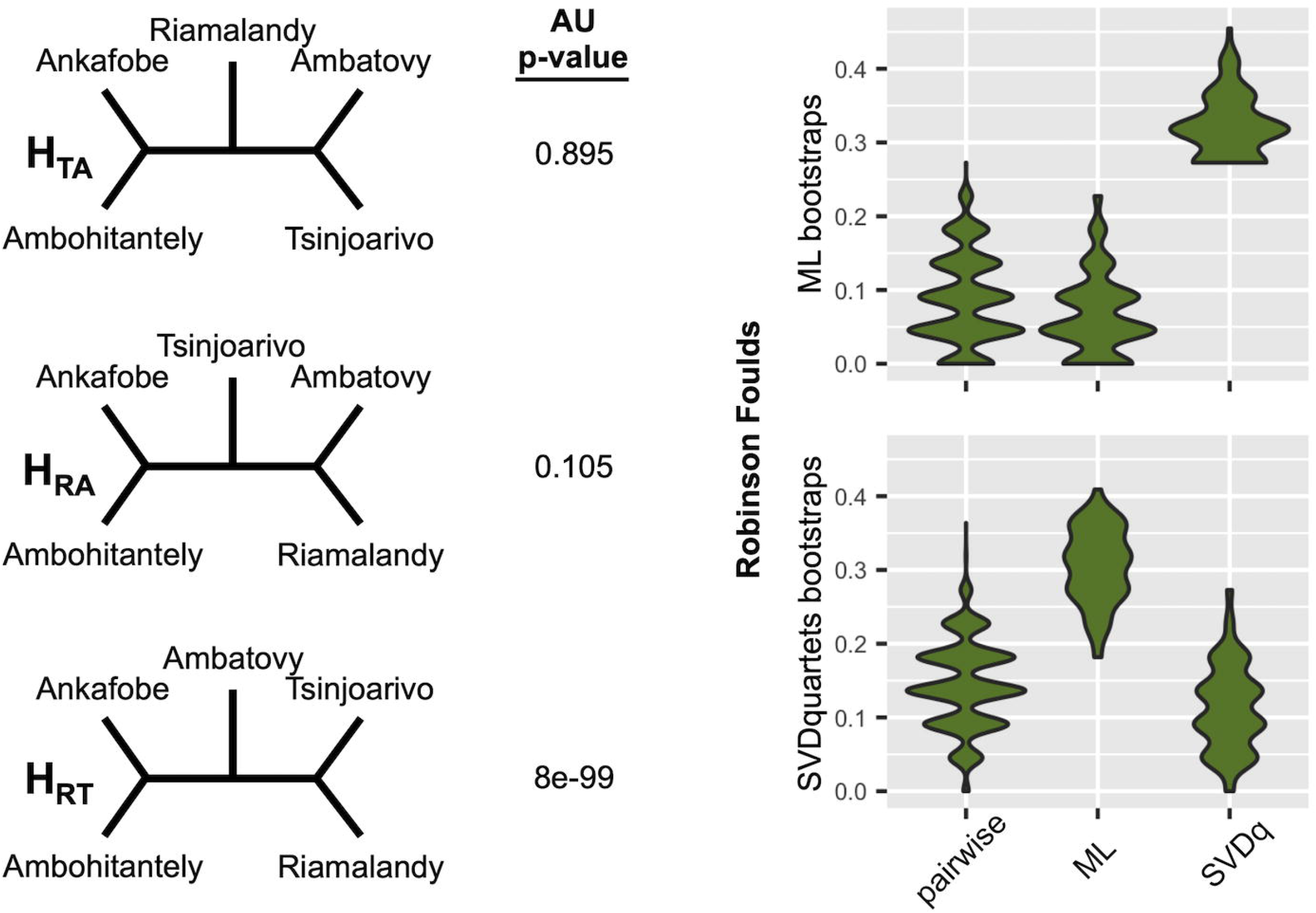
Phylogenetic hypothesis testing of eastern Madagascar. Three competing topological hypotheses were considered using both the AU test with site likelihoods and Bayes factors from SNAPP. Posterior probabilities of topologies were obtained from marginal likelihoods for SNAPP analyses and AU tests perform a multiscale bootstrap of site likelihoods to obtain a p-value that should be interpreted as strength against a topology. No analogous topology test exists for invariants, as in SVDquartets, but we did examine the RF distances among bootstrap trees from ML and SVDquartets analyses to summarize phylogenetic discordance among RAD loci. Pairwise are all bootstrap trees for ML or SVDquartets analyses compared with themselves. ML are all bootstrap trees, from either ML or SVDquartets analyses, compared to the ML tree. SVDq are all bootstrap trees, from either ML or SVDquartets analyses, compared to the SVDquartets bootstrap consensus tree. Although RF distances among SVDquartets bootstraps are typically higher, they are also closer to being normally distributed, implying either less uncertainty or potential biases in ML analyses.

**Table 1.** Marginal likelihood estimates and model selection with SNAPP.

| Steps | Topological Hypothesis | Marginal Likelihood | Bayes Factor<sup>†</sup> | Model Probability |
| --- | --- | --- | --- | --- |
| 20 | RA | -41399.18 | 0.00 | 1 |
|  | TA | -41847.70 | -897.05 | 1.61 x 10<sup>-195</sup> |
|  | RT | -41689.27 | -580.19 | 1.03 x 10<sup>-126</sup> |
| 40 | RA | -41531.26 | 0.00 | 1 |
|  | TA | -41848.94 | -635.35 | 1.08 x 10<sup>-138</sup> |
|  | RT | -42060.55 | -1058.58 | 1.36 x 10<sup>-230</sup> |
| 60 | RA | -41328.81 | 0.00 | 1 |
|  | TA | -41618.87 | -580.12 | 1.07 x 10<sup>-126</sup> |
|  | RT | -42057.39 | -1457.16 | 0 |

<sup>†</sup>Bayes factors computed by marginal log-likelihoods with respect to the best model as 2 \* (lnL<sub>i</sub> - lnL<sub>Max</sub>)

### Migration between eastern and Central Highland forests

Population genetic analyses revealed admixture between eastern and Central Highland *M. lehilahytsara* as well as population structure within the eastern rainforest (Tsinjoarivo versus Ambatovy and Riamalandy) that mirrored findings of SNAPP Bayes factors (Table 1). STRUCTURE analyses recovered an optimal *k* = 3 in 19 out of 20 sampled datasets (Supplementary Figures S9-S11). The remaining replicate recovered an optimal *k* = 2 (Supplementary Figure S11). All analyses supported a clustering of Central Highland and eastern *M. lehilahytsara*, with some degree of admixture between the two. The Ambohitantely site, while phylogenetically sister to Ankafobe, shares about half of its genotypes with Ankafobe and the other half with Ambatovy/Riamalandy. Tsinjoarivo often forms its own unique cluster, except in the one replicate where *k* = 2 and Tsinjoarivo clusters with other eastern forest populations.

Admixture between Central Highland and eastern *M. lehilahytsara* was also supported by analyses of genetic variation with *F*_*ST*_ statistics. Similar to STRUCTURE analyses, admixture between the CHS and eastern forests is largely limited to Ambohitantely and Ambatovy, respectively. Pairwise *F*_*ST*_ revealed similar levels of genetic divergence between Ambatovy and Ambohitantely to that of Ambatovy and Tsinjoarivo (Supplementary Table S3). The geographic structure between Central Highland and eastern *M. lehilahytsara* explains a small fraction of genetic variation, with most variation explained within populations (Supplementary Table S4). A Mantel test revealed little evidence for isolation-by-distance (pearson’s *r* = 0.09; *p* = 0.39), suggesting barriers to gene flow aside distance alone (Supplementary Figure S12).

Bayes factors were computed for different isolation-with-migration models to further investigate the patterns of admixture observed between the Ambatovy, Ambohitantely, and Ankafobe genetic clusters. The *n-islands* model was unambiguously found to be the best model for both the analyses that included all individuals modeled as two populations with a subset of data (Supplementary Figure 13; Supplementary Table 5) or three populations with all of the data (Figure 4). There were no large discrepancies in the estimated migration rates; however, the selected model suggested lower effective population sizes in Ankafobe and Ambohitantely compared to Ambatovy. Assuming a mutation rate of 1.64 × 10^−8^, MIGRATE estimated contemporary population sizes to be 12,652 for Ankafobe and Ambohitantely, and 17,226 for Ambatovy. Their population size estimates, although influenced by the prior, are not driven by the prior alone (Supplementary Figures S14-S16). Additionally, a higher population size in Ambatovy is supported across all models regardless of the migration and divergence scenarios (Figure 4). Because of Ambatovy’s higher effective population size, the number of migrants leaving towards Ambohitantely or Ankafobe per generation is approximately 1.2, compared to 0.8 effective migrants per generation leaving Ambohitantely or Ankafobe towards the east.

**Figure 4.**
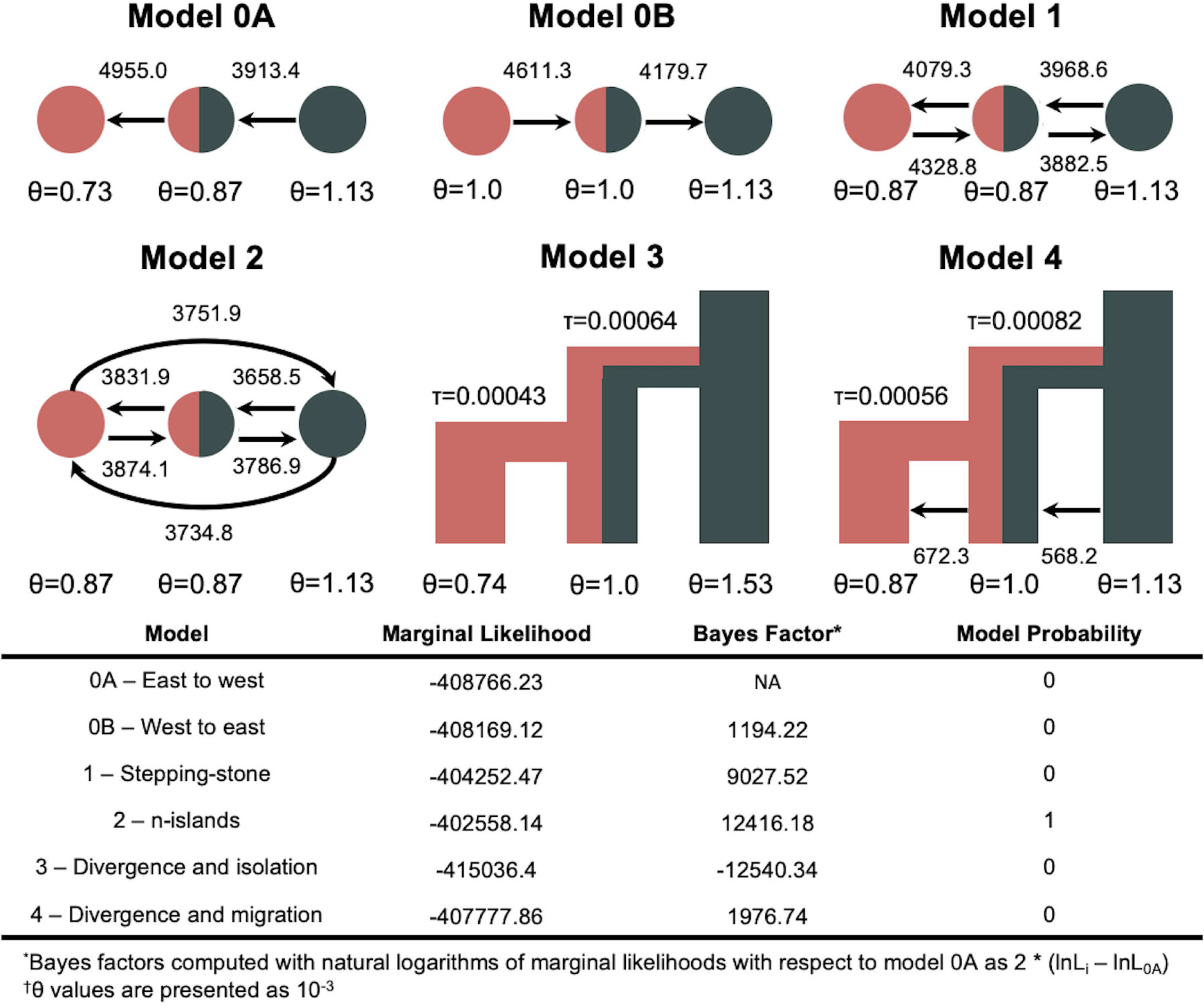
Migration models reveal gene flow between the Central Highlands and eastern forests. Bayes factors were used to test migration hypotheses between the eastern rainforest, Ambatovy (grey), and two fragmented forest patches in the Central Highlands, Ambohitantely (pink and grey) and Ankafobe (pink). These three locations were selected because of admixture detected in Ambatovy based on STRUCTURE analyses, and geographic locations are colored as such. θ parameters are scaled as 4*N*_*e*_μ and mutation-scaled migration parameters are shown near arrows. Divergence times are shown for relevant models in substitutions per site.

### Dating Population Divergences

Using the H_TA_ topology supported by Bayes factor analyses, divergence times were estimated using the MSC model. The topology was rooted with Riamalandy based on the analysis of mitochondrial data from *Microcebus lehilahytsara* and outgroup taxa (Supplementary Figure S17) and inference of clock-like trees from BPP (Supplementary Figure 5). Most MSC parameters converged across eight chains with potential scale reduction factors approaching 1 (Supplementary Table S6). A few parameters still retained high variation between chains for both θ and τ, but this discrepancy among chains was largely due to one or two chains with inefficient mixing for different prior combinations (Supplementary Figure S18-S25). Divergence times were not strongly influenced by the priors (Figure 5) and imply that divergence between the CHS and eastern forest, as well as the forest patches within them, occurred almost simultaneously. Absolute estimates inferred that divergence among our sampled lineages occurred near 25 KYA (Table 2).

**Figure 5.**
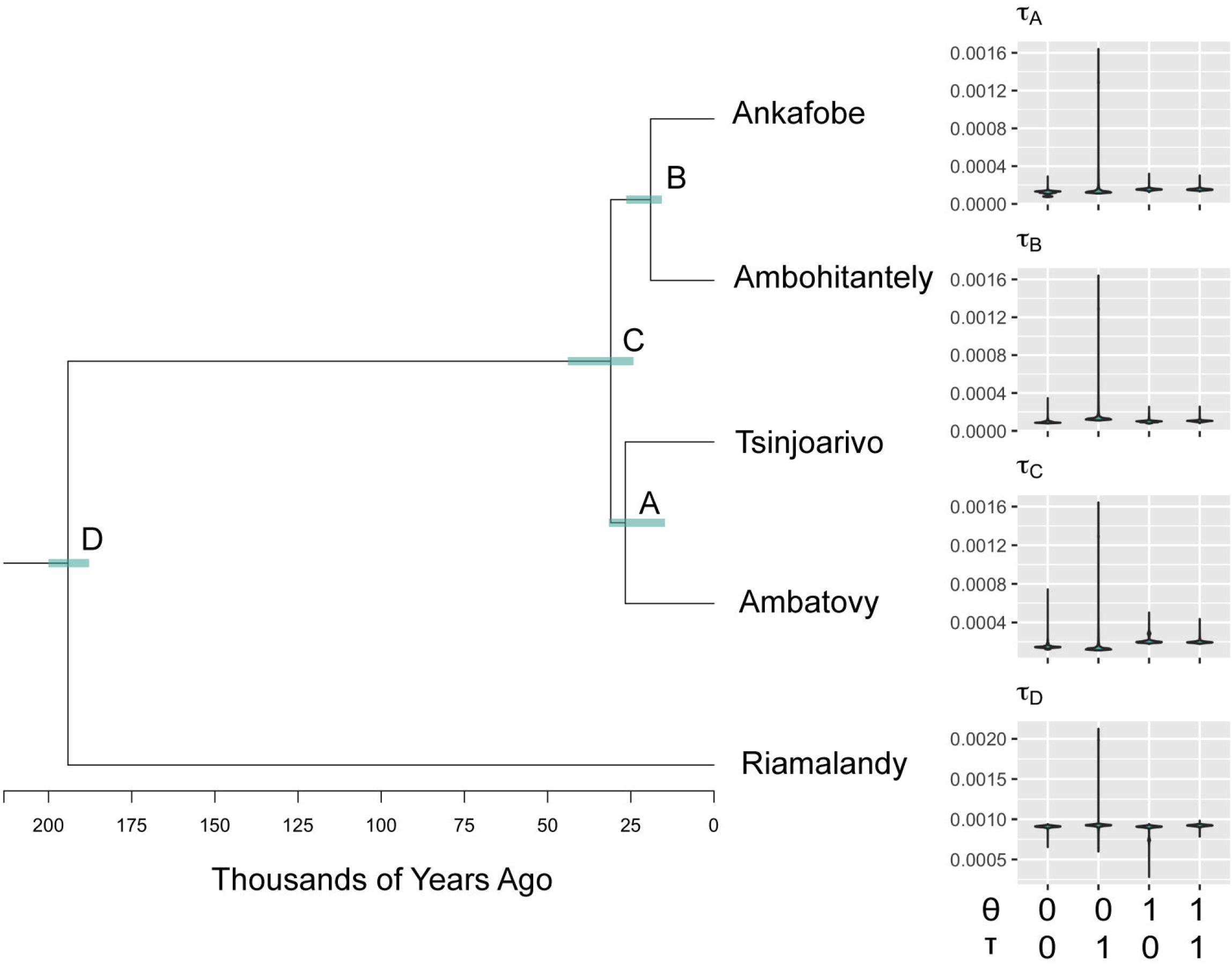
Divergence time estimates with BPP. Posterior distributions of MSC parameters for the combined posterior of prior choice 1. Branch lengths are in coalescent units and error bars represent 95% highest posterior density intervals around the median node height. Violin plots are the posterior distributions of node heights for prior choices 1 though 4 to show that while divergence time estimates are sensitive to prior choice, our biological conclusions are not.

**Table 2.** Absolute divergence times and population sizes estimated^†^ from BPP.

| Node | N<sub>e</sub> (x 10<sup>3</sup>) | 95% HPD | Divergence time (KYA) | 95% HPD |
| --- | --- | --- | --- | --- |
| Ankafobe | 19.2 | 17.3, 25.7 | - | - |
| Ambohitantely | 31.2 | 27.1, 40.9 | - | - |
| Tsinjoarivo | 23.1 | 14.0, 26.1 | - | - |
| Ambatovy | 55 | 33.5, 63.0 | - | - |
| Riamalandy | 204.5 | 193.4, 216.0 | - | - |
| A | 60.2 | 10.6, 201.3 | 26.7 | 15.9, 30.4 |
| B | 22 | 15.1, 34.2 | 19 | 16.9, 25.2 |
| C | 875.1 | 776.9, 955.0 | 31 | 25.4, 42.7 |
| D | 108.3 | 106.6, 110.1 | 194.1 | 189.0, 198.8 |

<sup>†</sup>Estimates are based on combined posterior for priors $\theta = 0$ and $\tau = 1$ .

### Population declines in the CHS

Population sizes experienced some slight contractions until 25 KYA (Figure 6) that coincide with the divergence of sampling locations (Table 2). Population sizes then increased through the Holocene and until the present except in Ankafobe. Ambatovy had consistently higher *N*_*e*_ estimates across replicates. Ankafobe had the lowest *N*_*e*_ estimates. Intermediate *N*_*e*_ estimates are observed for individuals at Ambohitantely and Tsinjoarivo. There is high uncertainty in *N*_*e*_ based on comparisons across replicates (Supplementary Figure S26), but individual analyses tended to converge (Supplementary Figures S27-S30).

**Figure 6.**
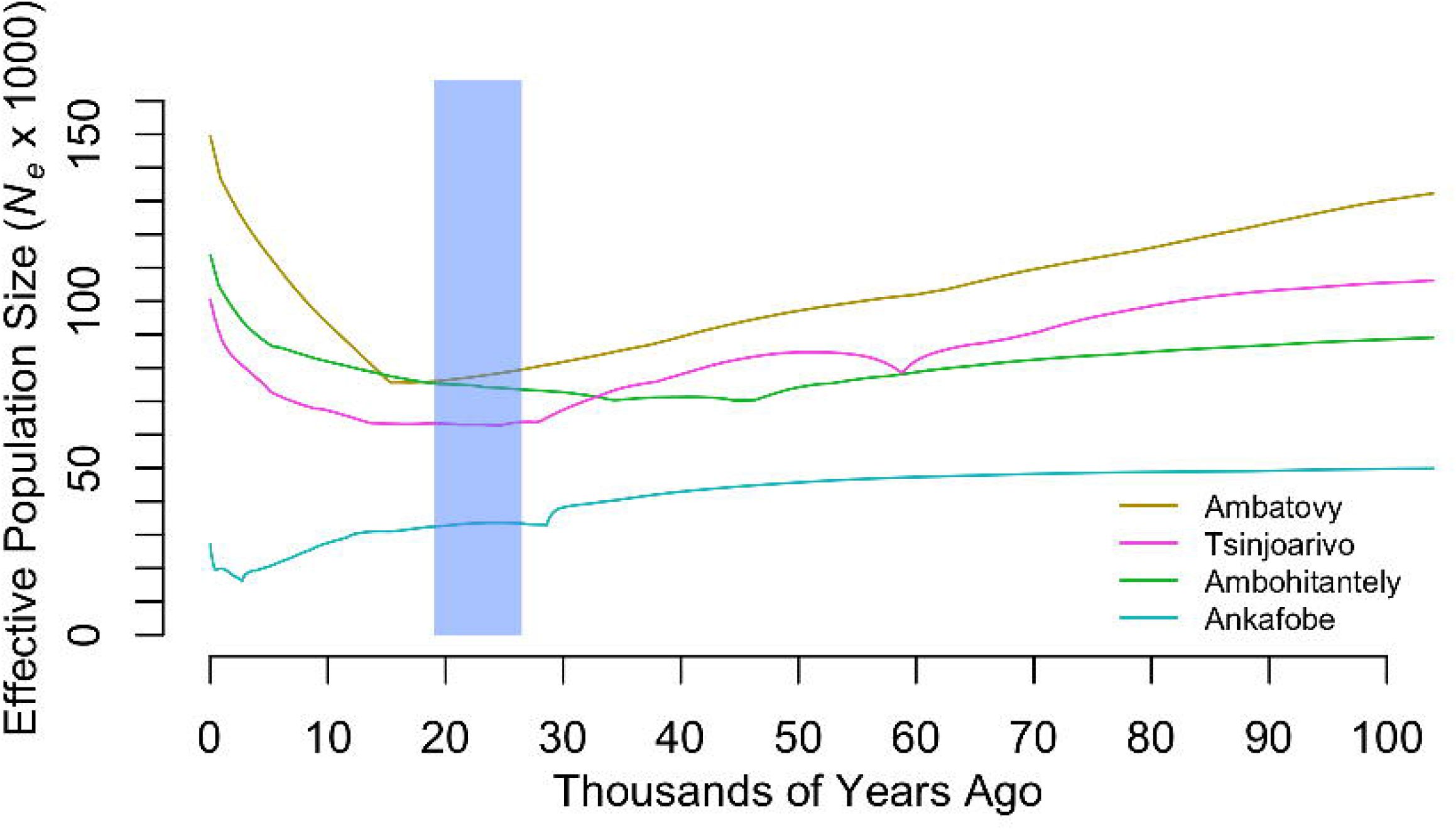
Population size change over time at sampling locations. Posterior distributions of θ at discretized bins of τ determined by the number of change points. Only the first sampled dataset is shown for each population. The blue box shows the duration of the LGM.

## Discussion

### Population structure of M. lehilahytsara predates human colonization of Madagascar

Our results demonstrated that *M. lehilahytsara* populations have been present in the CHS at least before the LGM. Phylogenetic analyses strongly supported a split between CHS and eastern forest sampling locations (Figures 2 and 3; Supplementary Figures S2-S8), although analyses of genetic structure (Figure 2; Supplementary Figure S11; Supplementary Figure 12; Supplementary Table S3) and migration rates (Figure 4) implied gene flow after the initial divergence. If, on the other hand, the divergence of *M. lehilahytsara* in the CHS and eastern forest had been recent and human mediated, we would expect at least some phylogenetic uncertainty due to low levels of genetic divergence, similar to Tsinjoarivo, Ambatovy, and Riamalandy. Three competing topological hypotheses were uncovered concerning the relationships among the eastern rainforest sampling locations, but the best possible species tree estimation and hypothesis testing implied the northern-most sampling location, Riamalandy, is sister to more southern sampling locations. All phylogenetic evidence suggested that *M. lehilahytsara* expanded as a ring around the higher elevated regions of the CHS and that the CHS acted to some extent as a barrier to gene flow during a period of vegetational change and aridification at the LGM (Burney et al., 2004; Samonds et al., 2019). Although, the northern end of this barrier may have remained permeable even through climate change, watersheds may have buffered the region from the aridification that is more evident in the southern extent of the CHS (Wilmé, Goodman, & Ganzhorn, 2006; Muldoon et al., 2012; Yoder et al., 2016; Samonds et al., 2019).

Rivers are known to function as barriers to gene flow in mouse lemurs (Olivieri et al., 2008), and our data suggest they are more significant in impeding mouse lemur migration than open grasslands. Tsinjoarivo likely represents a population isolated from Ambatovy and Riamalandy by biogeographic barriers. *M. lehilahytsara* from Tsinjoarivo was previously recognized as genetically distinct based on a study of few mitochondrial and nuclear loci, although Bayes factors did not indicate Tsinjoarivo to be an independent lineage under the MSC (Hotaling et al., 2016). The findings of Hotaling et al. (2016) were recapitulated here with our RADseq data, with little to no allele sharing between Tsinjoarivo and Ambatovy or Riamalandy (Figure 2; Supplementary Figure S11). The isolation of Tsinjoarivo from other eastern sampling locations is plausible, because in addition to geographic distance, our sampling locations are separated by the Mangoro River (Figure. 2). BPP analyses estimate the time of separation between Tsinjoarivo and Ambatovy to be near 27 KYA, which again coincides with a time of cool and dry conditions during the LGM (Scholz et al,. 2011; Clark et al., 2009).

### The mosaic of grassland and wooded savanna was permeable by mouse lemurs

Three lines of evidence support the hypothesis that *M. lehilahytsara* had a large ancestral distribution that covered both the CHS and eastern rainforest escarpment. Within this large geographic area, population substructure would have been formed due to paleoclimatic oscillations but with the maintenance of some level of post-divergence gene flow. First, based on our coalescent estimates of divergence times, the initial divergence between the CHS and eastern forests would have formed during a period coincident with the LGM (Figure 5 and Table 2). Mean ambient temperatures would have been 4°C cooler and precipitation much lower at this time (Burney et al., 2004), which may have caused some individuals to retreat towards watershed refugia (Wilmé et al., 2006) while others remained in the CHS. The divergences between *M. lehilahytsara* from Ankafobe and Ambohitantely, Ambatovy and Tsinjoarivo, and their common ancestors occurred near-simultaneously, and this result is not an artifact of prior choice from Bayesian MSC analyses (Supplementary Figures S18-S24). The pattern of divergence and population structure formed by reduced gene flow is reflected in θ estimates from BPP. MSC estimates suggested an increased population size proceeding divergence events (Table 2), which likely explains why a larger prior on θ resulted in a higher posterior probability and reduced uncertainty in the divergence time for the MRCA of CHS and eastern *M. lehilahytsara*.

Second, the strong preference for an *n-islands* model from MIGRATE (Figure 4; Supplementary Figure S12; Supplementary Table S5) also supported the demographic scenario of populations that diverged simultaneously from a single larger distribution. Although there is genetic structure, there is little discernable direction in gene flow, which may explain why models that incorporated divergence did not perform as well as the *n-islands* model. The migration rates inferred for our data are arguably high, between 0.8 and 1.2 migrants exchanged per generation. Such migration rates can reduce fixation indices (e.g. Hartl and Clark, 1989) and explain the relatively high proportion of admixture in Ambohitantely (Figure 2; Supplementary Figure S11) and low *F*_*ST*_ despite the many generations that must have passed since the initial divergence between the CHS and eastern forest populations (Supplementary Tables S3 and S4). Although, migration model parameters may be difficult to identify with many migration rate parameters (Beerli and Palczewski, 2010) and parameter identifiability is a concern for our short RAD loci (Supplementary Figures S14-S16) that individually contain very few informative coalescent events, we are nonetheless confident in these results. Given that we acknowledge the challenges of model testing and parameter estimation, the *n-islands* model agreed with the MSC results in that eastern *M. lehilahytsara* has a higher *N*_*e*_ than *M. lehilahytsara* in the CHS. Consistent with our model-based hypothesis testing, analyses of genetic variation with *F*_*ST*_ suggested that the mosaic of grassland and wooded savanna in the Central Highlands may have posed as much of a barrier to gene flow as the Mangoro river (Supplementary Table S3; Figure 2). Relatively little of the genetic variation is explained by grouping populations by the CHS and eastern forest compared to variation within populations (Supplementary Table S4).

The third piece of evidence for *M. lehilahytsara’s* large ancestral distribution is that analyses of population size recovered similar demographic trends at all four sampling locations analyzed with EBSP methods. Both the CHS and eastern forests reflect a slight population decrease before the LGM followed by a recovery through the Holocene (Figure 6; Supplementary Figures S26-S30). Because we do not see changes in the CHS while the eastern forests remained stable, population size changes are likely due to global or Madagascar-wide effects rather than smaller-scale regional changes like agriculture. Although, there are large disparities in *N*_*e*_ estimates between EBSP and other methods, which raises some concern for interpreting absolute population size values from molecular data, patterns inferred from the different analyses are consistent.

### Making conservation biology predictions from models of molecular evolution

Here we provided a range of present-day *N*_*e*_ estimates that are sensitive to model (i.e. isolation-with-migration, MSC, Extended Bayesian Skyline) and prior choice, but all analyses suggest *M. lehilahytsara* populations sizes are healthy (i.e. > 5000; Lynch and Lande, 1998) and have likely increased through the Holocene. The most recent IUCN evaluation of *M. lehilahytsara* considers the species to be vulnerable (Andriaholinirina et al., 2014c) while other mouse lemur species in the eastern rainforest are considered endangered (Andriaholinirina et al., 2014a; Andriaholinirina et al., 2014b; Baden et al., 2014). If *M. lehilahytsara* has adaptive traits that allowed the species to expand its range into the CHS while related species were restricted to lowland forests, *M. lehilahytsara* may have been more robust to Pleistocene climate change and even more recent anthropogenic disturbances. *M. lehilahytsara* has been regarded as a highland specialist (Radespiel et al., 2012) and has been documented to be able to enter a state of prolonged torpor (Blanco et al., 2017; Andriambeloson et al., 2020). We hypothesize that the capacity for extended torpor in *M. lehilahytsara* is an adaptive trait that has facilitated its increased geographic range compared to most other mouse lemur species. Although individual *M. lehilahytsara* populations in the CHS have maintained smaller population sizes compared to those in the eastern forests (Table 2; Figure 6; Supplementary Figure S26), there is increasing evidence that these populations are robust. For example, the Ankafobe forest patch is less than one Km^2^ yet the resident population is thriving (Blanco et al., in review).

We emphasize, however, that *M. lehilahytsara* in the CHS is not simply some sink population from an eastern forest source. CHS populations have been evolutionarily independent from the eastern escarpment for approximately 5000 generations if we consider the lower end of the 95% HPD intervals of divergence times for nodes A and B in our BPP analyses (Table 2). Even so, CHS populations have exchanged migrants (Figure 4) with the eastern forests despite the historically open grassland-woodland mosaic (Solofondranohatra et al., 2018). Indeed, the occasional migration to and from the east may have been important for sustaining large *N*_*e*_ despite the small range sizes. Our best migration model estimated an *N*_*e*_ of 13,000 for both Ankafobe and Ambohitantely and MSC estimates are 19,000 and 31,000 respectively. There are of course discrepancies in the absolute *N*_*e*_, but our results suggest that *N*_*e*_ has remained sufficiently large such that the strength of selection has been strong enough to overcome fixation of deleterious alleles by drift alone (Lynch & Lande, 1998). Nonetheless, the sustained lower *N*_*e*_ in the CHS compared to the eastern forests may imply a negative consequence of continued habitat loss and fragmentation if human activities are not mitigated.

### The importance of interpreting biodiversity through evolution

The processes that shaped existing *M. lehilahytsara* population structure must have occurred before human arrival and mid-Holocene droughts. Although charcoal records (Burney,1987a; Burney 1987b), stratigraphic pollen occurrence (Gasse & Van Campo, 1998; Burns et al., 2016), and megafauna subfossil extinctions (Hansford et al., 2018) all point to human-mediated large-scale ecological changes in the Central Highlands (Virah-Sawmy et al., 2010), our analyses implied that fundamental *M. lehilahytsara* population divergence coincides with Pleistocene climate change near the LGM – not after. A similar narrative has been recovered in other analyses of lemur populations (Quémére, Amelot, Pierson, Crouau-Roy, & Chikhi, 2012; Salmona et al., 2017) and in an endemic olive tree (Salmona et al. 2019). In summary, phylogeographic analysis of multiple species and localities agree with ecological (Bond et al. 2008) and evolutionary (Vorontsova et al., 2016; Hackel et al., 2018) studies of grasses that demonstrate that Malagasy savannas are of ancient origin and have long persisted as a mix of open grassland and woodland. Although fires due to human activity continue to ravage the Central Highlands (Burney, 1987a; Gasse & Van Campo, 1998; Virah-Sawmy et al., 2010; Burns et al., 2016; Samonds et al., 2019), leading to devastating forest loss in Madagascar (Vieilledent et al., 2018), ongoing habitat fragmentation is occurring on a background of habitat mosaicism that is likely associated with Pleistocene climate change.

## Ethics

Research was conducted under permits N 0233/07/MINENV.EF/SG/DGEF/DPSAP/SSE (Tsinjoarivo), N 229/14/MEEF/SG/DGF/DCB.SAP/SCB (Ambatovy), and N225/15/MEEMF/SG/DGF/DAPT/SCBT (Ankafobe) issued by the Ministry of Environment, Water and Forests of the Malagasy Government.

## Data Accessibility

All new sequence data has been made available through NCBI BioProject PRJNA560399. All individual SRA identifiers are available in Supplementary Table S7. Alignments and other data for analyses are available from the Dryad Digital Repository (Tiley et al., 2020).

## Author Contributions

GPT and ADY conceived the study. MBB, JMR, and RMR performed the fieldwork. ARS and PAH constructed RADseq libraries. GPT performed analyses. GPT, MBB, and ADY wrote the manuscript. All authors read and approved the final manuscript.

## Competing Interests

The authors declare no competing interests.

## Funding

The research was supported by Duke Tropical Conservation Initiative Grant to ADY, CI/Primate Action Fund and Duke Lemur Center/SAVA Conservation research funds to MBB, and Duke University research funds to ADY. ADY would also like to acknowledge support from the John Simon Guggenheim Memorial Foundation and the Alexander von Humboldt Foundation.

## Acknowledgements

This is Duke Lemur Center publication #XXXX.

## Notes

### Competing Interest Statement

The authors have declared no competing interest.

