## Supplementary Material for "Population genomic structure in Goodman’s mouse lemur reveals long-standing separation of Madagascar’s Central Highlands and eastern rainforests"

### Supplementary Information for

#### Phylogeographic analysis of Goodman's mouse lemur reveals historical interconnectivity of Madagascar's Central Highlands and eastern rainforests

**George P. Tiley[1,†], Marina B. Blanco[1,2], José M. Ralison[3], Rodin M. Rasoloarison[3,4], Amanda R. Stahlke[5], Paul A. Hohenlohe[5], Anne D. Yoder[1]**

[1]Department of Biology, Duke University, Durham, NC 27708, USA

[2]Duke Lemur Center, Duke University, Durham, NC 27705, USA

[3]Département de Biologie Animale, Université d'Antananarivo, BP 906, Antananarivo 101, Madagascar

[4]Behavioral Ecology and Sociobiology Unit, German Primate Centre, 37077 Göttingen, Germany

[5]Institute for Bioinformatics and Evolutionary Studies, Department of Biological Sciences, University of Idaho, Moscow, ID 83844, USA

#### Supplementary Methods

*Data assembly with STACKS*

Although there exists a well-assembled genome for the close relative of *M. lehilahytsara*, *M. murinus*, we chose to utilize *de novo* assembly methods in order to utilize the paired reads for our specific library preparation method, which was not processed by available reference-based methods at the time of writing. Because all nucleotide sites are important for correctly estimating equilibrium nucleotide frequencies and Gamma-distributed among-site rate variation parameters, we constructed fastas for loci circumscribed by STACKS from the raw fasta output while merging alleles with proper IUPAC codes with a custom Perl script *processLoci.pl*.

We used the RAD loci assembled with paired-end data to potentially improved orthology inference by removing loci that could represent the assembly of reads from paralogous genomic regions. This was accomplished by performing a BLASTN v2.6.0 search between each sequence for each cluster against the *M. murinus* 3.0 genome (GCA_000165445.3); if a sequence aligned to at least two different regions of the genome where the second-best hit had a bit score of that was within 5% of the best hit, we considered paralogy to be a problem for this sequence. All potentially paralogous sequences and loci were discarded from further analyses.

*Phylogenetic Estimation*

All SNAPP analyses applied a gamma prior on theta with alpha = 3 and beta = 5769 to get mean of 0.00052. An assumed mean theta of 0.00052 is biologically realistic based on previous analyses of nuclear protein-coding genes among population of mouse lemurs (Blair et al. 2014). A gamma prior was also used on tree height, $\Gamma(3,0.00234)$, assuming a root height of 0.001 and 5 lineages under the Yule prior to get mean of 1283.3. Lambda was constrained between 0.00001 and 10000 to prevent sampling extreme values that might make the MCMC algorithm less efficient. Species tree analyses ran for 10000000 generations sampling every 500 generations to get posteriors of 20000 samples. The first 1000000 samples were skipped as pre-burnin for two independent chains when conducting species tree analyses. Log-likelihoods were sampled efficiently in our SNAPP species tree analysis, so fewer samples were used for Bayes factor analyses. Each step ran for 1000000 generations after a 200000 generation burn-in, while sampling every 100 generations for posterior samples of 10000 each.

Species tree estimation with BPP used a uniform distribution on topologies. The prior on θ was distributed as inverse gamma with α=3 and β=0.0011 to get a fairly diffuse distribution around an approximate mean of 0.0055. Although, we used analytical integration of θ with a conjugate prior (Hey and Nielsen 2007) for species tree estimation. We used what we consider an appropriate prior given previous species tree estimation with coalescent methods for mouse lemurs (Yoder et al. 2016), which assumed τ ~ invGamma(3,0.002). This returns a mean root age near 0.001 coalescent units or 229 KYA assuming a mutation rate of 1.64 x $10^{-8}$ (Campbell et al. 2019) and generation time of 3.75 years. We investigated the effect of the prior choice on species tree estimation, and because the interpretation of our results relies heavily on our ability to translate coalescent time to absolute time and the underlying topology. Therefore, we also performed species tree estimation with a misspecified prior of τ ~ invGamma(3,0.2) and θ ~ invGamma(3, 0.11). This pushes the root age back to 22.9 MYA and would at least affect branch lengths if not the topology in the case where our RADseq data are not informative. We used four different prior combinations 1) θ ~ invGamma(3, 0.0011) and τ ~ invGamma(3,0.002) (prior 00), 2) θ ~ invGamma(3, 0.0011) and τ ~ invGamma(3,0.2) (prior 01) , 3) θ ~ invGamma(3, 0.11) and τ ~ invGamma(3,0.002) (prior 10), and 4) θ ~ invGamma(3, 0.11) and τ ~ invGamma(3,0.2) (prior 11). Eight independent chains of 2500 samples were collected after a 1000 sample burnin. We sampled every 100 generation and combined posteriors after evaluating convergence.

*Testing migration hypotheses among putative populations*

Because MIGRATE analyses are computationally demanding, we only analyzed phased data (two alleles per individual) for 4 individuals from the Ankafobe, Ambohitantely, and Ambatovy sites. These represent two genetically distinct Hardy-Weinberg groups and an admixed group (Ambohitantely) based on STRUCTURE results. We used the HKY substitution model, with kappa fixed at 4.762. HKY was selected by the difference in the Bayesian Information Criterion with PAUP, from the models available in MIGRATE, and we used that kappa estimate for the data being analyzed with MIGRATE. Although accounting for rate variation does improve model fit, the alpha estimate was very low, 0.017, and we saw little gain for building more model complexity into our MIGRATE analyses. For the θ prior, humans have a mutation rate of $1.2 \times 10^{-8}$ (Scally and Durbin 2012) and mice have a mutation rate of $0.54 \times 10^{-8}$ (Uchimura et al. 2015). Because a germline mutation rate for mouse lemur was not available when beginning MIGRATE analyses, we simply took the average of human and mouse to be $0.87 \times 10^{-8}$. Our best ideas of effective population size estimates for mouse lemurs tend to be between 10000 and 20000 before forest fragmentation (Olivieri et al. 2008). To get some mean for a prior distribution on theta we consider $\theta = 4N_e\mu$. Thus, $\theta = 4*(15000)*(0.87 \times 10^{-8}) = 0.000522$. Then, we assume a gamma distribution with an alpha of 4 to keep the prior fairly diffuse. A maximum θ of 0.2 was assumed, as this is well outside of the observed estimates of theta for nuclear protein-coding genes (Blair et al. 2014) and mitochondrial data (Schneider et al. 2010); this prevents sampling of biologically unrealistic values that could skew posterior summary statistics. The mutation scaled migration rate (*M*) used a uniform prior, but we increased the maximum bound to 100000. This provides an upper bound of 52 effective migrants per generation ($N_m$), assuming a θ of 0.000522 and $N_m = \theta M$. Efficiency of the MCMC algorithm was optimized by checking effective sample sizes for model parameters and mixing with TRACER (Rambaut et al. 2018) v1.7.

Each analysis was run with 4 replicates of 4 chains, with one cold chain and three heated chains. We sampled every 100 generations, discarding the first 10000 generations as burn-in and sampling 30000 generations for the posterior sample. We saved posterior samples from each analysis to MIGRATE's bayesallfile for evaluating mixing and if our sampling was sufficient with effective sample sizes. Although, analyzing convergence can be difficult with MIGRATE, we plotted posterior samples of thetas and migration parameters to look for smooth distributions. We also ran MIGRATE analyses without data so we could evaluate marginal priors; by comparison with posteriors we could assess if there was information about specific model parameters captured by the data, and if these parameter estimates would benefit from more sampling.

*Divergence time estimation with BPP*

Our primary goal was to estimate the divergence time between the Central Highland fragments (Ankafobe and Ambohitantely) and the eastern rainforest (Tsinjoarivo, Ambatovy, and Riamalandy). Therefore we estimated multispecies coalescent model parameters with BPP. Because of the computational rigor required to estimate multispecies coalescent parameters well, we reduced our sampling to two individuals per sampling location and only used RAD loci without missing data for these ten individuals. This led to 7022 RAD loci used for analysis while allowing us to estimate ancestral $\theta$ parameters for the 5 sampling locations and 4 internal nodes as well as 4 divergence times in coalescent units. For a $\theta$ prior, we used the inverse gamma distribution of BPP v4.0 with an alpha of 3 and beta of 0.0011. This provided a mean $\theta$ prior estimate of 0.00052, based on the same lines of evidence used for MIGRATE analyses. For divergence times, we used the default prior of an inverse gamma with alpha of 3 and beta of 0.002 for the root and a dirichlet with alpha of 3 for the other internal nodes.

In order to get a sufficient number of independent posterior samples, we ran eight independent chains and combined chains after evaluating convergence with CODA (Plummer et al. 2006) and TRACER (Rambautet al. 2018). Each chain sampled every 100 generations. We discarded the first 1000 samples (10000 generations) as burn-in and logged the following 2500 samples. To evaluate convergence of individual parameters, we estimated potential scale reduction factors among chains and plotted the potential scale reduction factor over increasing generations to observe if convergence diagnostics were likely to improve with additional sampling. We also sampled the posterior with the MCMC algorithm without data to obtain marginal priors and infer if there were any model parameters for which our data contained little information. We used a similar experimental design with misspecified priors as described earlier to evaluate sensitivity of parameter estimates to prior choice.

We do have uncertainty in rooting, what we consider, a population-level phylogeny with any one population, in this case Riamalandy. Mitochondrial data does support this decision (Fig. S17), as does the rooted species tree estimated from posterior distributions in BPP's species tree analysis (Fig. S5) and unconstrained SNAPP analyses (Fig. S3).

#### Supplementary Figures

**Figure S1 – Environmental variation across sampling sites.** Data comes from WorldClim at 2.5 second resolution. Ankafobe and Ambohitantely are typically subject to drier conditions and greater seasonality in the CHS compared to Ambatovy and Tsinjoarivo on the eastern rainforest edge.

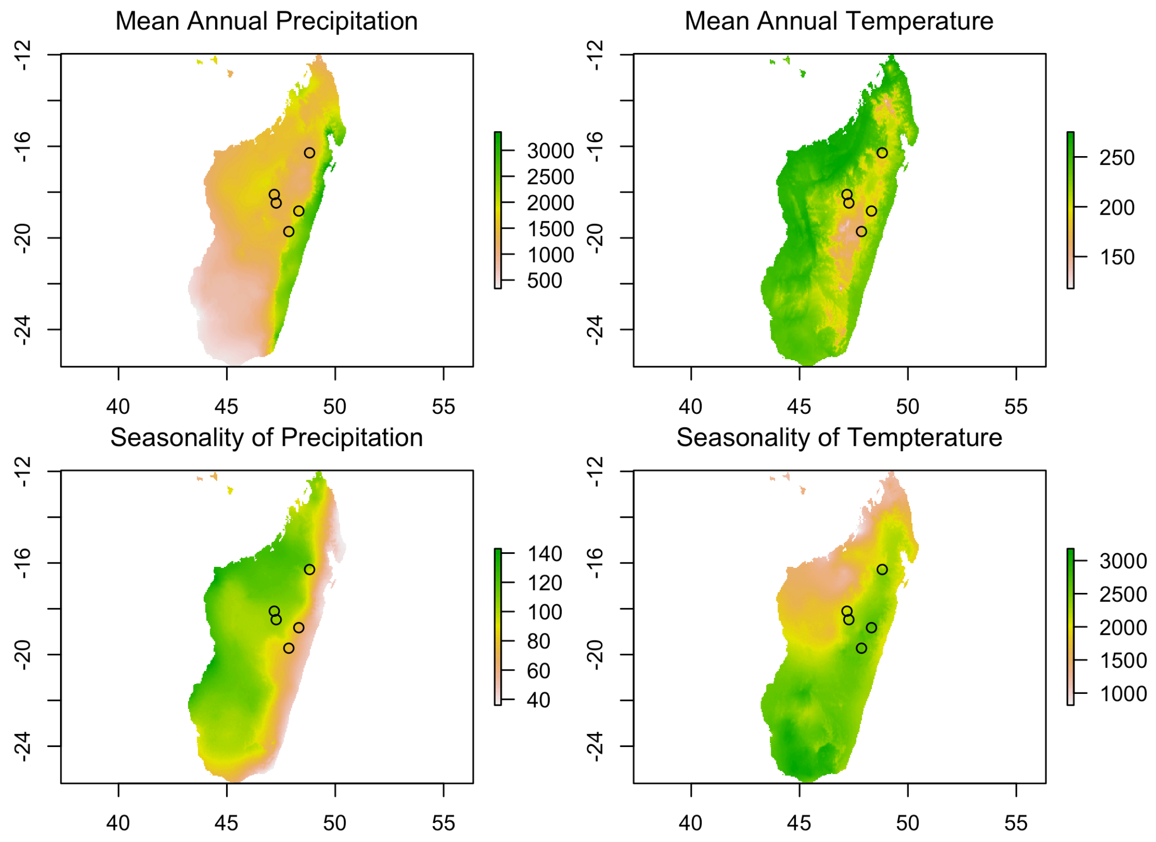

**Figure S2 – ML tree with branch lengths and bootstrap support from RAxML.** Bootstrap replicates were made from concatenated and unpartitioned RAD loci with at least 4 taxa present. Branches are colored by sampling location.

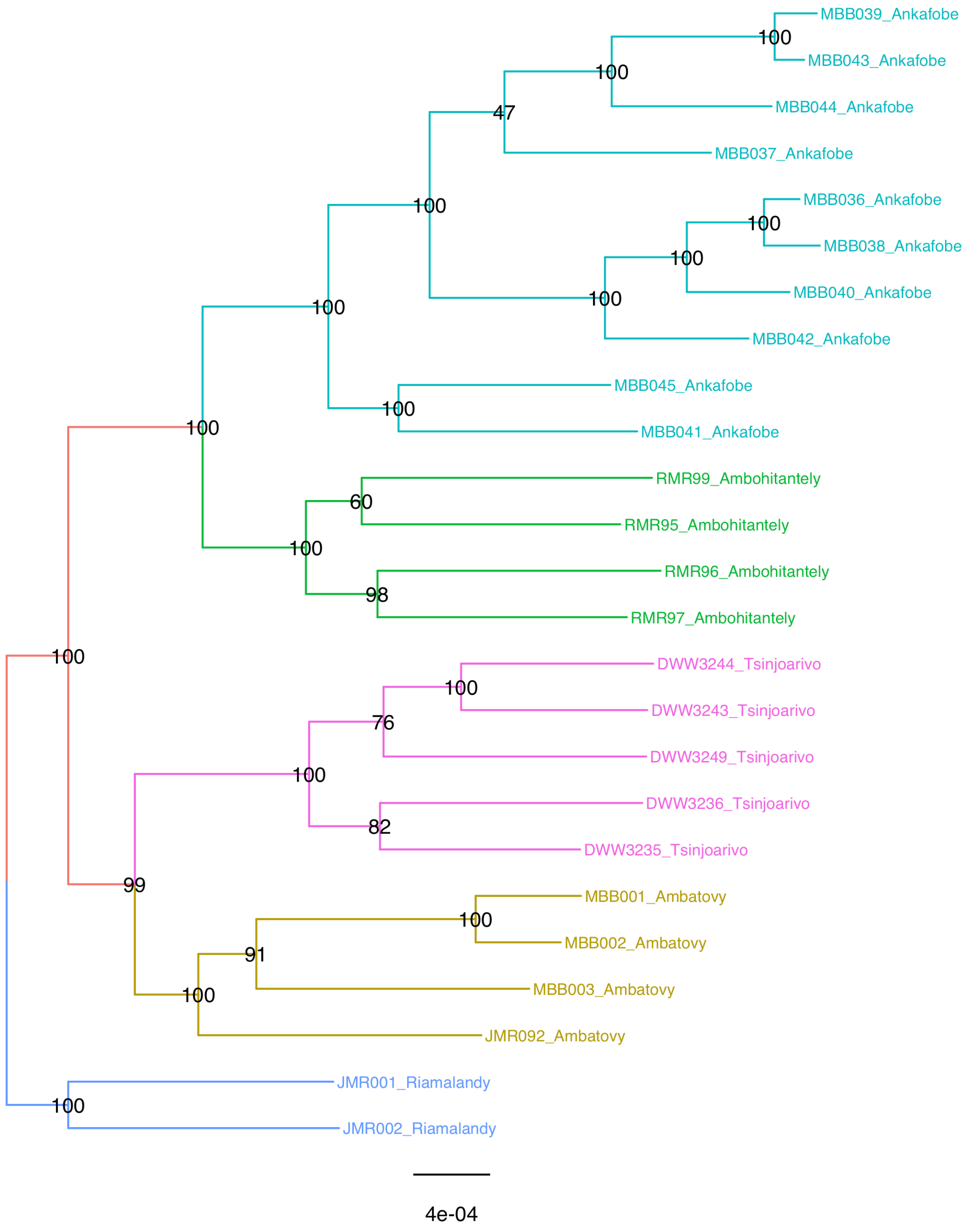

**Figure S3 – MCCT trees from SNAPP.** Sampling locations were treated as population identifiers. Branch lengths are substitutions per-site and node labels are posterior probabilities from 20000 post-burnin trees.

Chain 1

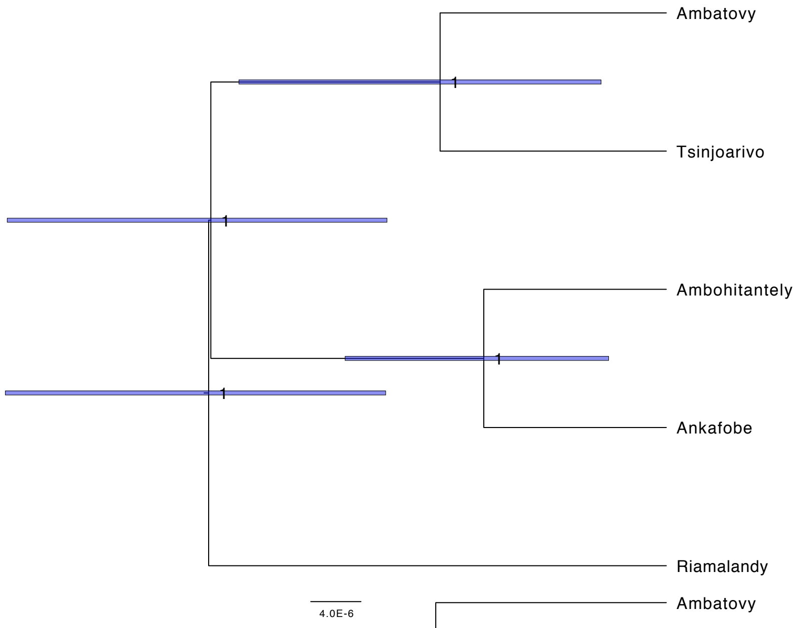


Chain 2

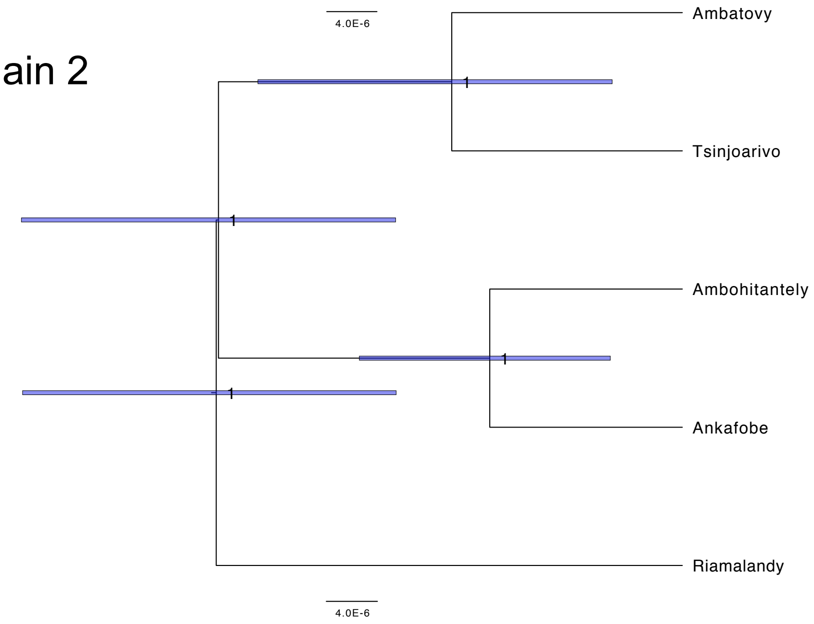

**Figure S4 – Convergence of posterior distributions from SNAPP.** One chain is red while the other is blue, with the overlap showing strong agreement in posterior parameter distributions. Node heights for the two runs are also given. Node “D” is the root and “B” is the MRCA of Ankafobe and Ambohitantely.

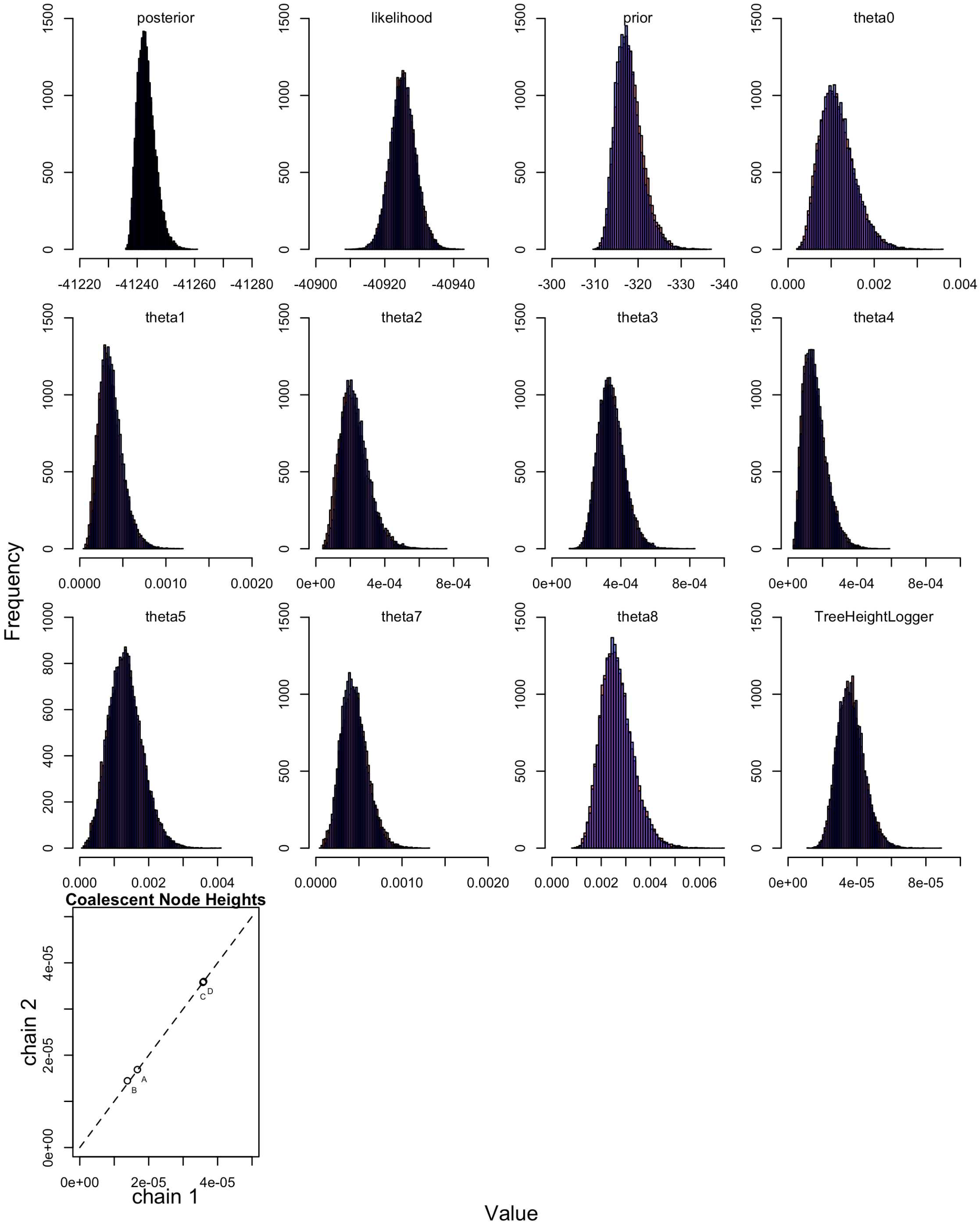

**Figure S5 – MCCTs estimated by BPP.** Trees are summarized from posteriors of 20000 samples with branch lengths in mean substitutions per site. The lower posterior probability for the contentious node between SVDquartets and other methods is only evident when perhaps θ is too low. Prior choices 10 and 11 produced tight HPDs. Mean divergence time estimates were nearly identical regardless of prior choice.

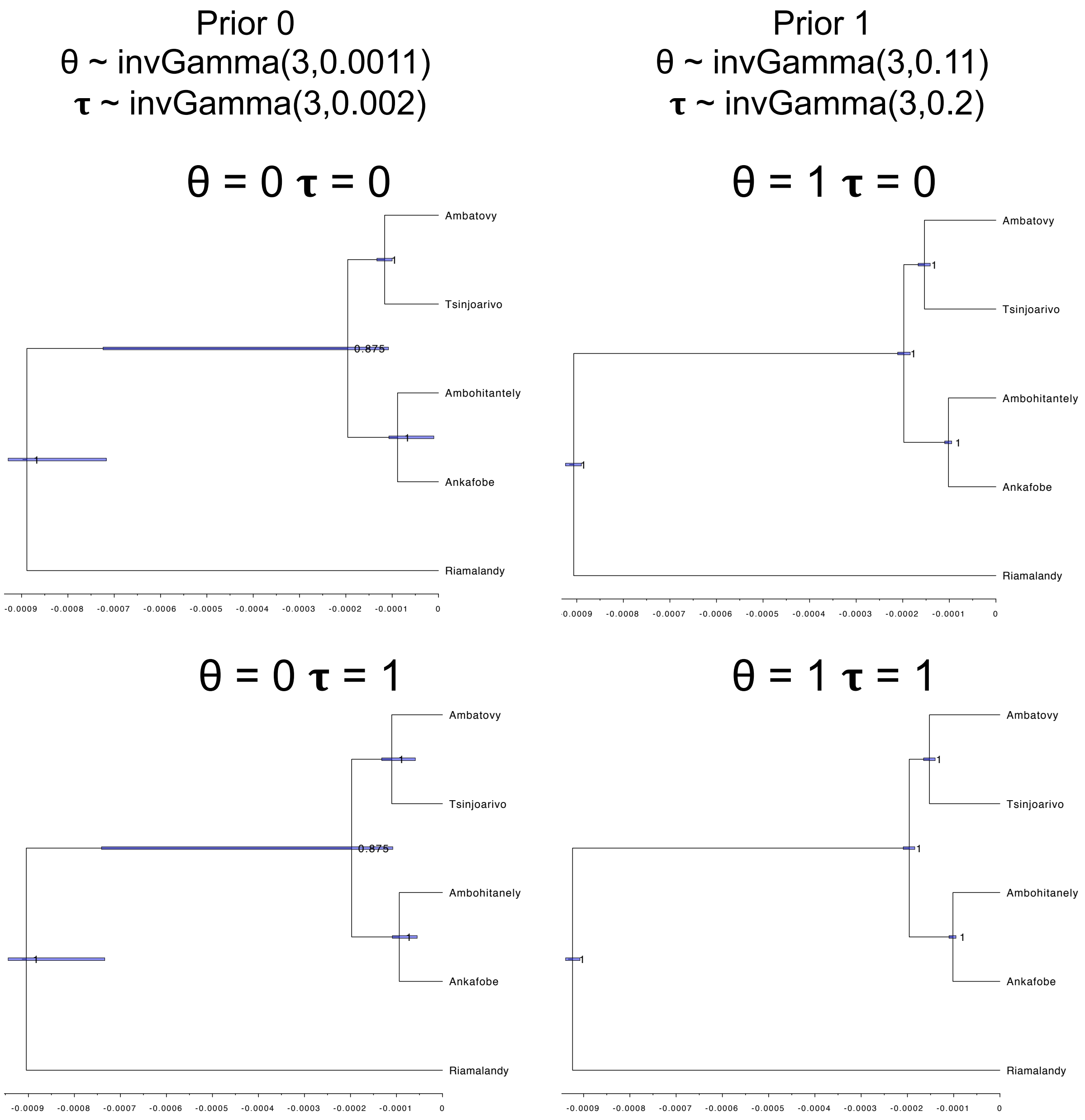

**Figure S6 – Bootstrap Consensus Tree Estimated by SVDquartets with variable sites only.** 100 bootstrap replicates were performed on a dataset that included all variable sites with at least 4 taxa per site. Branches are colored based on sampling location.

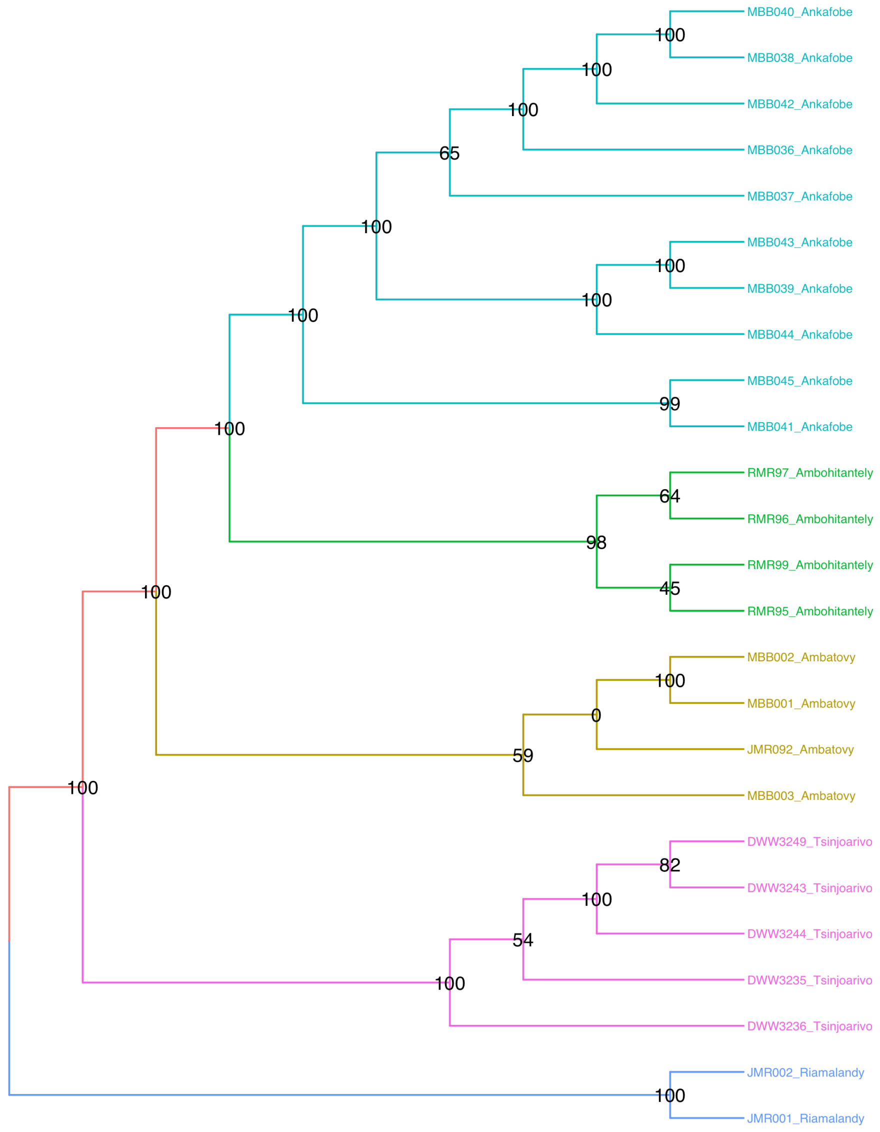

**Figure S7 – Bootstrap Consensus Tree Estimated by SVDquartets with all sites.** 100 bootstrap replicates were performed on a dataset that included all loci with at least 4 taxa per locus. Branches are colored based on sampling location.

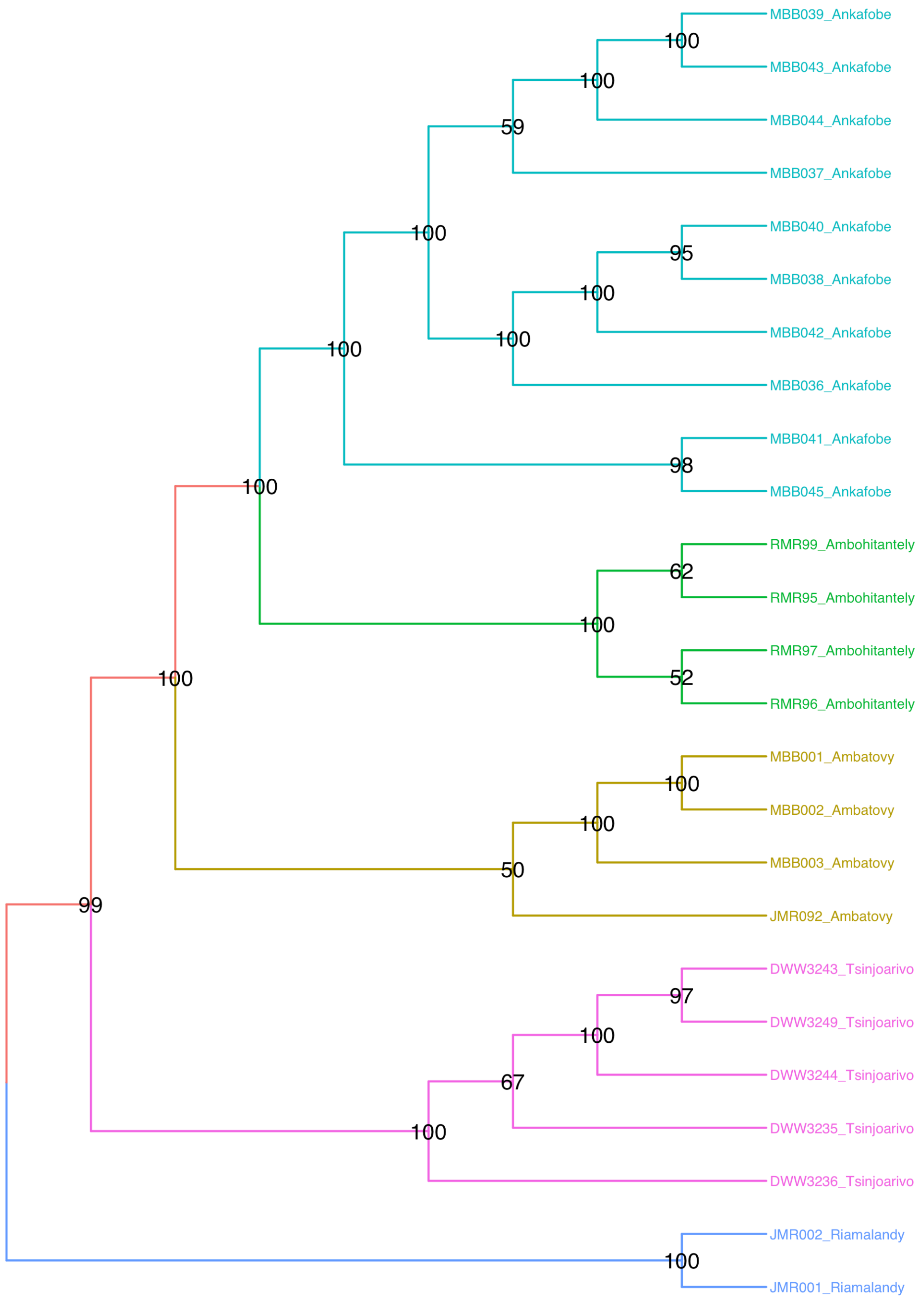

**Figure S8 – Resampled Biallelic Trees with SVDquartets.** Bootstrap consensus trees for 10 datasets that randomly sampled 1 biallelic SNP per locus. Sample 1 is identical to the data used for analyses with SNAPP. Ambatovy is not monophyletic in sample 9.

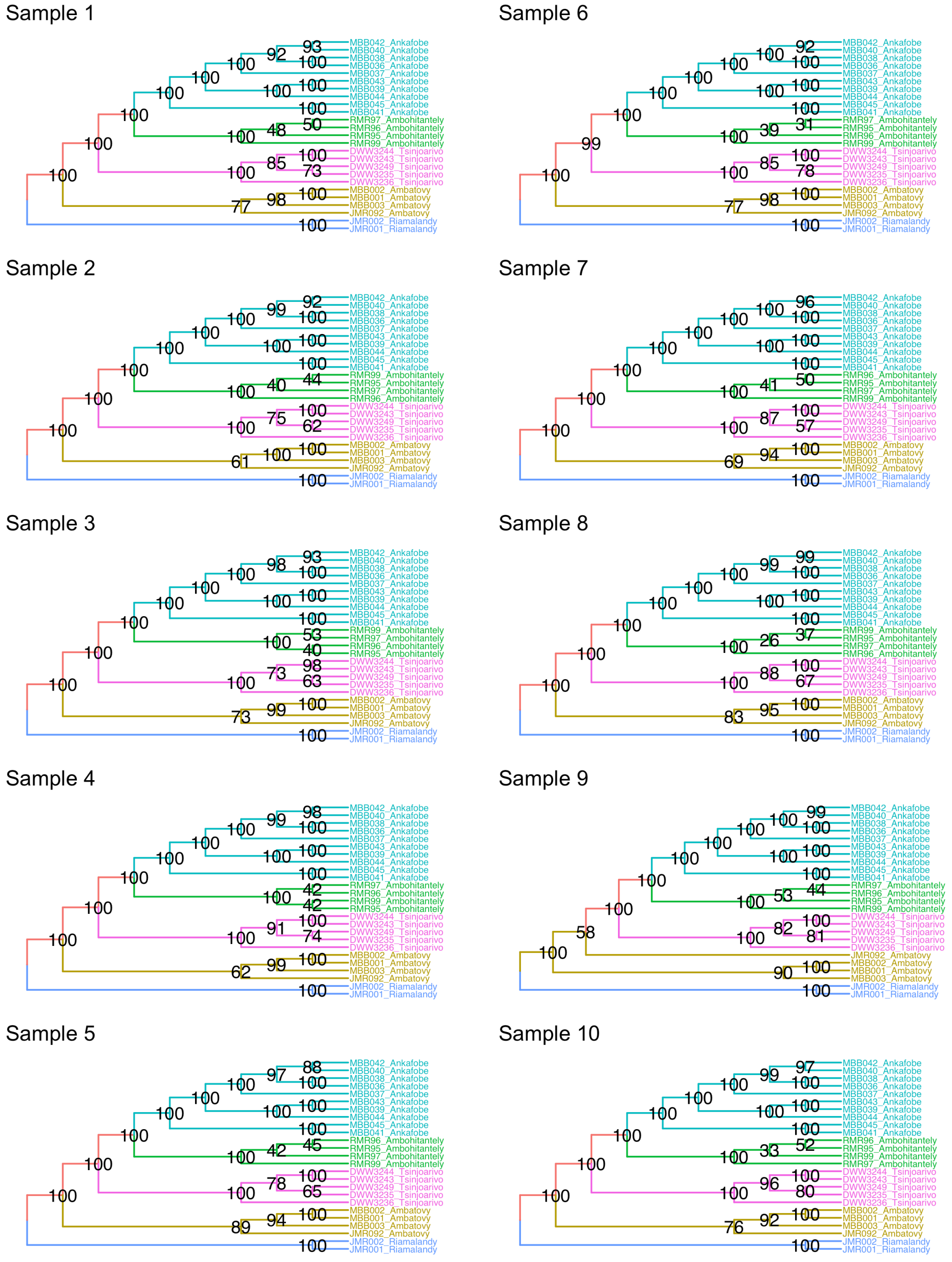

**Figure S9 – Distribution of log likelihoods of structure runs averaged across technical replicates.** The average lnL for 20 datasets of 10000 SNPS is given with their respective *k*. The maximum likelihood for individual and averaged analyses is *k*=3.

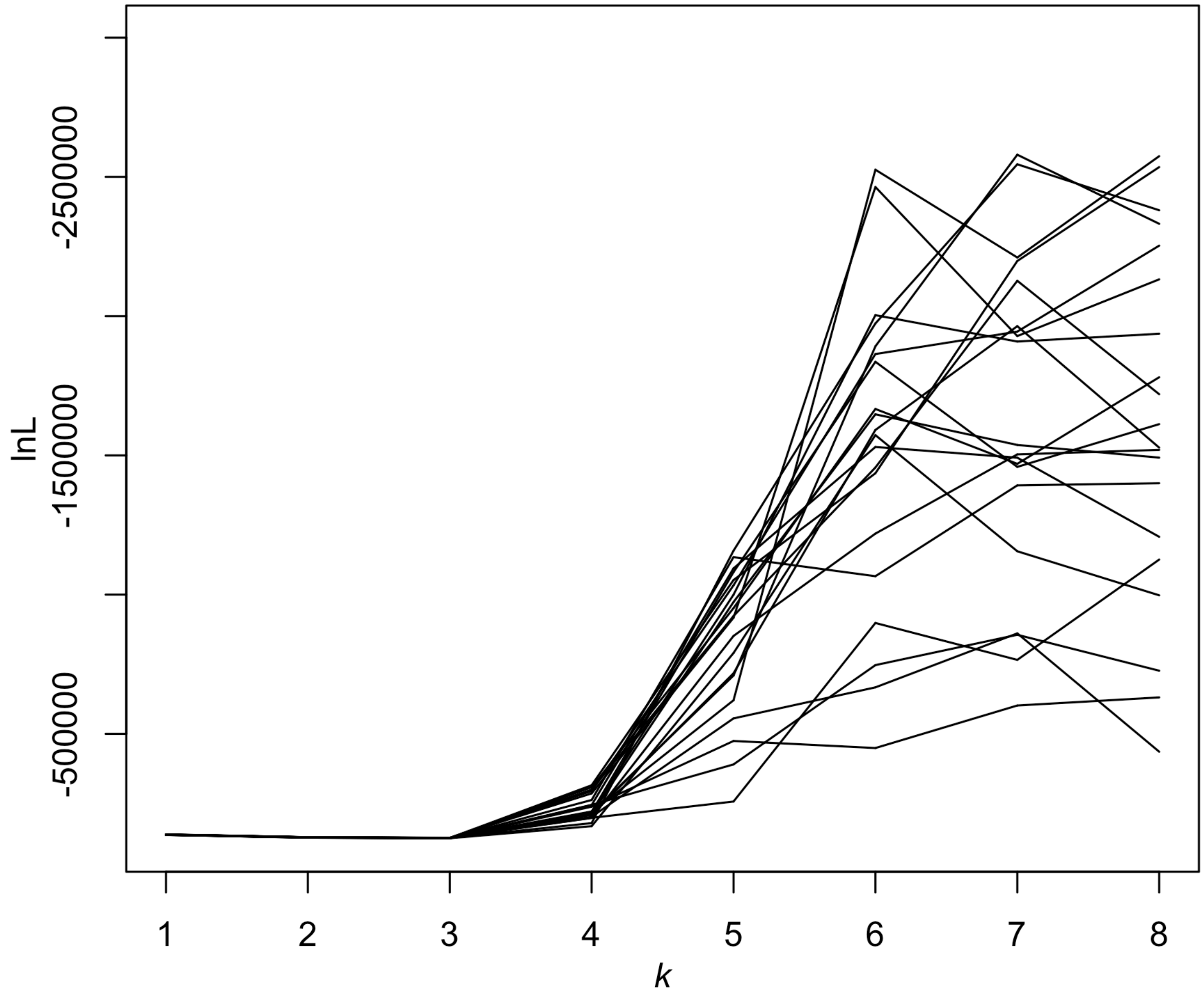

**Figure S10 – Distribution of Δ*k* summary statistics.** Δ*k* measures the rate of change in lnL with respect to increasing *k* clusters. 19 out of 20 of our replicates support an uppers limit of three clusters among *M. lehilahytsara* individuals.

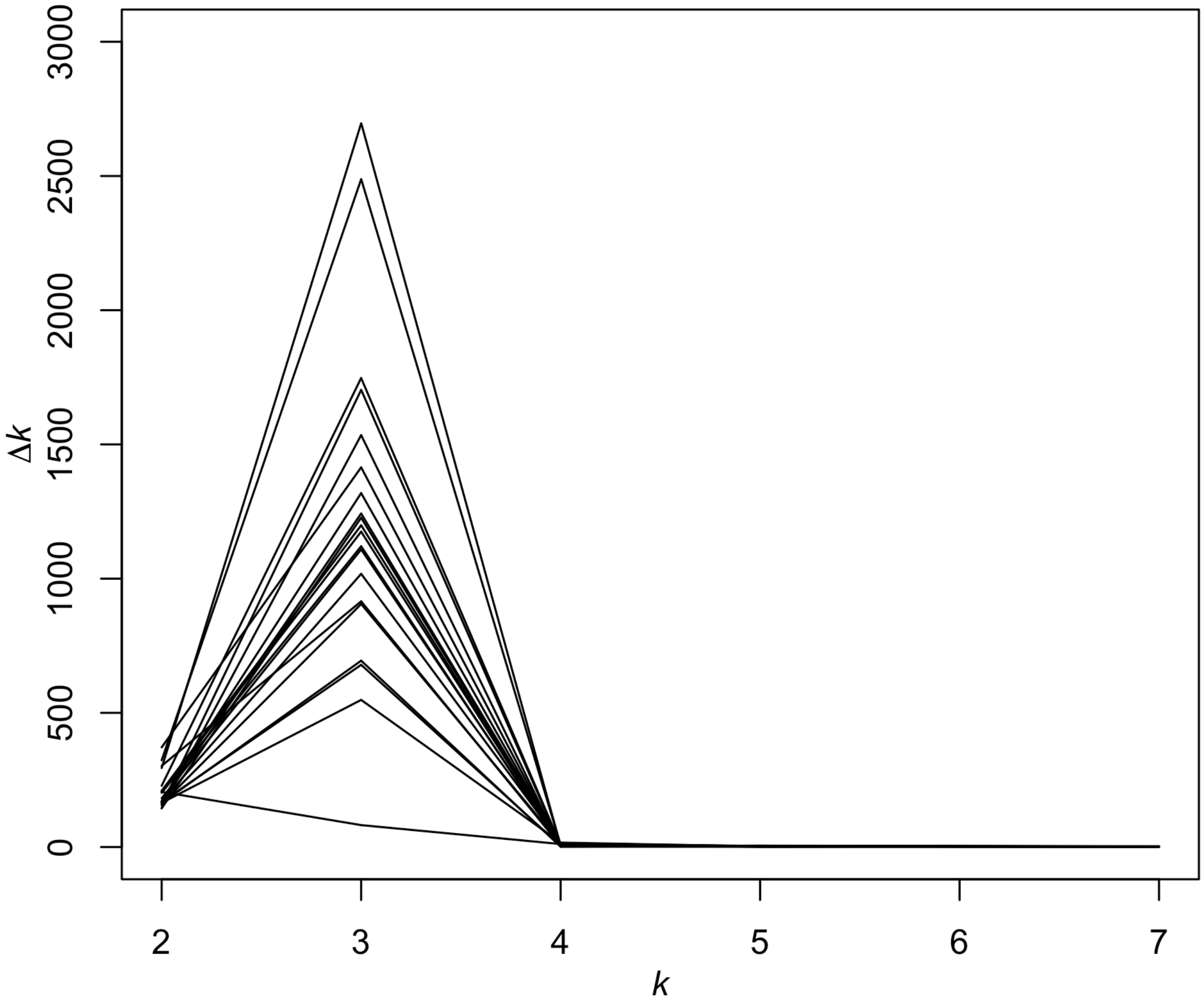

**Figure S11 – Posterior probabilities of cluster assignment for 20 Jackknife replicates.**

One of 10 independent runs is shown for each jackknife replicate of 10000 SNPS. An optimal *k* of 3 was selected for all but one replicate, which had a *k* of 2. There was no variation in *k* across 10 independent runs for each replicate.

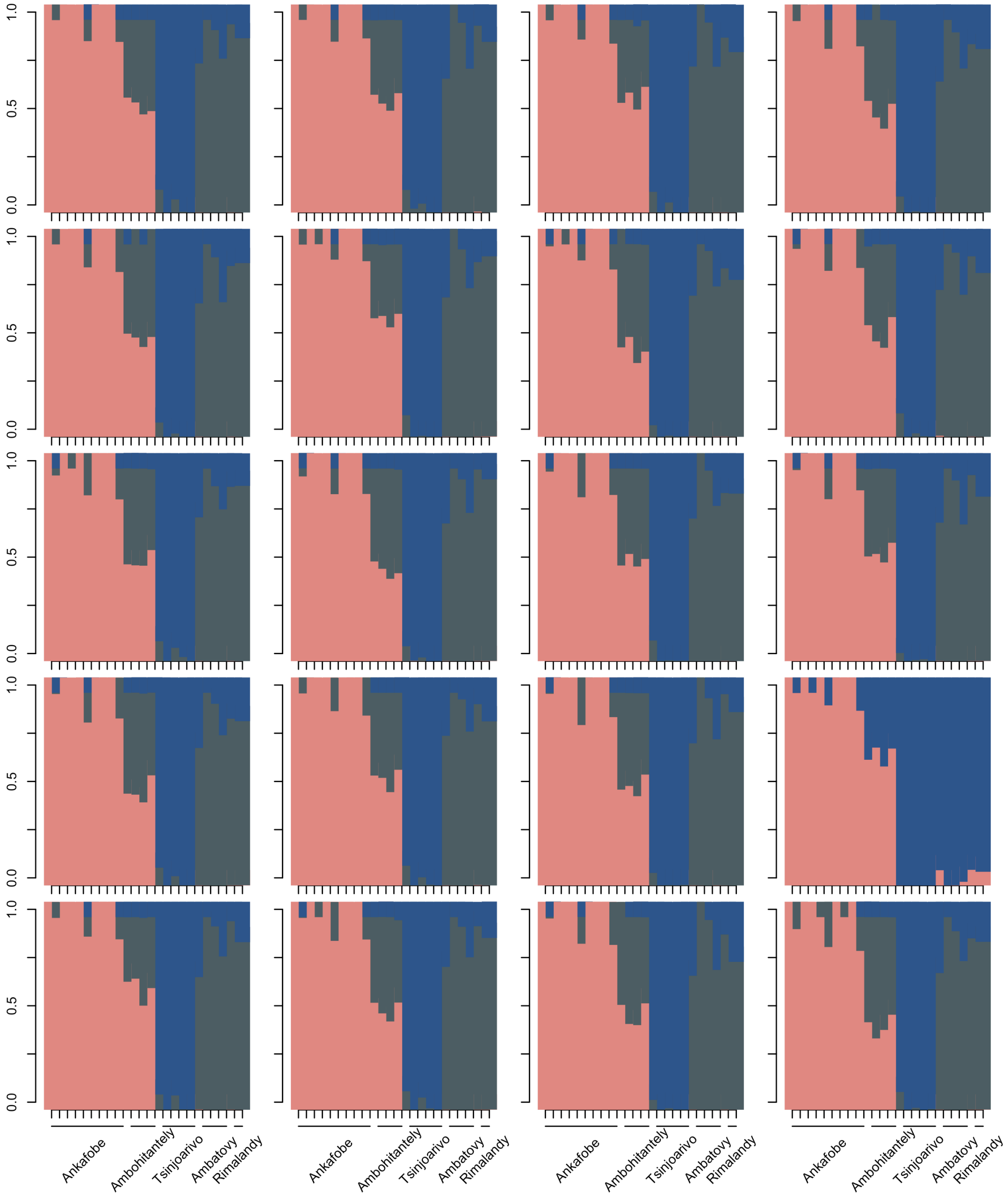

**Figure S12 – Scatterplot of physical distances and $F_{ST}$ between sampling locations.** The regression coefficient (b) and correlation coefficient (r) were calculated with a Mantel test in Arlequin. The p-value was calculated with 9999 permutations and the observed regression would be expected under chance alone. The highest levels of genetic variation observed all involve comparisons between the small 1Km$^2$ forest patch in the Central Highlands (Ankafobe) and sites in the contiguous eastern forests (Riamalandy, Ambatovy, and Tsinjoarivo).

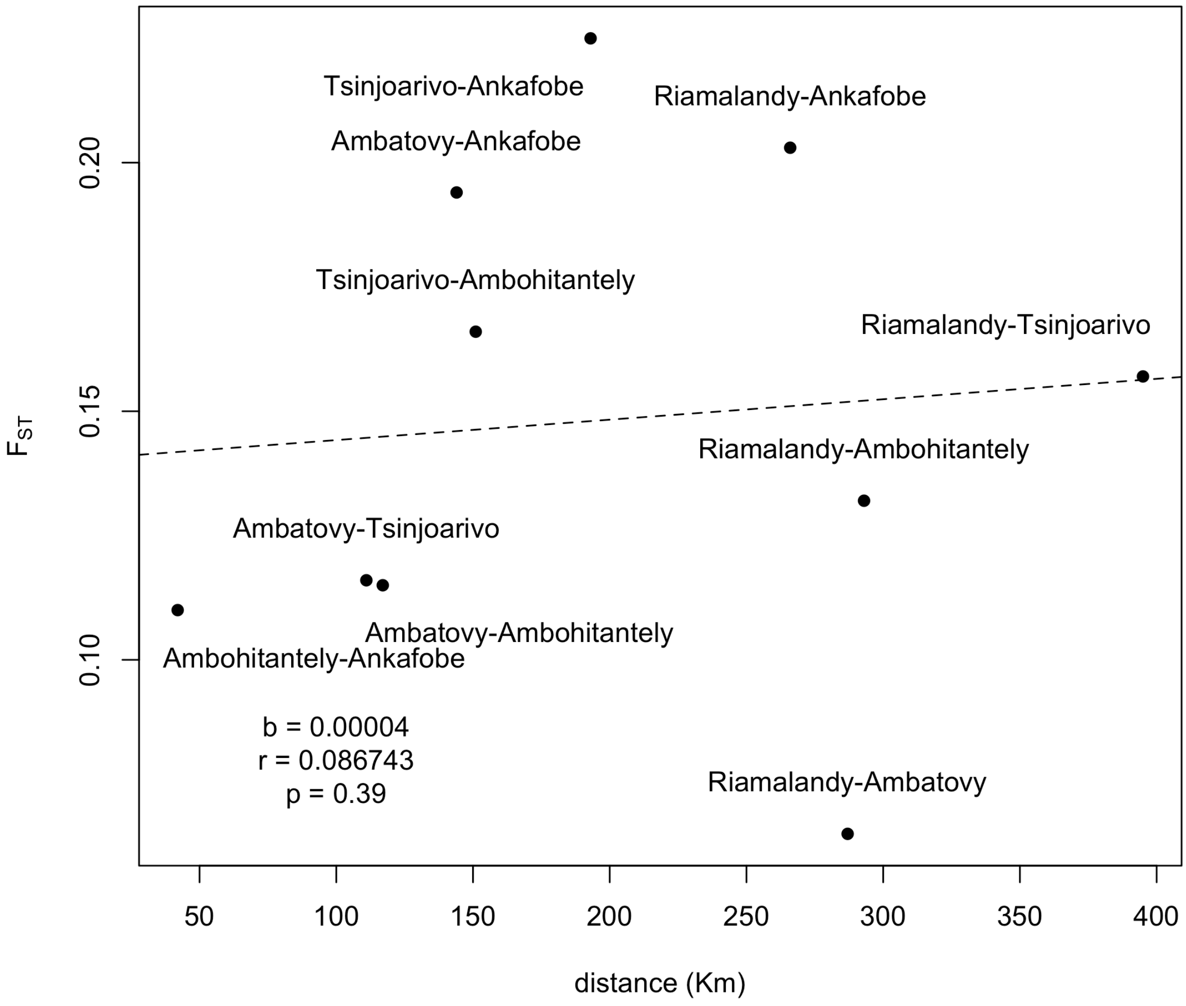

**Figure S13 – Migration models tested for two populations.** a) Ankafobe and Ambohitantely (pink) and Ambatovy, Tsinjoarivo, and Riamalandy are treated as two sperate populations. b) There are likely some parameter identifiability issues, especially for θ estimates which relied on a strong prior for efficient mixing. c) Model 2 is always preferred across replicates.

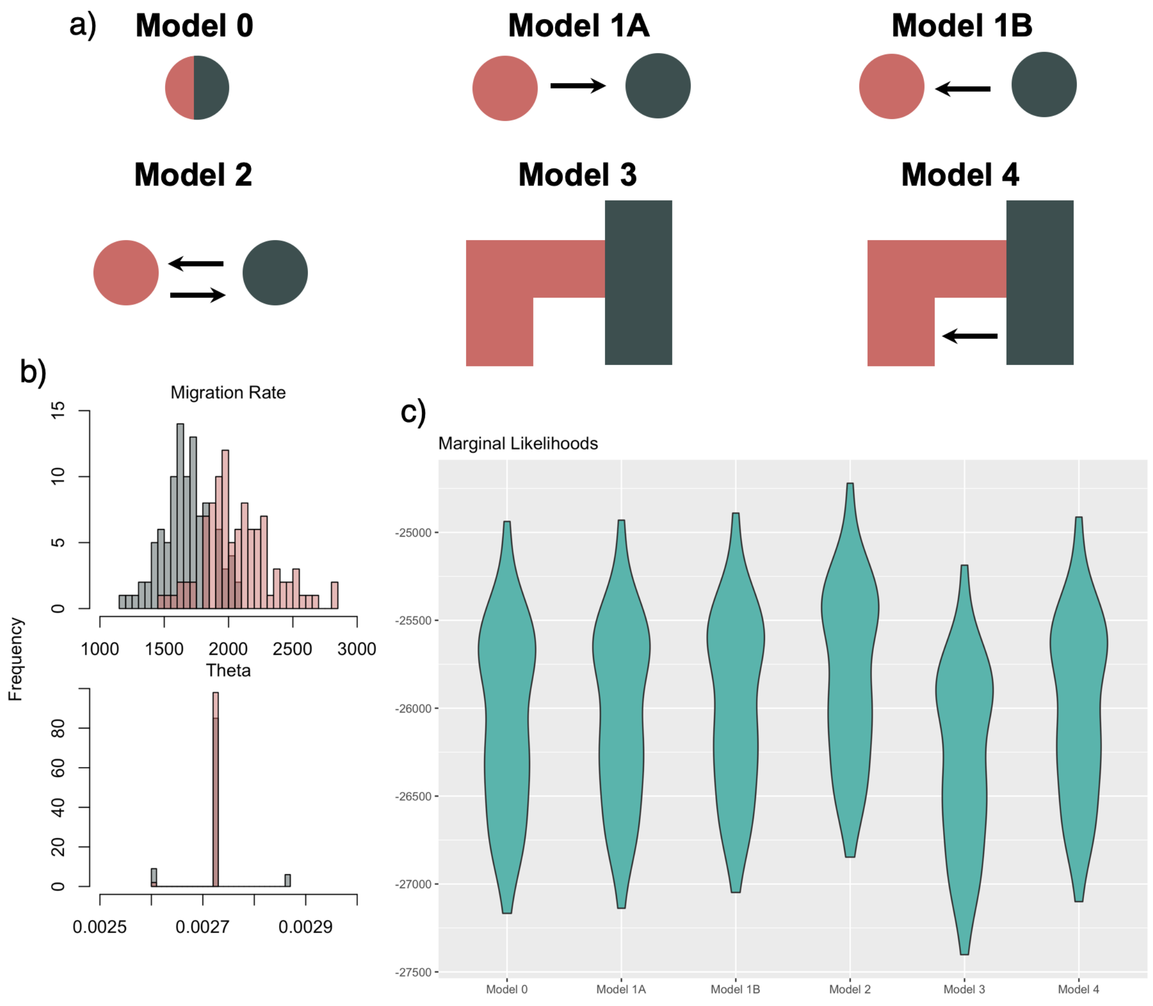

**Figure S14 – Posteriors and Marginal Priors for Ambatovy θ.** Distribution of 40000 posterior samples for the posterior (blue) and marginal prior (red) for a random selection of loci from replicate 1. Loci numbers are displayed above each plot. Samples are based on Model 2.

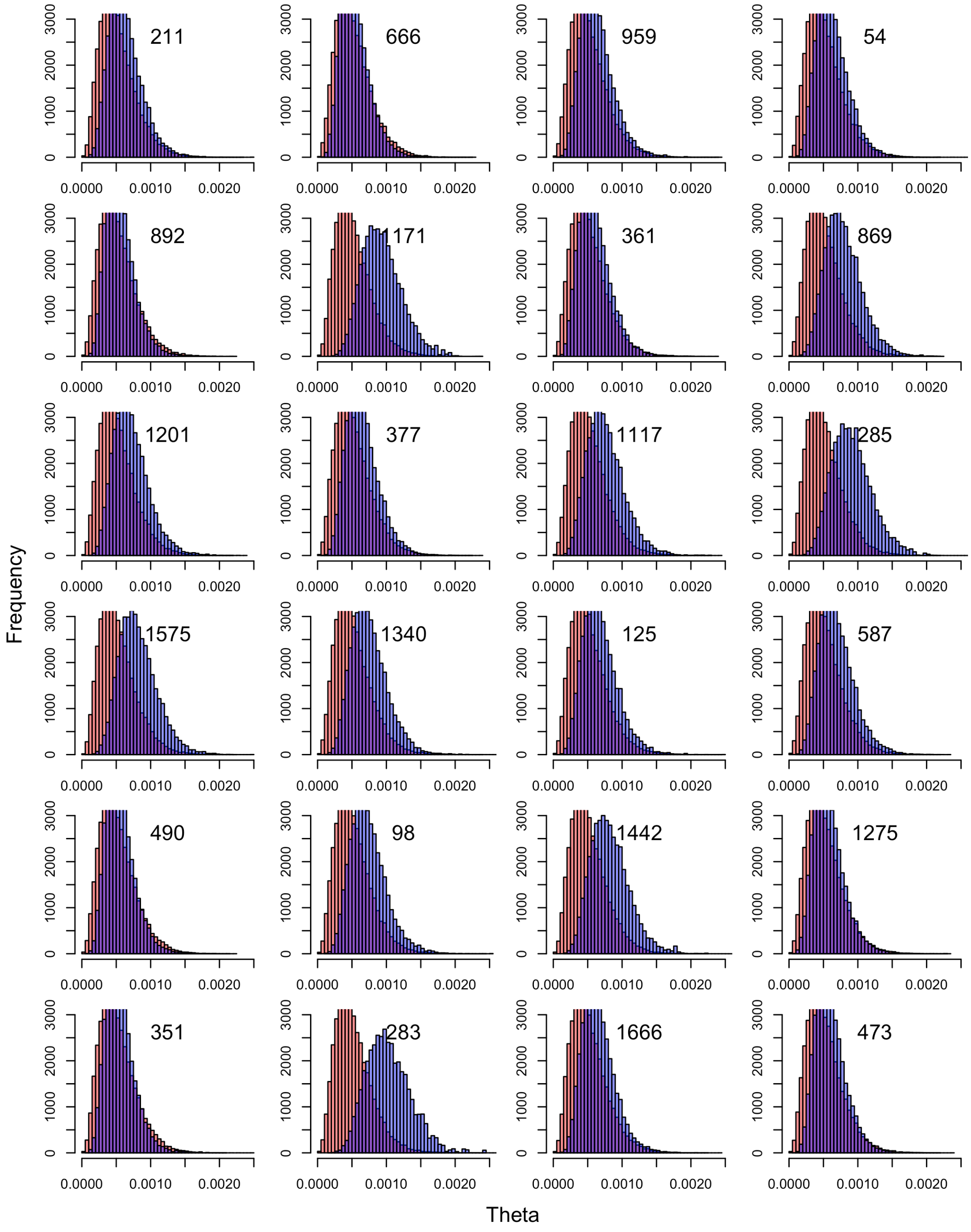

**Figure S15 – Posteriors and Marginal Priors for Ambohitantely θ.** Distribution of 40000 posterior samples for the posterior (blue) and marginal prior (red) for a random selection of loci from replicate 1. Loci numbers are displayed above each plot. Samples are based on Model 2.

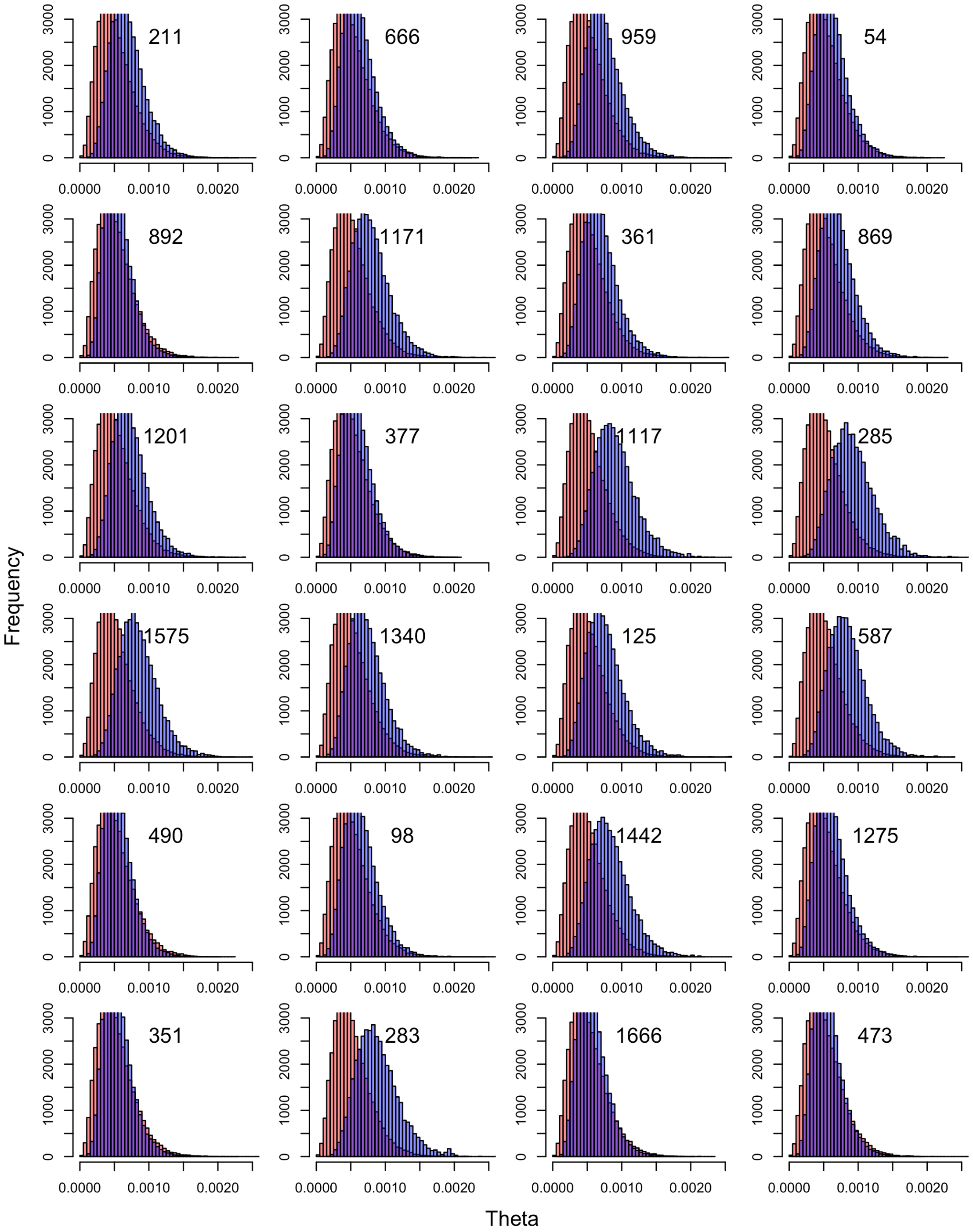

**Figure S16 – Posteriors and Marginal Priors for Ankafobe θ.** Distribution of 40000 posterior samples for the posterior (blue) and marginal prior (red) for a random selection of loci from replicate 1. Loci numbers are displayed above each plot. Samples are based on Model 2.

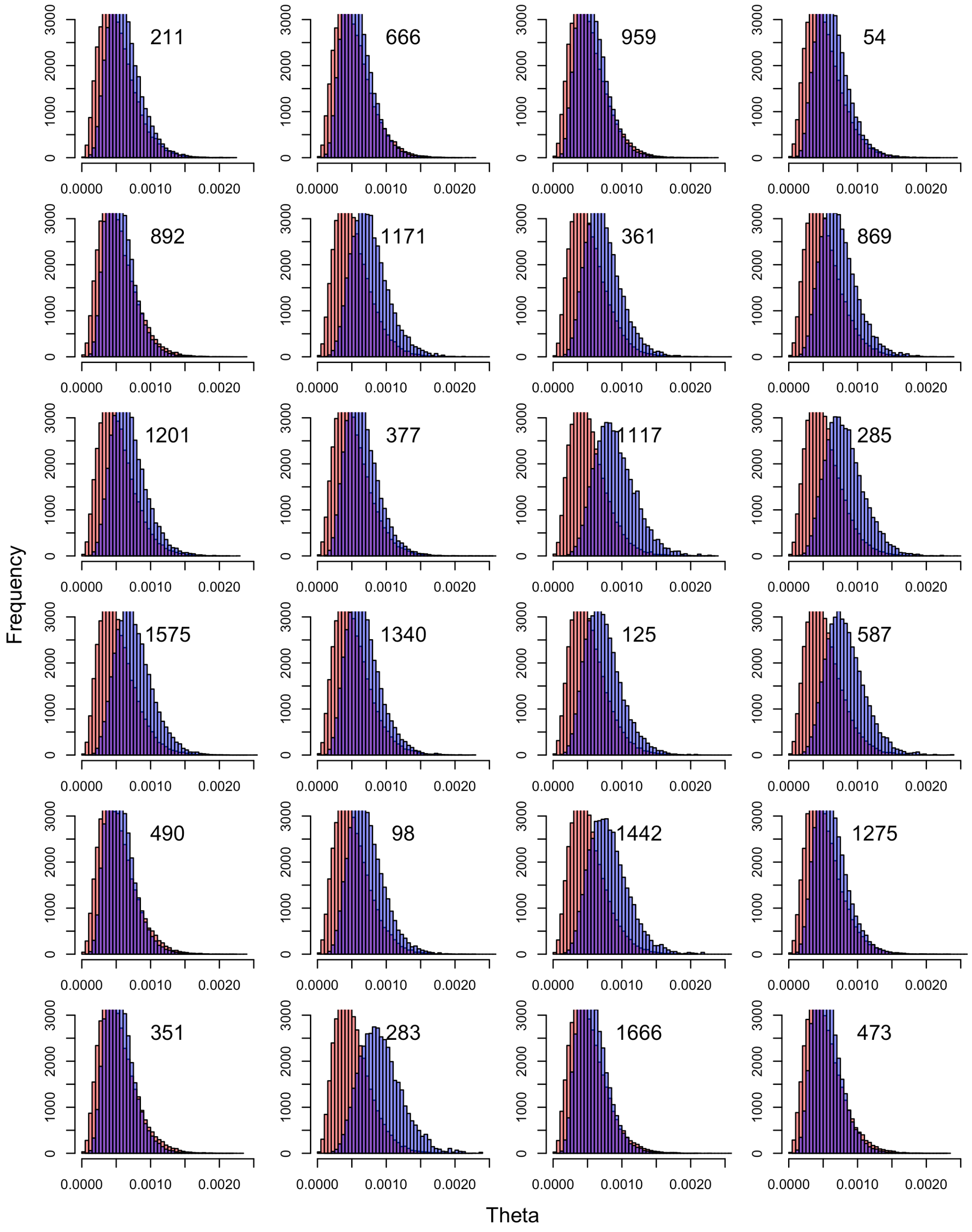

**Figure S17 – Mitochondrial majority rule consensus bootstrap tree for rooting.** Majority rule consensus tree obtained from 500 bootstrap replicates. Bipartitions observed in less than 50% of bootstrap trees were retained if they were compatible with bipartitions already observed in the 50% majority rule consensus tree.

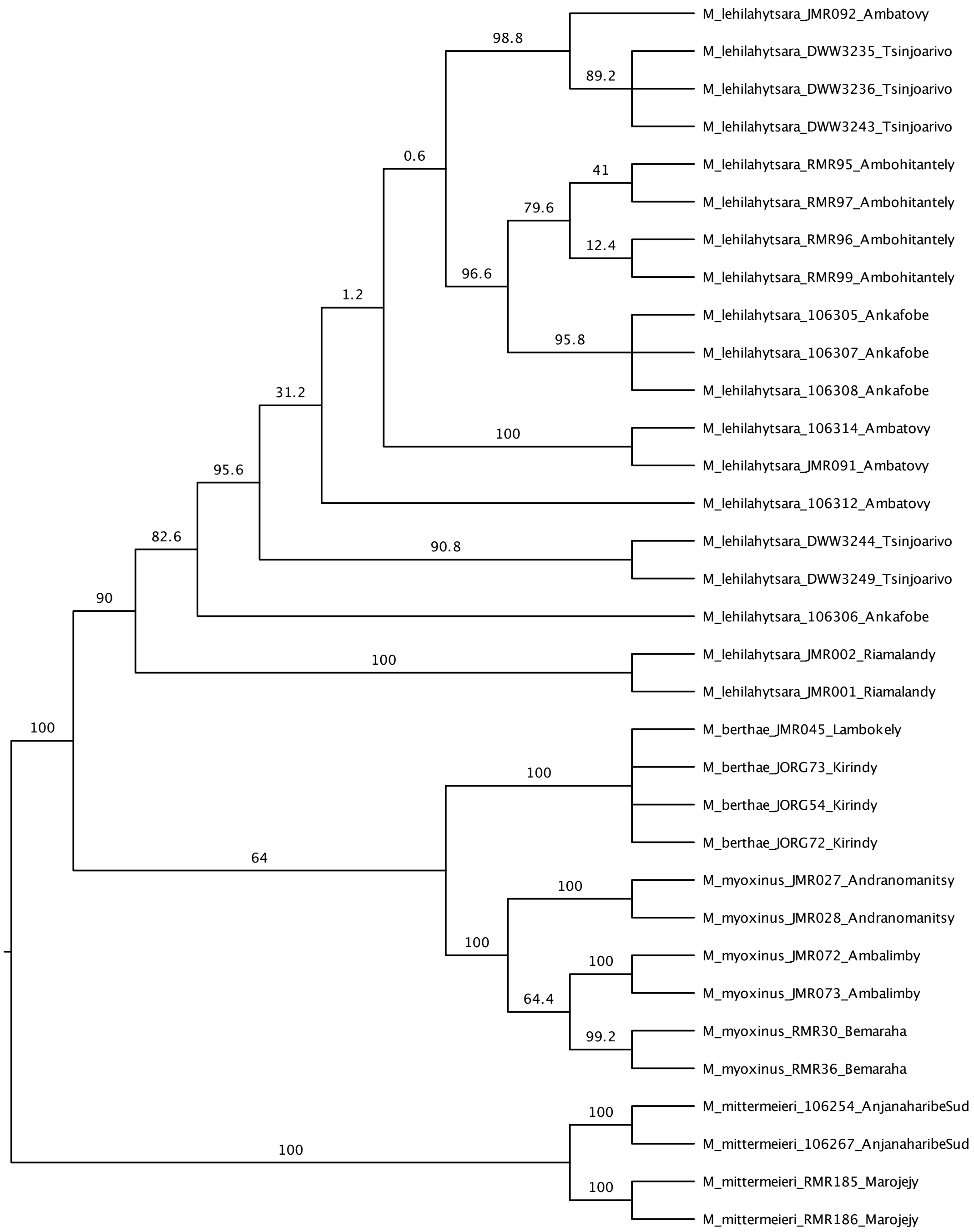

**Figure S18 – Convergence of tau parameters for BPP analysis 00.** Distributions of all thetas are given across 8 independent chains in blue. The marginal prior distribution is shown in yellow.

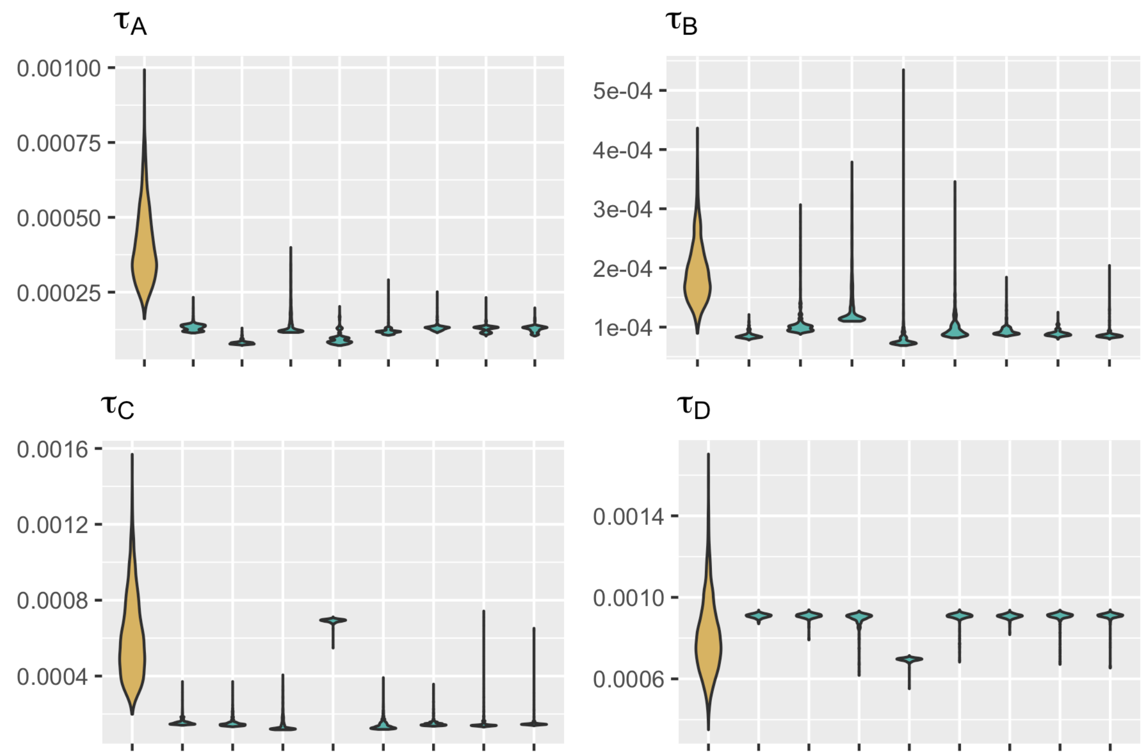

**Figure S19 – Convergence of theta parameters for BPP analysis 00.** Distributions of all thetas are given across 8 independent chains in blue. The marginal prior distribution is shown in yellow.

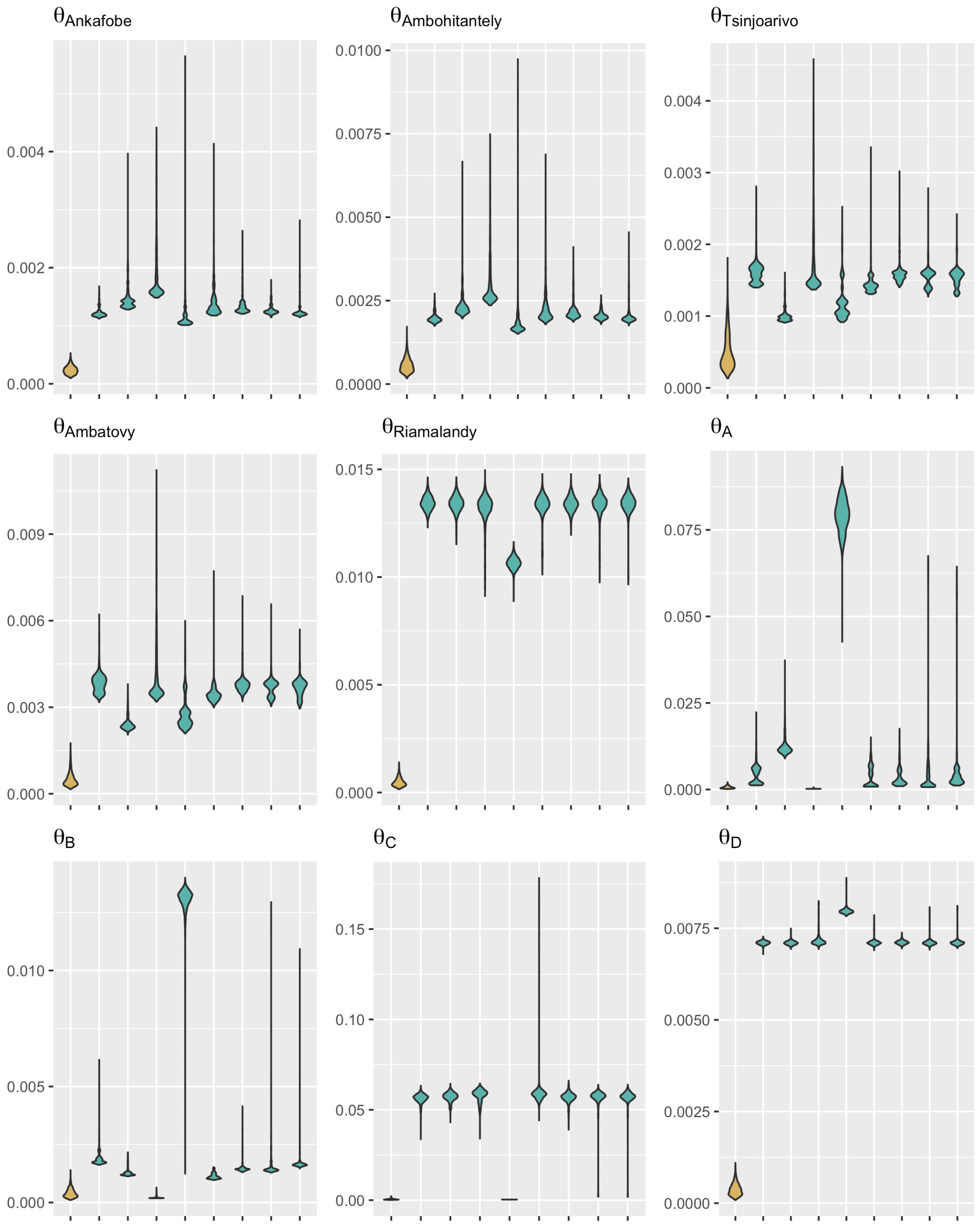

**Figure S20 – Convergence of tau parameters for BPP analysis 01.** Distributions of all thetas are given across 8 independent chains in blue. The marginal prior distribution is shown in yellow.

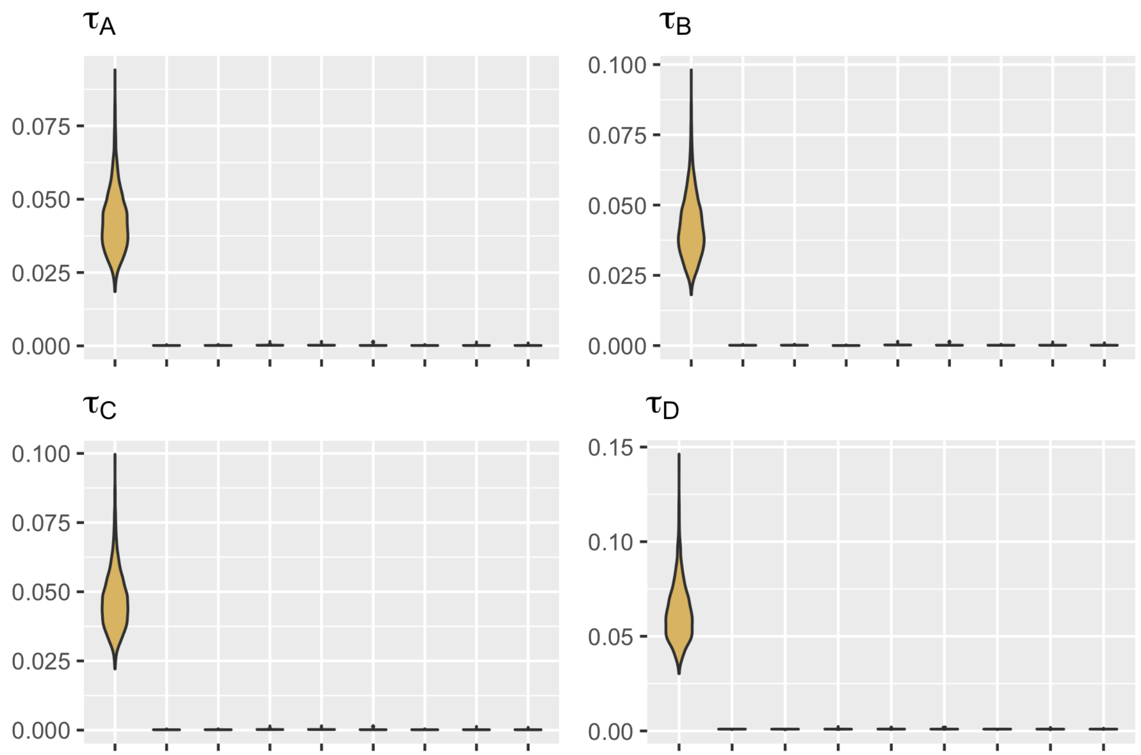

**Figure S21 – Convergence of theta parameters for BPP analysis 01.** Distributions of all thetas are given across 8 independent chains in blue. The marginal prior distribution is shown in yellow.

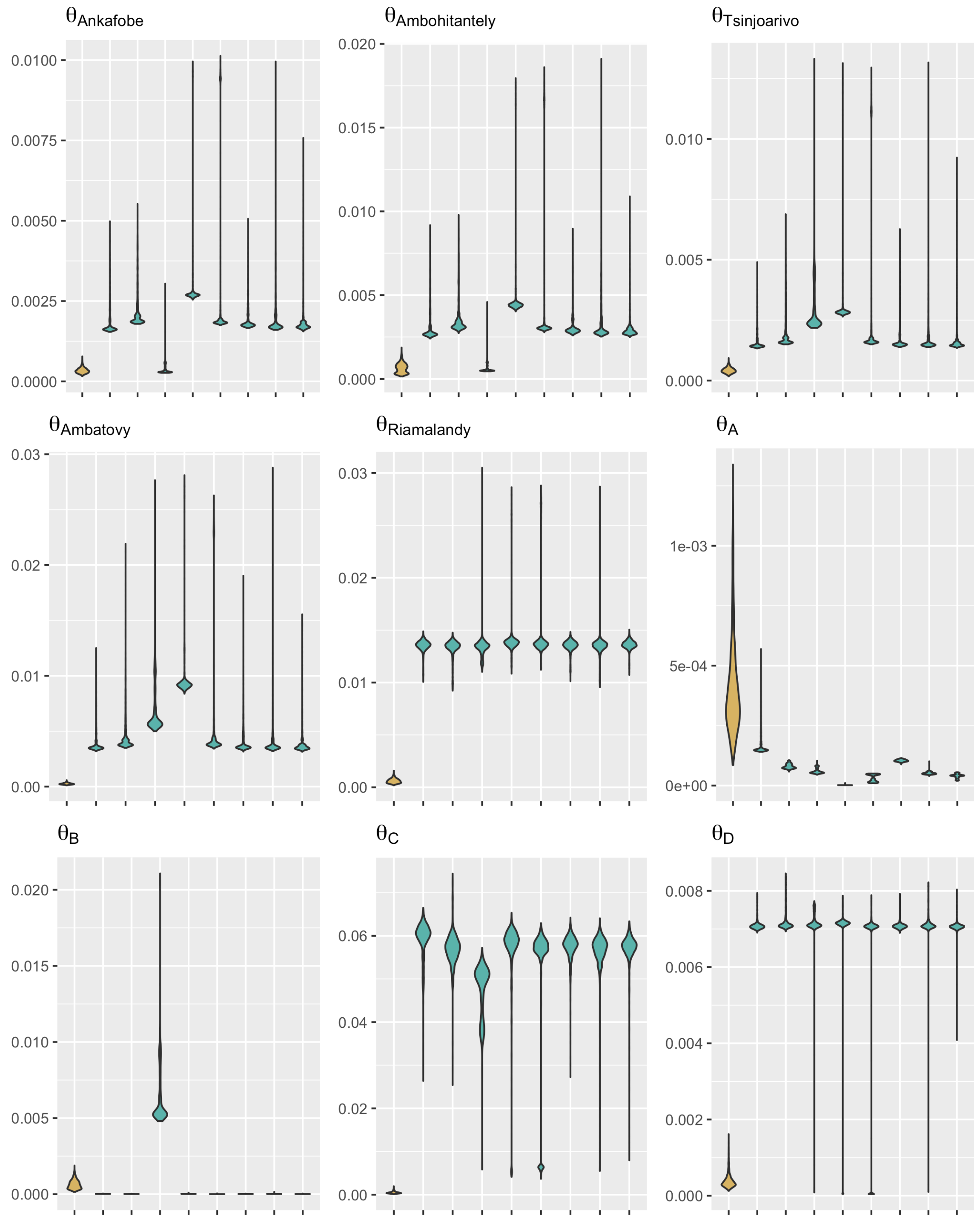

**Figure S22 – Convergence of tau parameters for BPP analysis 10.** Distributions of all thetas are given across 8 independent chains in blue. The marginal prior distribution is shown in yellow.

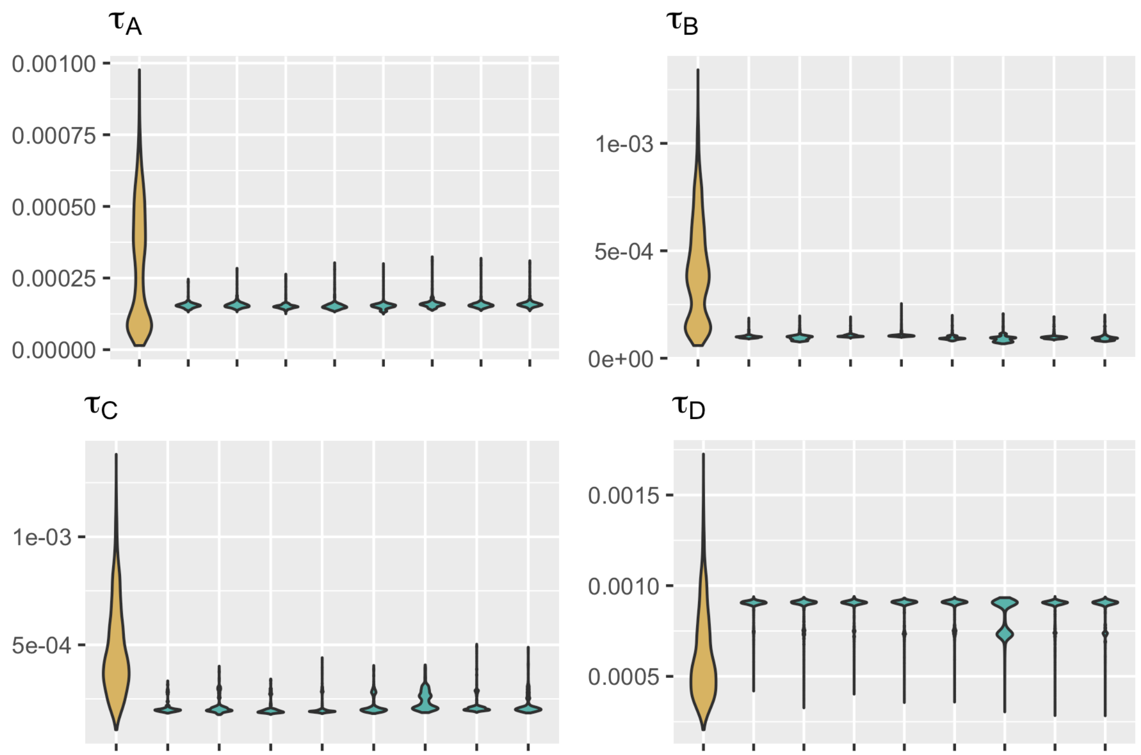

**Figure S23 – Convergence of theta parameters for BPP analysis 10.** Distributions of all thetas are given across 8 independent chains in blue. The marginal prior distribution is shown in yellow.

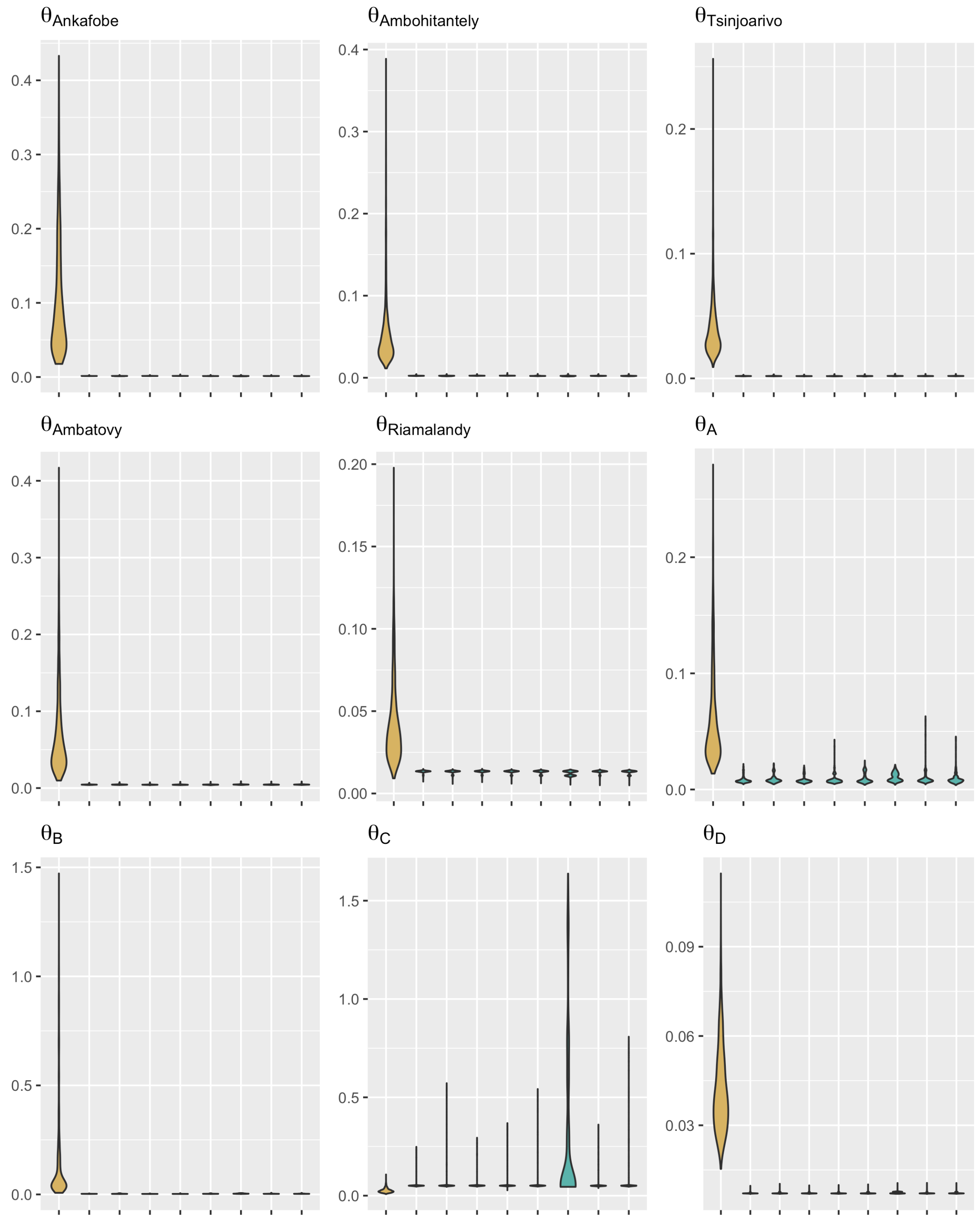

**Figure S24 – Convergence of tau parameters for BPP analysis 11.** Distributions of all thetas are given across 8 independent chains in blue. The marginal prior distribution is shown in yellow.

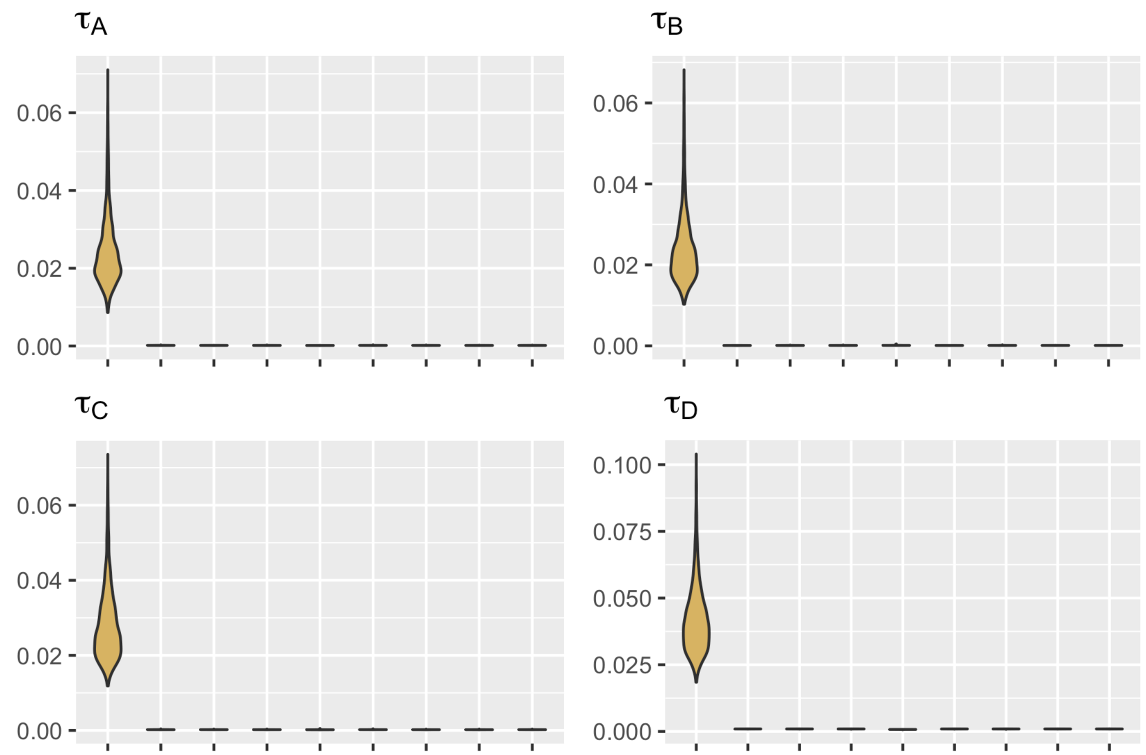

**Figure S25 – Convergence of theta parameters for BPP analysis 11.** Distributions of all thetas are given across 8 independent chains in blue. The marginal prior distribution is shown in yellow.

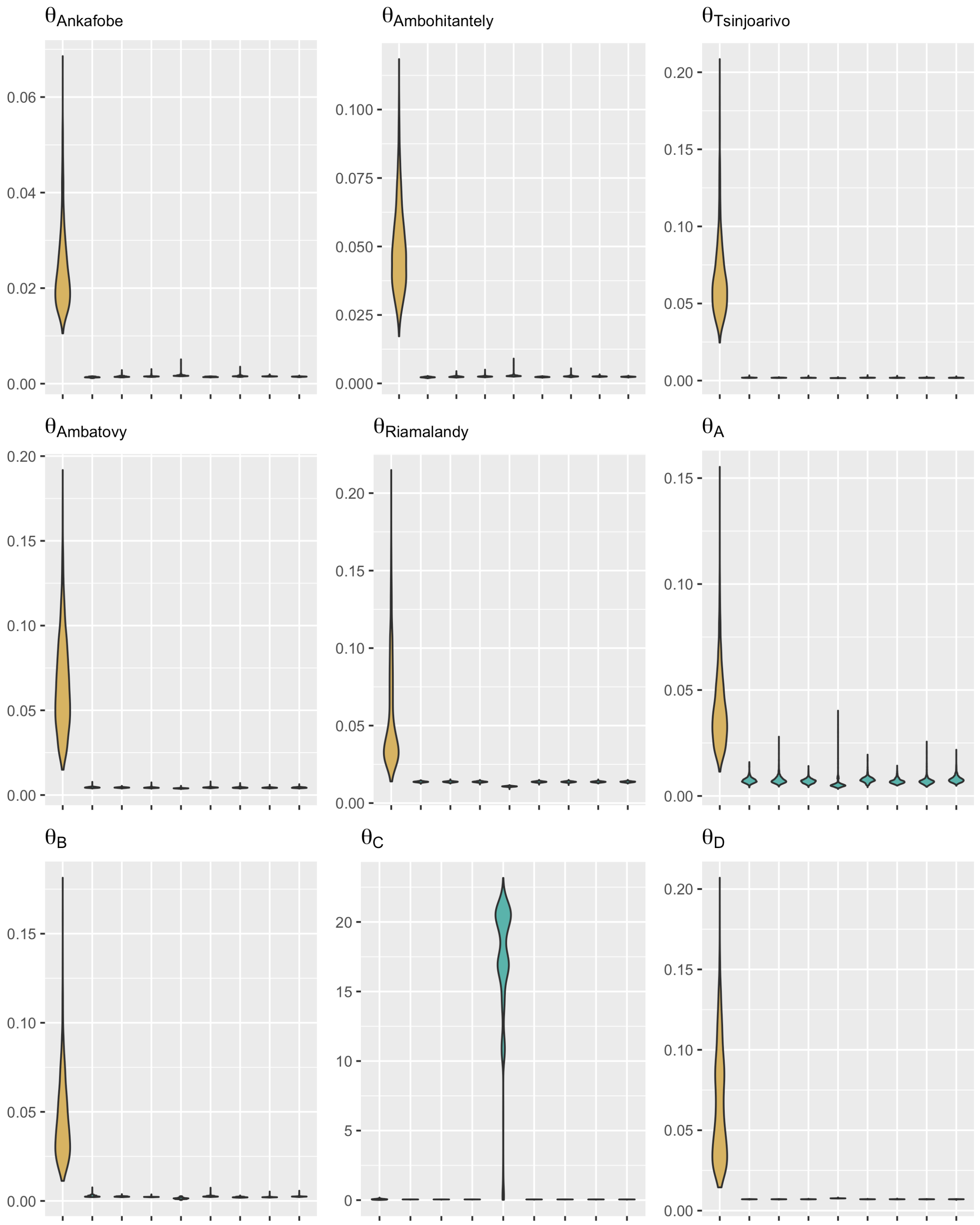

**Figure S26 – EBSP results for all replicates.** Mean $N_e$ estimates over time for 8 replicates of 100 RAD loci for each population. RAD loci are selected at random. Coalescent units were rescaled to absolute time.

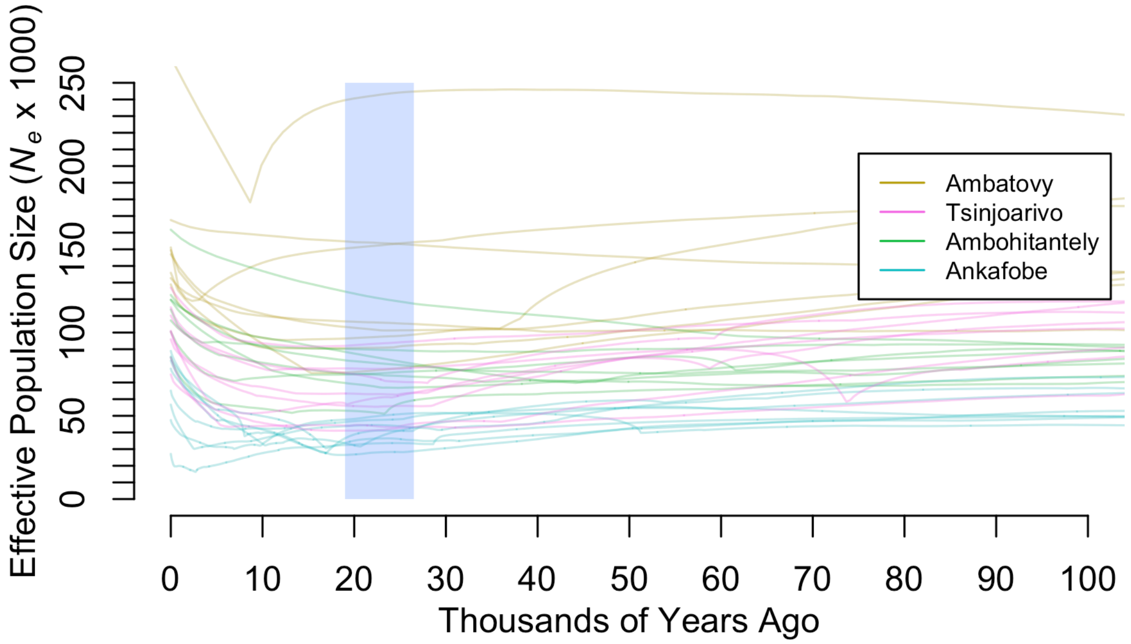

**Figure S27 – Convergence of EBSP results for Ankafobe replicates.** Lines are mean θ estimates and polygons are 95% highest posterior densities.

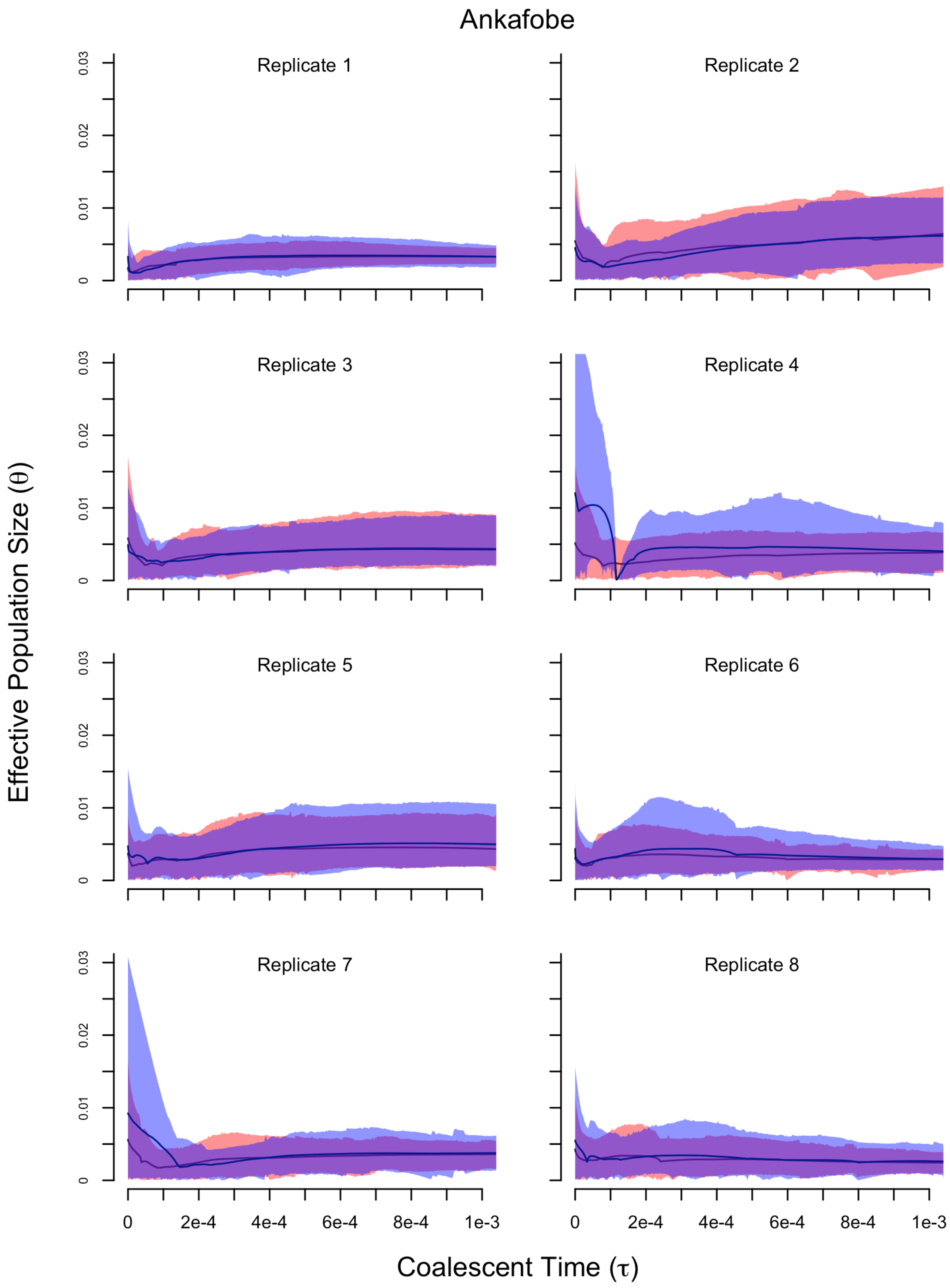

**Figure S28 – Convergence of EBSP results for Ambohitantely replicates.** Lines are mean θ estimates and polygons are 95% highest posterior densities.

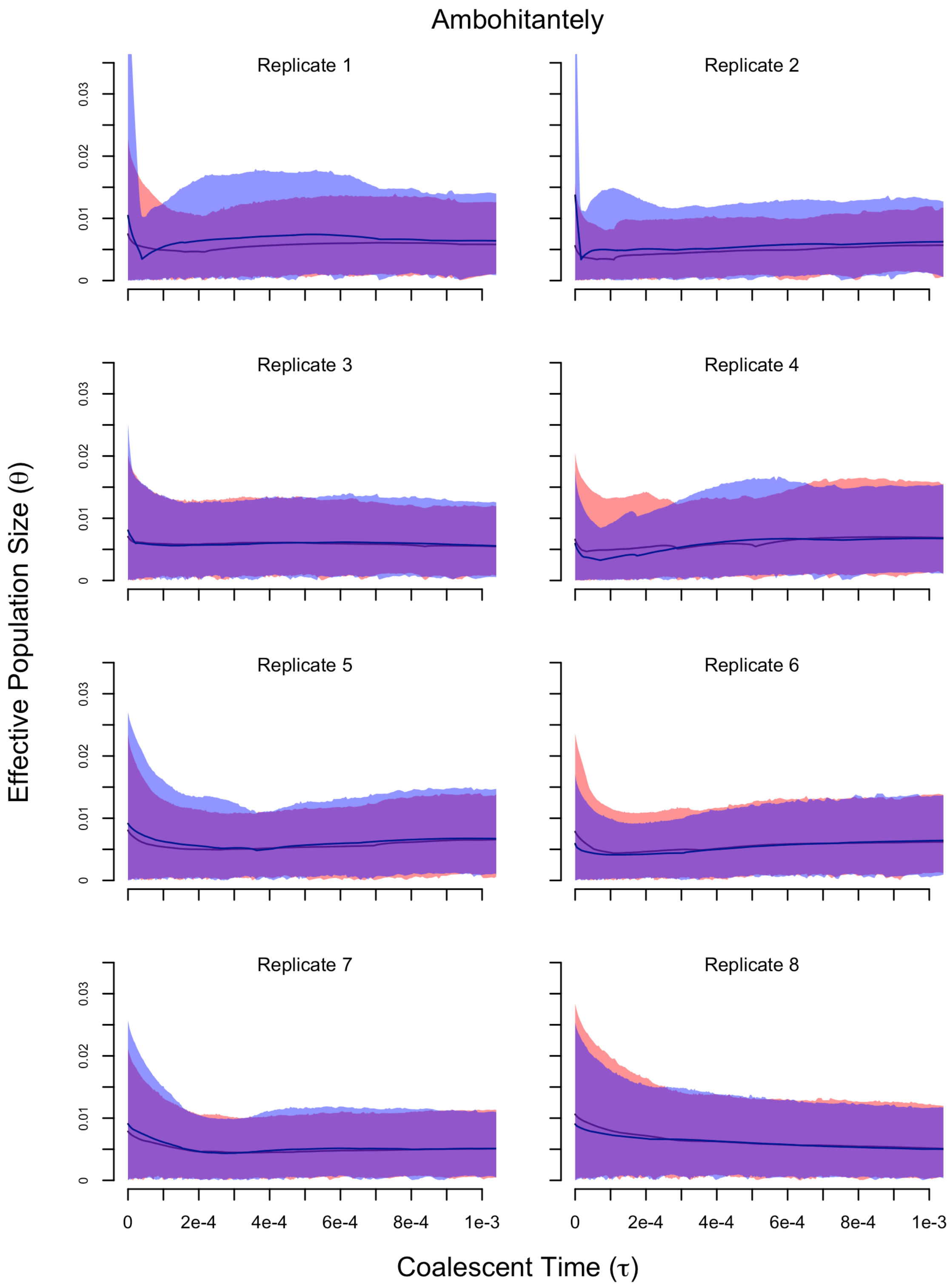

**Figure S29 – Convergence of EBSP results for Tsinjoarivo replicates.** Lines are mean θ estimates and polygons are 95% highest posterior densities.

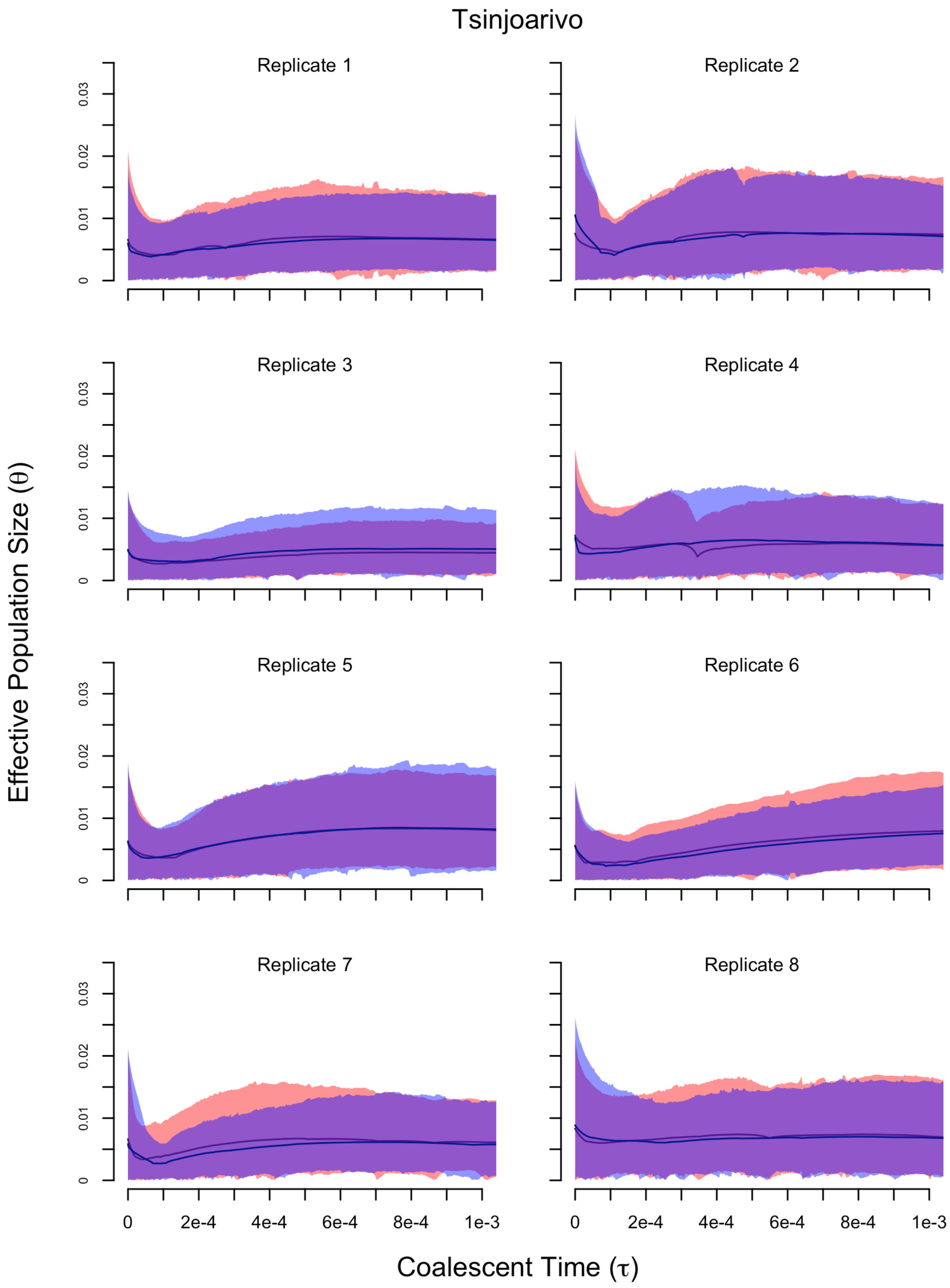

**Figure S30 – Convergence of EBSP results for Ambatovy replicates.** Lines are mean θ estimates and polygons are 95% highest posterior densities.

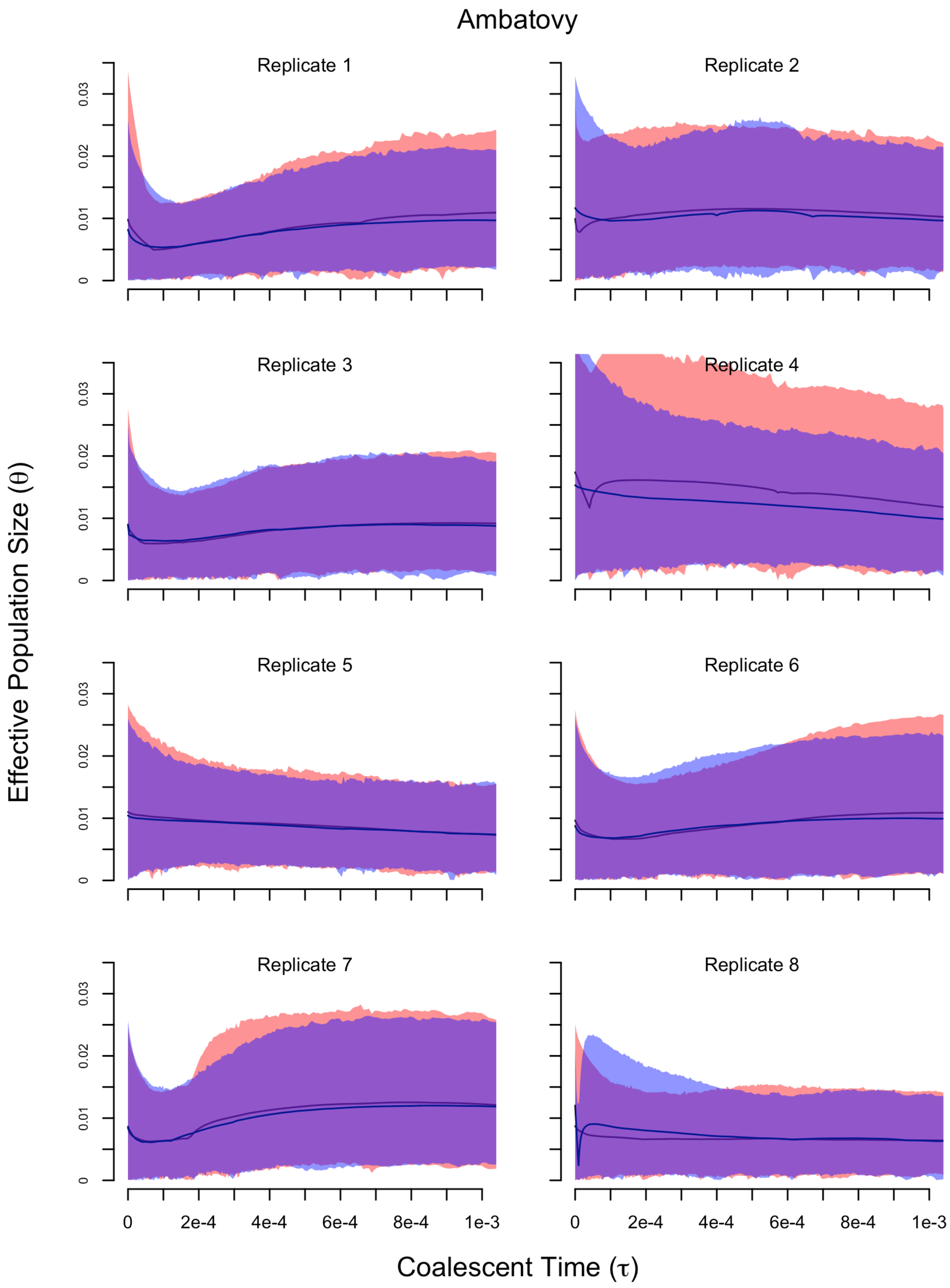

#### Supplementary Tables

**Table S1 – Sequencing results and numbers of recovered loci.** 92889 orthologous loci were recovered amongst *M. lehilahytsara* individuals included in this study. The percent of missing data is relative to these loci.

| Individual | Accession | Site | # of reads | Coverage | Pre-filter loci | Post-filter loci | missing loci | % missing |
|---|---|---|---|---|---|---|---|---|
| mleh001 | JMR092 | Ambatovy | 3756370 | 31.14 | 71238 | 68941 | 23948 | 25.78 |
| mleh002 | MBB001 | Ambatovy | 1456282 | 16.06 | 43647 | 42142 | 50747 | 54.63 |
| mleh003 | MBB002 | Ambatovy | 2998833 | 22.34 | 74595 | 72114 | 20775 | 22.37 |
| mleh004 | MBB003 | Ambatovy | 2461372 | 20.96 | 66347 | 64083 | 28806 | 31.01 |
| mleh005 | RMR95 | Ambohitanely | 2001022 | 17.71 | 42047 | 40590 | 52299 | 56.30 |
| mleh006 | RMR96 | Ambohitanely | 1239822 | 15.16 | 24025 | 23199 | 69690 | 75.03 |
| mleh007 | RMR97 | Ambohitanely | 2158853 | 18.73 | 41342 | 39916 | 52973 | 57.03 |
| mleh008 | RMR99 | Ambohitanely | 1609377 | 19.11 | 29399 | 28379 | 64510 | 69.45 |
| mleh009 | MBB036 | Ankafobe | 911098 | 14.04 | 23844 | 23074 | 69815 | 75.16 |
| mleh010 | MBB037 | Ankafobe | 2530444 | 23.61 | 52645 | 51000 | 41889 | 45.10 |
| mleh011 | MBB038 | Ankafobe | 2681815 | 24.31 | 54896 | 53206 | 39683 | 42.72 |
| mleh012 | MBB039 | Ankafobe | 2244826 | 20.03 | 46357 | 44814 | 48075 | 51.76 |
| mleh013 | MBB040 | Ankafobe | 1527387 | 14.66 | 41879 | 40483 | 52406 | 56.42 |
| mleh014 | MBB041 | Ankafobe | 2431633 | 18.35 | 74039 | 71513 | 21376 | 23.01 |
| mleh015 | MBB042 | Ankafobe | 1821984 | 15.59 | 43885 | 42466 | 50423 | 54.28 |
| mleh016 | MBB043 | Ankafobe | 1467158 | 13.41 | 38805 | 37449 | 55440 | 59.68 |
| mleh017 | MBB044 | Ankafobe | 1179384 | 12.54 | 28420 | 27409 | 65480 | 70.49 |
| mleh018 | MBB045 | Ankafobe | 2593183 | 19.71 | 76767 | 74178 | 18711 | 20.14 |
| mleh019 | JMR001 | Rimalandy | 1696595 | 17.05 | 47571 | 46098 | 46791 | 50.37 |
| mleh020 | JMR002 | Rimalandy | 2598543 | 21.68 | 67974 | 65739 | 27150 | 29.23 |
| mleh021 | DWW3235 | Tsinjoarivo | 2163701 | 16.96 | 62614 | 60491 | 32398 | 34.88 |
| mleh022 | DWW3236 | Tsinjoarivo | 1316210 | 12.89 | 37119 | 35843 | 57046 | 61.41 |
| mleh023 | DWW3243 | Tsinjoarivo | 2527672 | 17.75 | 68409 | 66134 | 26755 | 28.80 |
| mleh024 | DWW3244 | Tsinjoarivo | 2693204 | 19.45 | 66883 | 64646 | 28243 | 30.41 |
| mleh025 | DWW3249 | Tsinjoarivo | 1585086 | 14.00 | 47726 | 46128 | 46761 | 50.34 |

**Table S2 – Mitochondrial sequence data used for rooting.** Genbank accession numbers for sequence data used for mitochondrial phylogeny estimation. Outgroup taxa were included to infer the root of our RADseq phylogeny for BPP analyses.

| Taxon | RADseq ID | CytB | Cox2 |
|---|---|---|---|
| M_lehilahytsara_JMR001_Riamalandy | mleh019 | GU327336 | GU327134 |
| M_lehilahytsara_JMR002_Riamalandy | mleh020 | GU327337 | GU327135 |
| M_lehilahytsara_106312_Ambatovy | mleh002 | KX070723 | KX070709 |
| M_lehilahytsara_106314_Ambatovy | mleh003 | KX070725 | KX070711 |
| M_lehilahytsara_JMR091_Ambatovy | NA | KX070726 | KX070712 |
| M_lehilahytsara_JMR092_Ambatovy | mleh001 | KX070727 | KX070713 |
| M_lehilahytsara_RMR95_Ambohitantely | mleh005 | GU327263 | GU327062 |
| M_lehilahytsara_RMR96_Ambohitantely | mleh006 | GU327264 | GU327063 |
| M_lehilahytsara_RMR97_Ambohitantely | mleh007 | GU327265 | GU327064 |
| M_lehilahytsara_RMR99_Ambohitantely | mleh008 | GU327266 | GU327065 |
| M_lehilahytsara_106305_Ankafobe | mleh009 | KX070736 | KX070700 |
| M_lehilahytsara_106306_Ankafobe | mleh010 | KX070737 | KX070701 |
| M_lehilahytsara_106307_Ankafobe | mleh011 | KX070738 | KX070702 |
| M_lehilahytsara_106308_Ankafobe | mleh012 | KX070739 | KX070703 |
| M_lehilahytsara_DWW3235_Tsinjoarivo | mleh021 | MN939641 | MN939636 |
| M_lehilahytsara_DWW3236_Tsinjoarivo | mleh022 | MN939642 | MN939637 |
| M_lehilahytsara_DWW3243_Tsinjoarivo | mleh023 | MN939643 | MN939638 |
| M_lehilahytsara_DWW3244_Tsinjoarivo | mleh024 | MN939644 | MN939639 |
| M_lehilahytsara_DWW3249_Tsinjoarivo | mleh025 | MN939635 | MN939640 |
| M_myoxinus_JMR027_Andranomanitsy | NA | GU327354 | GU327152 |
| M_myoxinus_JMR028_Andranomanitsy | NA | GU327355 | GU327153 |
| M_myoxinus_JMR072_Ambalimby | NA | GU327357 | GU327155 |
| M_myoxinus_JMR073_Ambalimby | NA | GU327358 | GU327156 |
| M_myoxinus_RMR30_Bemaraha | NA | GU327201 | GU327015 |
| M_myoxinus_RMR36_Bemaraha | NA | GU327202 | GU327016 |
| M_mittermeieri_RMR185_Marojejy | NA | GU327312 | GU327111 |
| M_mittermeieri_RMR186_Marojejy | NA | GU327313 | GU327112 |
| M_mittermeieri_106254_AnjanaharibeSud | NA | KX070741 | KX070705 |
| M_mittermeieri_106267_AnjanaharibeSud | NA | KX070742 | KX070706 |
| M_berthae_JMR045_Lambokely | NA | GU327356 | GU327154 |
| M_berthae_JORG73_Kirindy | NA | GU327161 | GU326974 |
| M_berthae_JORG54_Kirindy | NA | GU327162 | GU326975 |
| M_berthae_JORG72_Kirindy | NA | GU327163 | GU326976 |

**Table S3 – Pairwise $F_{ST}$ between *M. lehilahytsara* sampling locations.** The lower triangular is $F_{ST}$ values and the diagonal is $F_{IS}$ values . Interpreting $F_{ST}$ values between Riamalandy and other sites may be tenuous due to the limited sampling of two individuals at that site.

| | **Riamalandy** | **Ambatovy** | **Tsinjoarivo** | **Ambohitantely** | **Ankafobe** |
|---|---|---|---|---|---|
| **Riamalandy** | -0.098 | | | | |
| **Ambatovy** | 0.065 | -0.184 | | | |
| **Tsinjoarivo** | 0.157 | 0.116 | -0.141 | | |
| **Ambohitantely** | 0.132 | 0.115 | 0.166 | -0.192 | |
| **Ankafobe** | 0.203 | 0.194 | 0.225 | 0.11 | -0.217 |

**Table S4 – AMOVA analysis of SNP data.** Sampling locations were grouped into the CHS (Ambohitantely and Ankafobe) and eastern forest (Riamalandy, Ambatovy, and Tsinjoarivo). The p-values are based on 9999 permutations.

| Source of Variation | df | Sum of Squares | Variance Components | Percentage of Variation | Fixation Index | p-value |
|---|---|---|---|---|---|---|
| Between CHS and eastern forest | 1 | 2256.71 | 41.64 | 7.24 | $F_{CT}$ = 0.072 | 0.098 ± 0.003 |
| Between populations in CHS or eastern forest | 3 | 3027.01 | 64.1 | 11.14 | $F_{SC}$ = 0.120 | < 1e-6 |
| Within Populations | 45 | 21132.07 | 469.6 | 81.62 | $F_{ST}$ = 0.184 | < 1e-6 |
| Total | 49 | 26415.79 | 575.34 | | | |

**Table S5 – Model selection results for two population MIGRATE analyses.** Model probabilities for each jackknife replicate of 100 RAD loci are given. The model that includes migration both to and from the east and west without divergence is preferred in all replicates.

| Jackknife Replicate | Model 0 | Model 1A | Model1B | Model 2 | Model 3 | Model |
|---|---|---|---|---|---|---|
| 0 | 5.30E-96 | 1.98E-86 | 4.71E-77 | 1 | 6.75E-205 | 2.24E-80 |
| 1 | 6.87E-111 | 2.19E-106 | 4.99E-84 | 1 | 5.06E-221 | 5.69E-96 |
| 2 | 1.78E-126 | 3.10E-109 | 5.61E-75 | 1 | 5.52E-220 | 5.19E-99 |
| 3 | 3.82E-108 | 4.01E-96 | 7.78E-74 | 1 | 1.97E-212 | 1.12E-97 |
| 4 | 7.80E-133 | 1.15E-105 | 4.13E-81 | 1 | 2.24E-211 | 5.59E-93 |
| 5 | 2.42E-114 | 4.97E-98 | 1.40E-76 | 1 | 1.06E-206 | 2.15E-84 |
| 6 | 5.01E-124 | 3.92E-91 | 8.74E-69 | 1 | 7.53E-193 | 4.08E-76 |
| 7 | 8.86E-109 | 6.59E-103 | 6.75E-77 | 1 | 9.77E-224 | 4.83E-91 |
| 8 | 4.17E-142 | 1.02E-102 | 2.43E-80 | 1 | 2.73E-210 | 7.12E-90 |
| 9 | 2.00E-94 | 3.29E-105 | 2.49E-79 | 1 | 4.79E-223 | 1.82E-100 |
| 10 | 7.47E-101 | 2.59E-97 | 7.23E-69 | 1 | 2.38E-223 | 8.89E-91 |
| 11 | 1.48E-141 | 2.86E-128 | 6.42E-85 | 1 | 1.16E-238 | 5.40E-111 |
| 12 | 5.62E-134 | 7.76E-104 | 1.31E-82 | 1 | 3.63E-220 | 1.14E-91 |
| 13 | 1.55E-128 | 7.30E-88 | 4.46E-79 | 1 | 7.21E-188 | 3.05E-76 |
| 14 | 6.28E-115 | 9.97E-108 | 3.91E-83 | 1 | 7.46E-224 | 3.62E-99 |
| 15 | 2.51E-105 | 4.74E-103 | 6.82E-81 | 1 | 1.82E-221 | 2.10E-94 |
| 16 | 9.14E-90 | 8.54E-87 | 1.98E-71 | 1 | 2.87E-191 | 1.42E-77 |
| 17 | 1.51E-124 | 4.11E-102 | 2.35E-72 | 1 | 4.11E-210 | 1.15E-94 |
| 18 | 2.75E-140 | 4.31E-101 | 8.76E-85 | 1 | 1.41E-195 | 7.15E-76 |
| 19 | 1.12E-144 | 1.96E-106 | 1.78E-81 | 1 | 2.99E-215 | 1.28E-92 |
| 20 | 1.00E-93 | 1.03E-93 | 1.48E-70 | 1 | 6.74E-204 | 1.05E-87 |
| 21 | 4.72E-97 | 7.47E-97 | 4.81E-70 | 1 | 9.80E-206 | 2.48E-88 |
| 22 | 1.75E-117 | 2.66E-112 | 7.92E-77 | 1 | 1.98E-237 | 2.22E-99 |
| 23 | 1.08E-114 | 1.25E-101 | 3.94E-82 | 1 | 1.56E-199 | 4.79E-88 |
| 24 | 1.60E-104 | 2.07E-100 | 1.19E-78 | 1 | 4.86E-213 | 3.46E-93 |
| 25 | 1.95E-123 | 3.38E-96 | 5.68E-88 | 1 | 3.22E-216 | 5.56E-87 |
| 26 | 1.57E-98 | 1.19E-94 | 2.01E-77 | 1 | 3.26E-206 | 1.09E-84 |
| 27 | 6.78E-133 | 7.93E-93 | 4.23E-74 | 1 | 5.61E-199 | 1.46E-78 |
| 28 | 3.01E-94 | 6.36E-97 | 6.80E-80 | 1 | 8.50E-209 | 1.55E-91 |
| 29 | 1.86E-117 | 6.09E-107 | 4.54E-86 | 1 | 3.44E-227 | 4.52E-92 |
| 30 | 3.86E-120 | 4.20E-103 | 8.45E-75 | 1 | 8.27E-214 | 9.42E-90 |
| 31 | 1.76E-130 | 3.57E-105 | 2.23E-79 | 1 | 3.17E-222 | 7.62E-93 |
| 32 | 2.82E-107 | 1.33E-90 | 6.40E-68 | 1 | 8.30E-196 | 7.24E-81 |
| 33 | 1.35E-101 | 6.69E-101 | 9.71E-67 | 1 | 2.21E-207 | 4.00E-91 |
| 34 | 2.64E-101 | 3.37E-99 | 6.09E-80 | 1 | 2.71E-204 | 3.13E-86 |
| 35 | 1.08E-103 | 2.49E-98 | 1.98E-78 | 1 | 2.38E-215 | 1.78E-88 |
| 36 | 3.98E-120 | 6.77E-94 | 4.50E-87 | 1 | 1.96E-207 | 1.54E-81 |
| 37 | 9.63E-118 | 4.56E-103 | 1.28E-71 | 1 | 1.45E-218 | 1.50E-92 |
| 38 | 7.69E-120 | 2.53E-111 | 4.90E-85 | 1 | 6.12E-229 | 5.15E-100 |
| 39 | 2.00E-122 | 6.48E-108 | 5.42E-81 | 1 | 2.78E-221 | 8.61E-98 |

| | | | | | | |
|---|---|---|---|---|---|---|
| 40 | 5.29E-130 | 8.32E-100 | 4.70E-80 | 1 | 4.32E-218 | 4.65E-88 |
| 41 | 3.99E-133 | 1.55E-99 | 2.07E-76 | 1 | 1.37E-208 | 3.01E-86 |
| 42 | 9.77E-104 | 2.15E-95 | 4.53E-78 | 1 | 5.57E-204 | 8.18E-82 |
| 43 | 1.23E-119 | 3.02E-95 | 1.47E-79 | 1 | 5.23E-191 | 1.32E-74 |
| 44 | 1.46E-139 | 4.94E-127 | 4.65E-88 | 1 | 1.02E-241 | 1.17E-110 |
| 45 | 4.82E-113 | 5.89E-94 | 4.18E-74 | 1 | 6.06E-202 | 6.44E-82 |
| 46 | 2.04E-109 | 5.70E-101 | 1.25E-83 | 1 | 1.66E-222 | 6.81E-92 |
| 47 | 2.22E-122 | 4.15E-102 | 1.67E-79 | 1 | 8.53E-218 | 1.27E-93 |
| 48 | 2.60E-103 | 2.16E-93 | 3.55E-76 | 1 | 1.40E-201 | 4.62E-85 |
| 49 | 5.74E-119 | 2.39E-109 | 8.78E-86 | 1 | 5.73E-246 | 1.26E-104 |
| 50 | 3.41E-95 | 2.38E-88 | 5.03E-68 | 1 | 4.72E-202 | 1.96E-79 |
| 51 | 6.14E-118 | 6.04E-96 | 3.49E-77 | 1 | 3.65E-211 | 1.12E-85 |
| 52 | 3.11E-110 | 8.90E-107 | 1.75E-81 | 1 | 8.32E-216 | 2.03E-95 |
| 53 | 2.48E-142 | 8.85E-97 | 6.57E-82 | 1 | 1.11E-190 | 1.33E-80 |
| 54 | 3.55E-95 | 5.80E-92 | 1.63E-74 | 1 | 3.21E-203 | 2.63E-84 |
| 55 | 2.35E-121 | 5.65E-101 | 5.72E-71 | 1 | 4.95E-224 | 3.96E-91 |
| 56 | 1.07E-104 | 3.35E-92 | 5.79E-68 | 1 | 6.82E-201 | 1.96E-83 |
| 57 | 1.20E-131 | 7.60E-92 | 8.01E-74 | 1 | 6.01E-195 | 2.92E-78 |
| 58 | 1.09E-124 | 2.56E-101 | 2.91E-77 | 1 | 3.52E-212 | 1.03E-87 |
| 59 | 1.04E-117 | 9.50E-101 | 8.34E-78 | 1 | 1.02E-218 | 2.71E-92 |
| 60 | 1.75E-104 | 1.26E-100 | 5.00E-73 | 1 | 1.86E-212 | 7.48E-90 |
| 61 | 4.07E-102 | 2.09E-87 | 4.18E-66 | 1 | 3.28E-185 | 5.81E-77 |
| 62 | 5.85E-134 | 9.47E-100 | 1.25E-79 | 1 | 5.05E-205 | 1.75E-86 |
| 63 | 5.15E-135 | 6.44E-94 | 7.94E-74 | 1 | 9.04E-194 | 6.38E-75 |
| 64 | 3.47E-121 | 2.88E-103 | 3.34E-80 | 1 | 1.00E-217 | 1.07E-86 |
| 65 | 1.42E-121 | 1.09E-96 | 2.93E-71 | 1 | 6.77E-210 | 6.82E-89 |
| 66 | 3.70E-104 | 1.68E-92 | 4.51E-80 | 1 | 1.29E-204 | 1.70E-77 |
| 67 | 1.48E-117 | 3.32E-105 | 5.44E-79 | 1 | 7.87E-218 | 6.22E-92 |
| 68 | 3.67E-128 | 8.79E-102 | 1.01E-82 | 1 | 9.69E-221 | 4.84E-84 |
| 69 | 2.15E-119 | 1.58E-99 | 3.72E-71 | 1 | 5.31E-201 | 5.14E-88 |
| 70 | 7.62E-101 | 1.21E-104 | 2.00E-75 | 1 | 1.62E-211 | 3.37E-95 |
| 71 | 4.45E-102 | 5.22E-94 | 4.01E-73 | 1 | 6.62E-201 | 1.01E-83 |
| 72 | 9.63E-91 | 6.94E-88 | 8.85E-66 | 1 | 1.95E-201 | 5.69E-81 |
| 73 | 3.19E-104 | 1.09E-94 | 1.63E-72 | 1 | 2.72E-198 | 2.97E-84 |
| 74 | 1.85E-107 | 1.86E-97 | 7.57E-75 | 1 | 2.37E-203 | 7.46E-85 |
| 75 | 3.38E-108 | 2.16E-92 | 8.66E-77 | 1 | 3.85E-189 | 7.51E-80 |
| 76 | 1.08E-123 | 1.05E-98 | 7.27E-79 | 1 | 1.14E-206 | 1.87E-91 |
| 77 | 4.18E-108 | 5.32E-101 | 9.52E-75 | 1 | 4.21E-223 | 4.01E-96 |
| 78 | 1.41E-125 | 2.74E-111 | 4.34E-84 | 1 | 7.67E-235 | 2.25E-104 |
| 79 | 2.29E-115 | 2.10E-90 | 2.37E-74 | 1 | 3.19E-194 | 1.15E-81 |
| 80 | 5.11E-140 | 2.20E-103 | 3.27E-83 | 1 | 1.90E-214 | 6.36E-89 |
| 81 | 2.30E-113 | 2.30E-98 | 3.66E-73 | 1 | 8.69E-206 | 2.13E-88 |
| 82 | 1.28E-118 | 3.76E-114 | 4.70E-96 | 1 | 2.23E-245 | 3.27E-106 |
| 83 | 1.33E-114 | 8.87E-94 | 4.32E-71 | 1 | 1.51E-190 | 2.70E-78 |
| 84 | 9.32E-109 | 5.19E-115 | 1.48E-74 | 1 | 9.53E-234 | 1.18E-103 |
| 85 | 1.83E-126 | 6.28E-107 | 9.25E-79 | 1 | 2.99E-219 | 7.62E-93 |
| 86 | 9.17E-103 | 5.55E-94 | 3.59E-77 | 1 | 1.12E-203 | 4.36E-82 |
| 87 | 4.90E-112 | 6.35E-92 | 2.08E-79 | 1 | 2.50E-199 | 5.24E-80 |
| 88 | 4.28E-110 | 1.49E-89 | 9.77E-85 | 1 | 9.40E-209 | 3.02E-80 |

| | | | | | | |
|---|---|---|---|---|---|---|
| 89 | 1.21E-125 | 1.79E-98 | 8.58E-85 | 1 | 1.10E-223 | 2.42E-95 |
| 90 | 4.32E-121 | 2.11E-110 | 9.46E-80 | 1 | 7.54E-213 | 1.07E-104 |
| 91 | 8.32E-131 | 3.00E-108 | 5.68E-76 | 1 | 1.06E-215 | 5.08E-91 |
| 92 | 7.66E-134 | 8.30E-103 | 9.07E-83 | 1 | 5.94E-206 | 1.50E-94 |
| 93 | 5.34E-99 | 5.60E-98 | 2.39E-78 | 1 | 1.44E-219 | 1.97E-81 |
| 94 | 5.12E-102 | 3.73E-91 | 1.39E-65 | 1 | 1.83E-196 | 2.05E-85 |
| 95 | 2.81E-117 | 3.44E-103 | 3.89E-81 | 1 | 3.27E-218 | 4.61E-96 |
| 96 | 4.23E-98 | 5.20E-108 | 1.28E-73 | 1 | 1.94E-214 | 1.21E-102 |
| 97 | 6.04E-131 | 2.44E-94 | 1.50E-71 | 1 | 1.29E-191 | 2.22E-85 |
| 98 | 4.62E-116 | 1.68E-100 | 2.86E-70 | 1 | 2.85E-212 | 3.01E-93 |
| 99 | 7.77E-120 | 1.61E-103 | 7.68E-77 | 1 | 2.20E-208 | 5.76E-97 |

**Table S6 – Potential scale reduction factors for BPP analyses.** Mean PSRFs for all MSC parameters based on eight independent chains for each combination of priors. A 0 means the prior formulated with previous data and a 1 means the one increased by two orders of magnitude. Both θ and **τ** had high variation for node C, indicating some convergence issues with that node.

| **Parameter** | **θ=0 τ=0** | **θ=0 τ=1** | **θ=1 τ=0** | **θ=1 τ=1** |
|---|---|---|---|---|
| $\theta_{Ankafobe}$ | 1.13 | 1.10 | 1.37 | 1.60 |
| $\theta_{Ambohitantely}$ | 1.04 | 1.10 | 1.19 | 1.33 |
| $\theta_{Tsinjoarivo}$ | 1.32 | 1.10 | 1.03 | 1.06 |
| $\theta_{Ambatovy}$ | 1.37 | 1.07 | 1.02 | 1.06 |
| $\theta_{Riamalandy}$ | 1.07 | 1.12 | 1.16 | 1.01 |
| $\theta_A$ | 1.01 | 1.01 | 1.03 | 1.01 |
| $\theta_B$ | 1.13 | 1.07 | 1.03 | 1.01 |
| $\theta_C$ | 1.29 | 1.42 | 1.08 | 1.46 |
| $\theta_D$ | 1.37 | 1.72 | 1.06 | 1.04 |
| $\tau_A$ | 1.06 | 1.13 | 1.08 | 1.01 |
| $\tau_B$ | 1.60 | 1.13 | 1.03 | 1.14 |
| $\tau_C$ | 1.28 | 1.13 | 1.35 | 1.24 |
| $\tau_D$ | 1.40 | 1.13 | 1.03 | 1.06 |
| lnL | 1.03 | 1.04 | 1.01 | 1.01 |

**Table S7 – NCBI identifiers and metadata for individuals.**

| Individual | Accession | Site | Lat | Lon | Alt (m) | Collection Date | sex | Collector | SRA | BioSample |
|---|---|---|---|---|---|---|---|---|---|---|
| mleh001 | JMR092 | Ambatovy | -18.800 | 48.350 | 1011 | January 2009 | female | JMR | SRR10878411 | SAMN13832325 |
| mleh002 | MBB001 | Ambatovy | -18.825 | 48.318 | 1107 | February 2015 | female | MBB | SRR10878410 | SAMN13832326 |
| mleh003 | MBB002 | Ambatovy | -18.825 | 48.318 | 1113 | February 2015 | male | MBB | SRR10878399 | SAMN13832327 |
| mleh004 | MBB003 | Ambatovy | -18.825 | 48.318 | 1094 | February 2015 | male | MBB | SRR10878393 | SAMN13832328 |
| mleh005 | RMR95 | Ambohitantely | -18.476 | 47.274 | 1277 | January 2006 | NA | RMR | SRR10878392 | SAMN13832329 |
| mleh006 | RMR96 | Ambohitantely | -18.476 | 47.274 | 1277 | January 2006 | female | RMR | SRR10878391 | SAMN13832330 |
| mleh007 | RMR97 | Ambohitantely | -18.476 | 47.274 | 1277 | January 2006 | male | RMR | SRR10878390 | SAMN13832331 |
| mleh008 | RMR99 | Ambohitantely | -18.476 | 47.274 | 1277 | January 2006 | male | RMR | SRR10878389 | SAMN13832332 |
| mleh009 | MBB036 | Ankafobe | -18.104 | 47.187 | 1478 | September 2015 | female | MBB | SRR10878388 | SAMN13832333 |
| mleh010 | MBB037 | Ankafobe | -18.104 | 47.187 | 1474 | September 2015 | male | MBB | SRR10878387 | SAMN13832334 |
| mleh011 | MBB038 | Ankafobe | -18.104 | 47.187 | 1465 | October 2015 | female | MBB | SRR10878409 | SAMN13832335 |
| mleh012 | MBB039 | Ankafobe | -18.104 | 47.187 | 1463 | October 2015 | female | MBB | SRR10878408 | SAMN13832336 |
| mleh013 | MBB040 | Ankafobe | -18.103 | 47.187 | 1435 | February 2016 | male | MBB | SRR10878407 | SAMN13832337 |
| mleh014 | MBB041 | Ankafobe | -18.117 | 47.195 | 1444 | February 2016 | male | MBB | SRR10878406 | SAMN13832338 |
| mleh015 | MBB042 | Ankafobe | -18.103 | 47.187 | 1435 | February 2016 | male | MBB | SRR10878405 | SAMN13832339 |
| mleh016 | MBB043 | Ankafobe | -18.106 | 47.187 | 1441 | February 2016 | female | MBB | SRR10878404 | SAMN13832340 |
| mleh017 | MBB044 | Ankafobe | -18.106 | 47.187 | 1464 | February 2016 | female | MBB | SRR10878403 | SAMN13832341 |
| mleh018 | MBB045 | Ankafobe | -18.119 | 47.193 | 1426 | February 2016 | female | MBB | SRR10878402 | SAMN13832342 |
| mleh019 | JMR001 | Riamalandy | -16.285 | 48.815 | 823 | November 2004 | Female | JMR | SRR10878401 | SAMN13832343 |
| mleh020 | JMR002 | Riamalandy | -16.285 | 48.815 | 823 | November 2004 | Male | JMR | SRR10878400 | SAMN13832344 |
| mleh021 | DWW3235 | Tsinjoarivo | -19.721 | 47.857 | 1396 | December 2007 | NA | MBB | SRR10878398 | SAMN13832345 |
| mleh022 | DWW3236 | Tsinjoarivo | -19.721 | 47.857 | 1396 | December 2007 | NA | MBB | SRR10878397 | SAMN13832346 |
| mleh023 | DWW3243 | Tsinjoarivo | -19.721 | 47.857 | 1396 | December 2007 | NA | MBB | SRR10878396 | SAMN13832347 |
| mleh024 | DWW3244 | Tsinjoarivo | -19.721 | 47.857 | 1396 | December 2007 | NA | MBB | SRR10878395 | SAMN13832348 |
| mleh025 | DWW3249 | Tsinjoarivo | -19.721 | 47.857 | 1396 | December 2007 | NA | MBB | SRR10878394 | SAMN13832349 |
